# Local genetic adaptation to habitat in wild chimpanzees

**DOI:** 10.1101/2024.07.09.601734

**Authors:** Harrison J. Ostridge, Claudia Fontsere, Esther Lizano, Daniela C. Soto, Joshua M. Schmidt, Vrishti Saxena, Marina Alvarez-Estape, Christopher D. Barratt, Paolo Gratton, Gaëlle Bocksberger, Jack D. Lester, Paula Dieguez, Anthony Agbor, Samuel Angedakin, Alfred Kwabena Assumang, Emma Bailey, Donatienne Barubiyo, Mattia Bessone, Gregory Brazzola, Rebecca Chancellor, Heather Cohen, Charlotte Coupland, Emmanuel Danquah, Tobias Deschner, Laia Dotras, Jef Dupain, Villard Ebot Egbe, Anne-Céline Granjon, Josephine Head, Daniela Hedwig, Veerle Hermans, R. Adriana Hernandez-Aguilar, Kathryn J. Jeffery, Sorrel Jones, Jessica Junker, Parag Kadam, Michael Kaiser, Ammie K. Kalan, Mbangi Kambere, Ivonne Kienast, Deo Kujirakwinja, Kevin E. Langergraber, Juan Lapuente, Bradley Larson, Anne Laudisoit, Kevin C. Lee, Manuel Llana, Giovanna Maretti, Rumen Martín, Amelia Meier, David Morgan, Emily Neil, Sonia Nicholl, Stuart Nixon, Emmanuelle Normand, Christopher Orbell, Lucy Jayne Ormsby, Robinson Orume, Liliana Pacheco, Jodie Preece, Sebastien Regnaut, Martha M. Robbins, Aaron Rundus, Crickette Sanz, Lilah Sciaky, Volker Sommer, Fiona A. Stewart, Nikki Tagg, Luc Roscelin Tédonzong, Joost van Schijndel, Elleni Vendras, Erin G. Wessling, Jacob Willie, Roman M. Wittig, Yisa Ginath Yuh, Kyle Yurkiw, Linda Vigilant, Alex Piel, Christophe Boesch, Hjalmar S. Kühl, Megan Y. Dennis, Tomas Marques-Bonet, Mimi Arandjelovic, Aida M. Andrés

## Abstract

How populations adapt to their environment is a fundamental question in biology. Yet we know surprisingly little about this process, especially for endangered species such as non-human great apes. Chimpanzees, our closest living relatives, are particularly interesting because they inhabit diverse habitats, from rainforest to woodland-savannah. Whether genetic adaptation facilitates such habitat diversity remains unknown, despite having wide implications for evolutionary biology and conservation. Using 828 newly generated exomes from wild chimpanzees, we find evidence of fine-scale genetic adaptation to habitat. Notably, adaptation to malaria in forest chimpanzees is mediated by the same genes underlying adaptation to malaria in humans. This work demonstrates the power of non-invasive samples to reveal genetic adaptations in endangered populations and highlights the importance of adaptive genetic diversity for chimpanzees.

**One-Sentence Summary:** Chimpanzees show evidence of local genetic adaptation to habitat, particularly to pathogens, such as malaria, in forests.

## Main Text

Understanding how primates are adapted to their environments provides insights into our own evolution and vital information for conservation efforts. This is particularly relevant and urgent for our closest living relatives, non-human great apes, all of which are either endangered or critically endangered. Chimpanzees (*Pan troglodytes*) have the largest geographic and ecological range of any non-human ape (2.6 million km² (*1*)) spanning a variety of environments across Equatorial Africa, from dense tropical rainforest to open woodland-savannah mosaics (hereafter ‘savannah’ for simplicity (*2*)). Aside from humans, they are the only great apes that inhabit savannah habitats (*2*). Yet, each of the four subspecies of chimpanzee (central (*P. t. troglodytes*), eastern (*P. t. schweinfurthii*), Nigeria-Cameroon (*P. t. ellioti*) and western (*P. t. verus*) (*3–5*)) are endangered (westerns critically so) with numbers continuing to decline due to hunting, habitat destruction and infectious diseases (*1*, *6–8*). This decline has widespread negative impacts, as chimpanzees are important conservation flagship species for biodiversity protection and crucial ecosystem engineers (*9–11*).

Between the forest and savannah extremes, chimpanzees occupy a gradient of habitats known as forest-savannah mosaics (*12*). Forests, which are likely closest to chimpanzee ancestral habitats (*3*, *13*), have closed canopies with high availability of food and water throughout the year, and therefore tend to support high population densities (*2*). Forests also harbour a great diversity of pathogens and disease vectors (*14*). Conversely, savannahs are on the edge of chimpanzee distribution in East and West Africa and are characterised by open canopies, higher temperatures, lower annual rainfall and higher rainfall seasonality (*2*, *15*).

The occupation of such a range of habitats is facilitated by chimpanzee behavioural diversity (*16*). Savannah chimpanzees exhibit unique thermoregulatory behaviours (*17*, *18*) and on average trend towards greater behavioural diversity than forest chimpanzees (*16*)–a potential adaptation to higher environmental variability. Behavioural adaptations also include tool use in a range of contexts such as foraging (*19–21*), water extraction (*22–24*) and communication (*25*). Nevertheless, behaviour does not fully compensate for stressors, as shown by physiological stress in response to pathogens (*26–29*) and environmental pressures (*15*, *30*). Another mechanism that can facilitate the occupation of diverse habitats is genetic adaptation, just as local adaptation has contributed to genetic population differentiation in humans (*31*) despite great behavioural flexibility (*32*, *33*). In fact, humans have evolved local genetic adaptations to environmental pressures that differ between forest and savannah habitats, including pathogens (*34–36*) such as malaria (*37*, *38*), and climatic variables such as temperature and water availability (*39*, *40*), diet (*41–43*), and solar exposure (*44*). Culture can also promote genetic adaptations, such as in human adaptations to diet and zoonotic diseases associated with animal domestication (*43*, *45*).

Establishing if genetic differences underlie local adaptation in chimpanzees is important to understanding primate evolution and critical for chimpanzee conservation. If adaptive genetic differences exist among populations, this genetic diversity must be conserved to maintain existing adaptations and adaptive potential (*46*, *47*). Additionally, recent genetic adaptations highlight key selective pressures that likely shape fitness in the wild today, and can help establish which populations may be more vulnerable to environmental change (*46*). This is particularly relevant in the face of anthropogenic climate change, which is increasing temperatures and precipitation seasonality within the chimpanzees’ range (*2*). Furthermore, chimpanzees are excellent models for understanding our own evolution, particularly in savannah regions, which resemble early hominin habitats (*2*, *48–52*). Lastly, the close genetic similarity between humans and other great apes (*53*) has resulted in zoonotic disease transmissions (*54*, *55*) such as HIV-1/AIDS (*56*) and malaria (*57*). Understanding how chimpanzees have evolved to reduce the pathogenicity of microorganisms can thus reveal potential targets for treatments and vaccines (*58–61*).

We have a growing understanding of chimpanzee demographic history thanks to population genomics studies (*3*, *5*, *62*), which have identified genetic differentiation among populations within each subspecies. However, our knowledge of genetic adaptation lags behind, largely due to sample limitations. Because existing genomic datasets include only dozens of captive chimpanzees of unknown geographic origin (*5*, *62–64*), previous studies investigated adaptation only at the subspecies level (*63*, *65–73*), revealing interesting subspecies-level adaptations e.g., to pathogens such as SIV (*65*, *66*). However, habitats vary greatly within chimpanzee subspecies ranges (*12*); therefore, subspecies comparisons are uninformative on adaptations to many potential selective pressures. Investigation of fine-scale local adaptation is essential for understanding adaptation in chimpanzees but requires large numbers of DNA samples from wild individuals of known geographic origin, coupled with detailed environmental data.

Non-invasive sampling is the only ethical and feasible option for studying wild populations of many protected species (*74*), including non-human apes; however, recent technical and analytical advancements are beginning to enable population genomic analyses on such samples (*3*, *75–80*). Using faecal samples from wild individuals collected as part of the Pan African Programme: The Cultured Chimpanzee (PanAf) (*3*, *81*), we have generated full exome (protein-coding regions of the genome) sequences from hundreds of chimpanzees across their geographic and environmental range. We demonstrate that genomic data from non-invasive samples can be used to reveal the fine-scale adaptive history of endangered primates. Specifically, when integrated with environmental data, the exomes reveal evidence of local genetic adaptation to habitat conditions in chimpanzees. In forests, we find evidence of pathogen-mediated adaptation, including to malaria via the same loci that mediate malaria adaptation in humans.

### Samples and sequences

Faecal samples of 828 unique individuals were collected from 52 sampling sites across the geographic range of all four chimpanzee subspecies as part of PanAf (*3*, *81*). This represents a ten-fold increase in sample size and a massive increase in geographic coverage over existing genome-wide datasets (*5*, *62*) of any non-human ape. The scale and resolution of the dataset are only comparable to the chromosome 21 (chr21) sequences of the same individuals (*3*).

Non-invasive samples typically contain low levels of endogenous DNA. Thus, we target-captured and sequenced full exomes (akin to chr21 (*3*)) because they are informative for the vast majority of functional sites in the genome, including both sequenced protein-coding and linked regulatory regions (e.g., promoters). Samples were strictly filtered to omit those with first-order relatives, contamination or low read depth (Supplemental Note 3). To mitigate the potential effects of the moderate read depth and take advantage of the large sample size, we used genotype likelihoods and allele frequency-based methods, which minimised the effects of individual sequencing errors.

As expected, using either exomes or chr21 (*3*), population structure analyses separate samples into four subspecies (Figs. S12, S14 and S15), and within-subspecies population structure inferred with the exomes closely matches results from chr21 (*3*) (Figs. S12-16). Each sample site was considered a genetic unit, which we refer to as a ‘population’, except for four populations formed by combining very closely related sample sites (details in Materials and Methods and Supplemental Note 4). After removing populations with fewer than eight samples, the final dataset contains 388 exomes (385 chr21) from 30 populations: 5 central, 9 eastern, 2 Nigeria-Cameroon and 14 western. The resulting exomes have a median read depth per sample of 5.30-fold (0.51- to 52.27-fold) in the exome target space (60Mbp). The signatures of local adaptation within and across subspecies were investigated in four ‘subspecies-datasets’ containing populations from all subspecies (All), central and eastern together (Central-Eastern), Nigeria-Cameroon (Nigeria-Cameroon) and western (Western). Central and eastern were combined because of their low genetic differentiation (F_ST_=0.10, Fig. 1) (*65*). The unfolded site frequency spectra (SFS) conform to expectations given the inferred demographic history of these subspecies (Fig. S18).

**Fig. 1.**
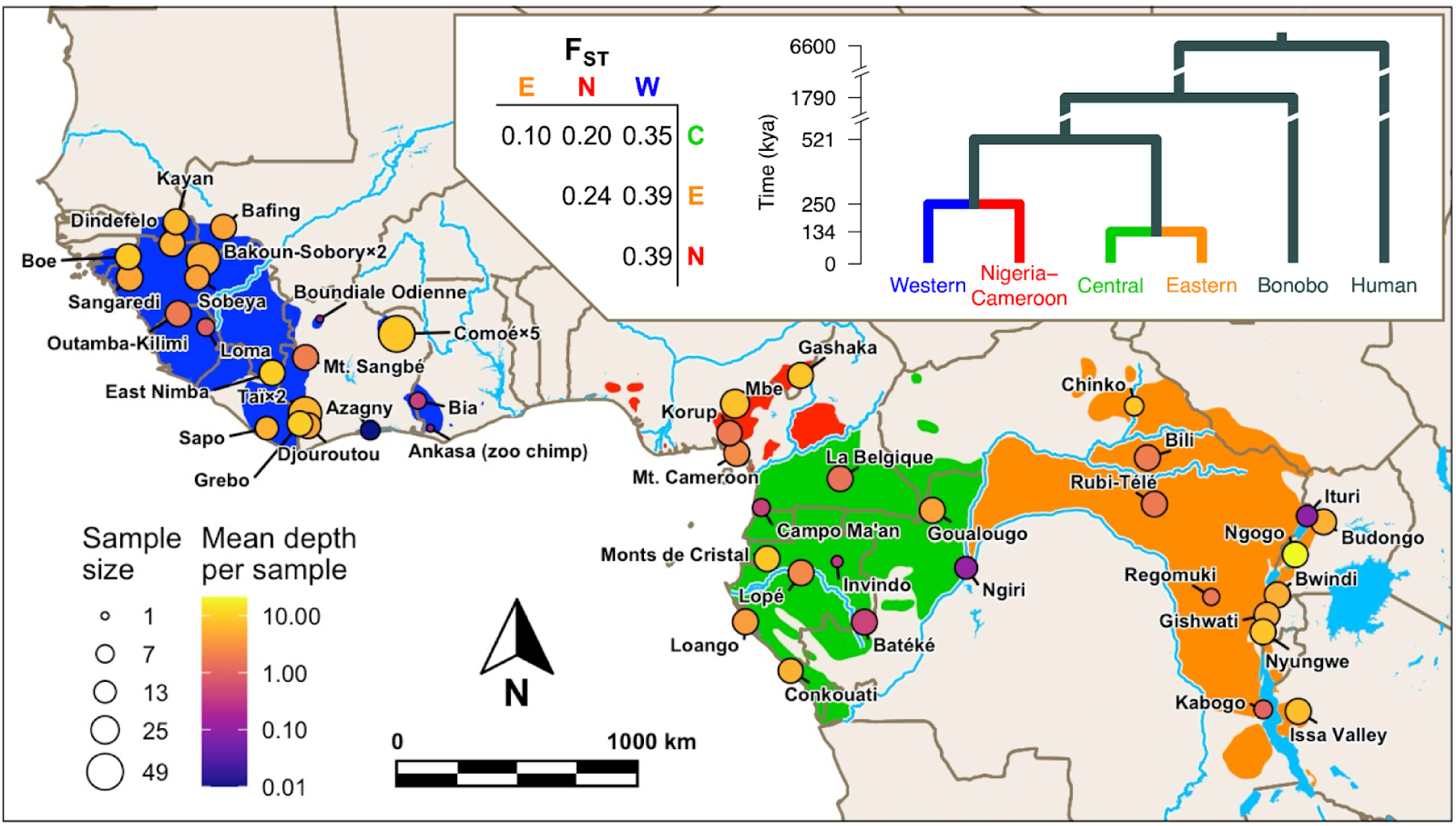
Chimpanzee exome dataset distribution, sample size and coverage. Top panel: Hominini phylogenetic tree highlighting chimpanzee subspecies with estimated evolutionary split times (*62*, *82*) (thousands of years ago (kya)) and pairwise F_ST_ between subspecies (*65*). Main panel: map of West, Central and East Africa indicating the location of sample sites, sample sizes and mean exome sequencing read depth per sample. Each point represents a sample site except for five geographically close sites sampled at Comoé, two at Taï and two at Bakoun-Sobory. The geographic distribution of each subspecies is shown (green=central, orange=eastern, red=Nigeria-Cameroon, blue=western) (*1*) with major rivers and lakes indicated in light blue. The sample sizes and populations in the final filtered dataset used for selection analyses are shown in Fig. 3.

### Allele frequency population differentiation

Local adaptation increases the frequency of alleles only where they are beneficial, generating large allele frequency differences among populations. We first investigated local positive selection by analysing population differentiation with a genetics-only hypothesis-free analysis using the BayPass (*83*) core model. BayPass estimates the genome-wide population allele frequency covariance matrix, which is used to standardise allele frequencies for each single-nucleotide polymorphism (SNP) with respect to population structure (BayPass effectively accounts for population structure here, see Supplemental Note 4); the variance across populations of these standardised frequencies is summarised in the test statistic X^t^X* (*84*). SNPs under local adaptation are expected to have exceptionally large population allele frequency differentiation, and therefore the highest X^t^X* values in the genome. Null expectations under neutrality were generated using the non-genic regions of chr21 (non-genic-chr21) in these samples (*3*). Generating an empirical null frees the analysis from demographic assumptions and accounts for many potential confounding factors because non-genic-chr21 has an almost identical demographic history, sample size and read depth to the exome and has been processed in the exact same way (Supplemental Note 6.2). Candidate targets of positive selection (hereafter ‘candidate SNPs’) were defined as the exome SNPs with higher X^t^X* than the values corresponding to estimated false positive rates (FPR) of 0.5%, 0.1% and 0.05% using the non-genic-chr21 X^t^X* distribution while accounting for read depth (details in Materials and Methods and Supplementary Note 6.3.2).

If local adaptation drives population differentiation, we expect exomes to show an excess of highly differentiated SNPs compared to neutral expectations. Contrary to this expectation, there are fewer SNPs with very large X^t^X* values in the exome compared to null expectations (Fig. 2, Fig. S27), potentially reflecting the effects of purifying selection in the exome. It remains possible that specific selection pressures have driven local genetic adaptation in chimpanzees but we do not observe evidence of this at the genome level using exomes alone.

**Fig. 2.**
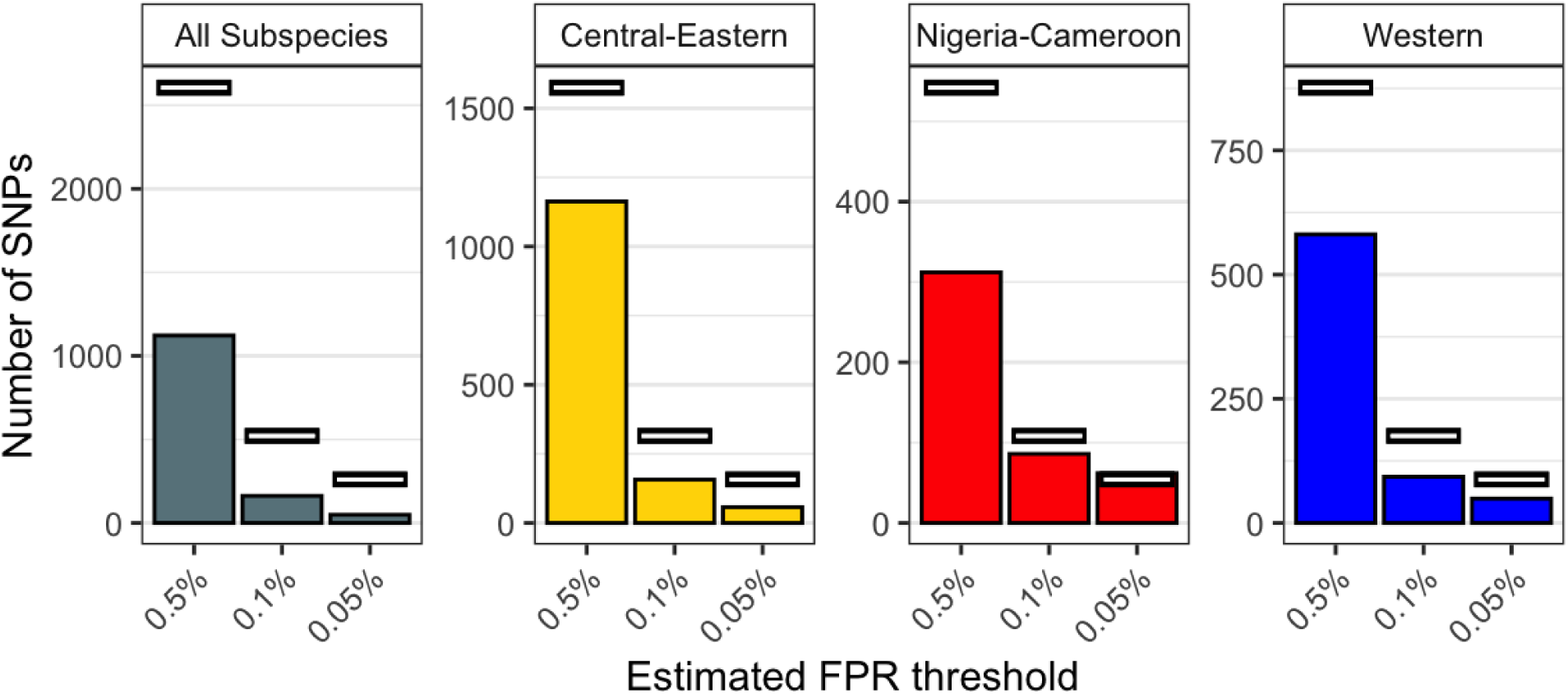
Number of genetics-only candidate SNPs. The number of candidate SNPs from the genetics-only test (bars) compared to the null expectation (white lines) at X^t^X* thresholds corresponding to estimated FPRs of 0.5%, 0.1% and 0.05%, for each subspecies-dataset. Note that y-axis scales are not consistent across panels.

### Genetic adaptation to habitat

Integrating genetic and environmental data increases the power to detect signatures of local adaptation (*85*, *86*) and allows us to directly test the hypothesis that chimpanzees have adapted to selection pressures that vary between habitats. We thus performed a genotype-environment association (GEA) test by integrating an environmental covariable into the analysis using the BayPass AUX model (*83*). BayPass calculates a Bayes factor (BF) for each SNP that indicates the strength of evidence for the linear correlation between population allele frequencies and the environmental covariable while accounting for population structure (Supplemental Note 6.4.5). SNPs evolving under local adaptation are expected to be highly correlated with the relevant environmental covariable and therefore have the highest BF in the genome.

Environmental metrics based on temperature, precipitation or land cover do not correspond well with researcher-defined forest and savannah regions (*12*). Therefore, we used a floristic measure informed by a large-scale biogeographic analysis which identified very different tree species compositions between forest and savannah regions and has been shown to produce more accurate maps of habitat distributions across Africa (*87*). Specifically, we used the percentage of trees identified as ‘forest specialists’ (*87*) among all the classified trees recorded at each sample site (hereafter ‘forest-tree-percentage’) (Fig. 3) (see Materials and Methods and Supplemental Note 2 for details). This variable is ideal because the data was collected within the known ranges of the sampled populations and the same field protocol can be applied to new sample sites, making our data and results comparable to future studies incorporating additional sample sites. This variable is not used here to test for adaptation to tree species compositions *per se*; rather, it is used to describe the chimpanzee habitat gradient, which summarises many potential selective pressures. The GEA analysis was run with this covariable in each subspecies-dataset except *Nigeria-Cameroon*, as it has only two populations. Candidate SNPs were selected as in the genetics-only test (details in Supplementary Note 6.4.1).

**Fig. 3.**
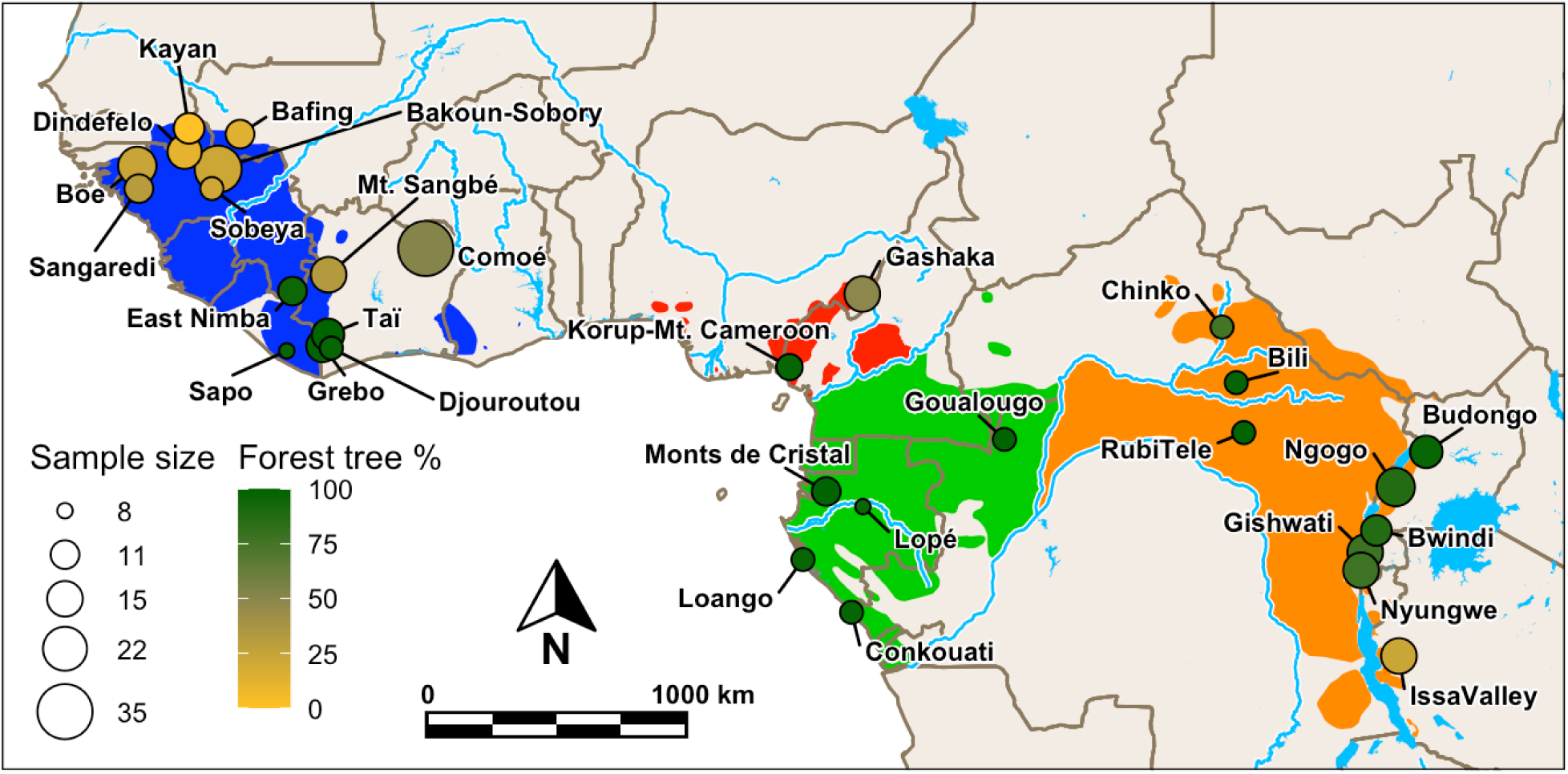
BayPass analysis dataset. Map of West, Central and East Africa showing the location, sample size (after filtering) and forest-tree-percentage for each population in the BayPass analyses. The ranges of the four subspecies are shown (green=central, orange=eastern, red=Nigeria-Cameroon, blue=western) (*1*) with major rivers and lakes indicated in light blue.

In contrast to the genetics-only results, the GEA shows a substantial excess of SNPs strongly associated with forest-tree-percentage in the exome when compared with neutral expectations in *All* and *Central-Eastern* (Fig. 4). This excess is what we expect under local adaptation associated with habitat. This excess is not present in *Western,* as expected if the strong population bottleneck experienced by this subspecies (*3*, *5*, *62*) increased drift and reduced the efficacy of natural selection, or if recent gene flow (*3*) inhibited the evolution of local adaptation (Fig. 4). Nevertheless, this does not exclude the possibility that strong selective forces may have driven local adaptation in the western subspecies, and the SNPs with the highest BFs in the exome are the best candidate targets of positive selection. Read depth and population substructure do not drive our candidates, and independent tests confirm that candidate allele frequencies correlate strongly with forest-tree-percentage (Figs. S39 and S44-46). For all thresholds and subspecies-datasets, the minimum BF is very high: over 14.7 for FPR<0.5%, over 18.3 for FPR<0.1% and over 19.5 for FPR<0.05% (Fig. S38). Jeffrey’s rule (*88*) defines 15<BF<20 as ‘very strong evidence’ and BF>20 as ‘decisive evidence’, demonstrating that a vast majority of candidate SNPs have very strong evidence of being associated with habitat, with almost all SNPs in the 0.1% tail having decisive evidence (Fig. S37).

**Fig. 4.**
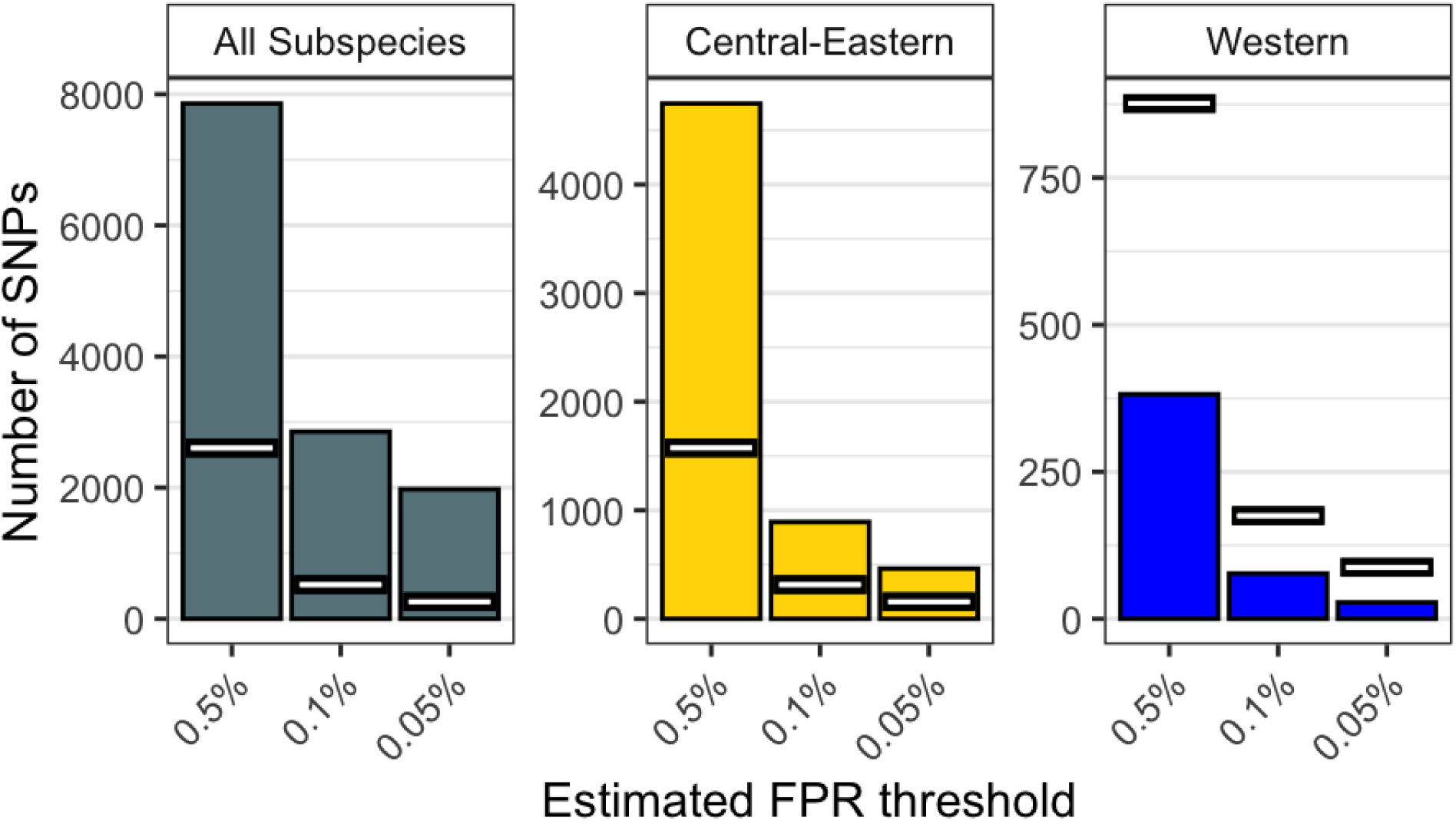
Number of GEA candidate SNPs. The number of candidate SNPs from the GEA (bars) compared to the null expectation (white lines) at BF thresholds corresponding to estimated FPRs of 0.5%, 0.1% and 0.05%, for each subspecies-dataset tested. Note that y-axis scales are not consistent across panels.

These results provide strong evidence for local genetic adaptation to habitat in chimpanzees, revealing the presence of important genetic differences among wild populations, even within subspecies, that likely shape fitness in an environment-dependent way.

Under the reasonable assumption that novel adaptations are more commonly mediated by the novel, derived allele than the ancestral one, we assigned SNPs as associated with forest or savannah adaptations according to the sign of their correlation coefficient. There is an excess of SNPs with high BFs in the exome for both savannah and forest candidates (Fig. S42) in *All* and *Central-Eastern,* suggesting that adaptation in either direction contributes to the overall excess. To interpret these loci biologically, we investigated the genes the candidate SNPs fall within (hereafter ‘candidate genes’) by testing for an overrepresentation of functional categories in hypothesis-free gene set enrichment analyses. Given the relevance of pathogens as selective pressures (*65*, *66*), we also performed a hypothesis-driven enrichment analysis of pathogen-related genes (details in Supplemental Note 7). Interestingly, these analyses point to potential differential adaptations in savannah and forest chimpanzees.

### Adaptations to savannah

Savannah candidate genes belong to many categories associated with physiological traits when compared to forest candidate genes (Fig. 5A). There are over six times more “General” gene categories with an FDR<0.5 in savannah than forest candidates (260 vs 42, Figs. 5A and S50), although only two of these categories are significantly enriched (FDR < 0.05) (Fig. 5B) (negative regulation of nitrogen compound metabolic processes and negative regulation of cellular macromolecule biosynthetic processes). This would be compatible with a large degree of polygenicity in the genetic adaptation of chimpanzees to the environmental extremes of savannahs. The diversity of categories and their overlap in genes makes it difficult to infer selection pressures that may drive this signal.

**Fig. 5.**
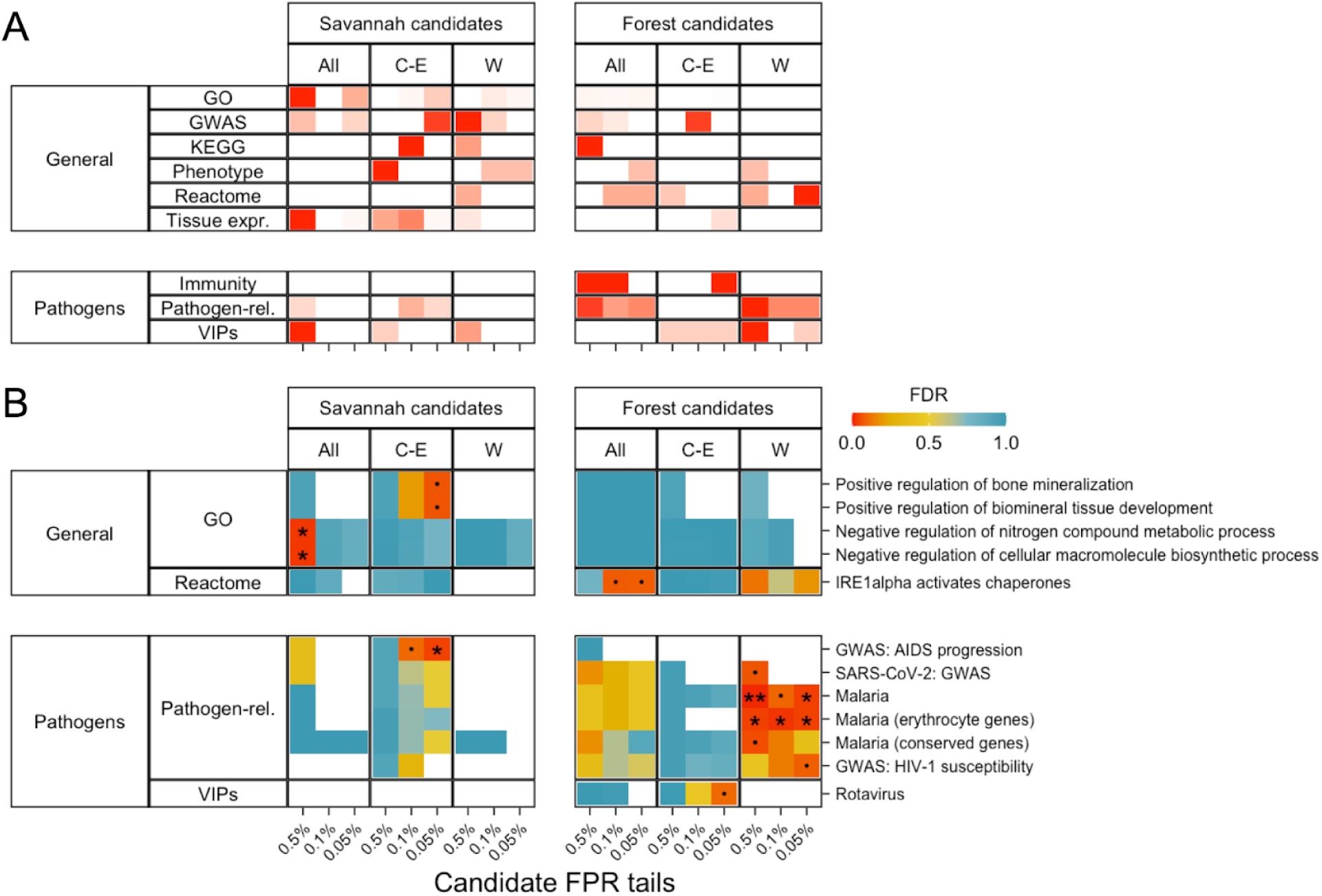
GEA candidate gene set enrichment results. Results for 0.5%, 0.1% and 0.05% FPR tails for savannah and forest candidate SNPs are shown. Vertical panels indicate results from each subspecies-dataset. Horizontal panels show the broad categories that the gene sets belong to. Multiple testing correction was done within each gene set enrichment analysis run (i.e., each tail and gene set database such as ‘Pathogen-related’, ‘GWAS’, ‘Phenotype’ etc.). (**A**) the number of gene sets with FDR<0.5, cells are coloured in a gradient from white (0) to red (the largest value per row) (Fig. S50 shows the numbers in each cell). (**B**) shows the FDR values for the most enriched gene sets with FDR<0.1 for at least one candidate tail in at least one subspecies-dataset (‘.’ FDR<0.1, ‘*’ FDR<0.05, ‘**’ FDR<0.01).

Restricted availability of water in savannahs during the dry season is a potential selection pressure (*17*, *30*) that could partially explain the enrichment of physiological categories in our candidates. Nevertheless, there is no significant enrichment in the two dehydration response gene categories we analysed (*89*, *90*) (Supplemental Note 7), neither in the GEA nor genetics-only candidates (Fig. S51). Chimpanzees may thus have adapted to dehydration stress through genes not included in these categories or through behavioural adaptations (e.g., well digging (*22*)). Alternatively, dehydration stress may be present but independent of habitat (*15*).

There is limited evidence of adaptation to pathogens in savannah populations (Fig. 5A, left bottom corner). Savannah candidates are significantly enriched (FDR < 0.05) for only one pathogen-related gene set: genes associated with AIDS progression in GWAS at the *Central-Eastern* 0.05% tail (Fig. 5B), which contains only two candidate genes (Table S2). Viruses similar to SIV/HIV are not known to be associated with savannahs, instead, this result may be explained by adaptation in Issa Valley, which has a high prevalence of SIV (*91*, *92*) and a particularly low forest-tree-percentage in *Central-Eastern* (Fig. 3). Being extreme in forest-tree-percentage means that Issa Valley weighs heavily on the *Central-Eastern* savannah candidates, although without fully driving them (Supplemental Note: 6.4.6). Analysis of additional populations will help establish to what extent the evidence of adaptation is general across central and eastern savannah populations, and identify the specific adaptive mechanisms and selective factors. In any case, the excess of exonic savannah candidate SNPs in *All* and *Central-Eastern* suggests that chimpanzees do harbour genetic adaptations to savannah habitats.

### Adaptations to forest

While forest candidates show weak enrichment in general physiological categories, they show a pattern of stronger enrichment in pathogen-related genes as shown in stronger enrichment for general “immunity genes” (*93*) and “innate immunity genes” (*94*) in the forest than savannah candidates in *All* and *Central-Eastern* (Fig. 5A). This pattern is also evident for individual pathogen categories in *All* and especially in *Western*, although not in *Central-Eastern* (Fig. 5). This is consistent with the higher population densities (*2*) and increased pathogen exposure (*14*) in forests resulting in a greater infectious disease burden. In humans, local adaptation has likely also been driven by high pathogen diversity (*95*), particularly in tropical forests (*34–36*). *Central-Eastern* does not show this pattern, likely due to the presence of eastern populations from montane forests, which are considerably cooler than lowland forests and therefore have lower levels of vector-borne diseases such as malaria (*14*) (Supplemental Note 7.2.1). Enrichment of pathogen-related categories in the *Western* forest candidates suggests that, although we do not see evidence of positive selection on the genome-scale, strong selection at a limited number of pathogen-related genes is likely driving local adaptation in this subspecies.

Focusing on individual pathogens, the strongest and clearest signal is enrichment for malaria-related categories in *Western* forest candidates (Fig. 5B, Table S2). They are significantly enriched (FDR<0.05) in “Malaria related genes” (*96*) at the 0.5% and 0.05% tails and “Erythrocyte genes related to malaria” (*97*) at all three tails; enrichment in *Plasmodium*-interacting proteins that are conserved across mammals (*98*) (thus excluding haemoglobin and glycophorin genes, see below) very narrowly exceeds the significance threshold (FDR=0.050) in the 0.5% tail. Interestingly, malaria infection probability in chimpanzees is closely correlated with canopy cover (*14*), which is itself highly correlated with forest-tree-percentage in our dataset (Pearson r = 0.92, p = 9.044×10^-13^) (*99*). Malaria is a major selection pressure and has driven some of the clearest examples of local adaptation in humans (*37*, *38*). Only five genetic variants have been significantly associated with severe malaria in human GWAS (*100*, *101*). Strikingly, two of these loci, which encode for haemoglobin (HB) and glycophorin (GYP) genes (Fig. 6A), contain chimpanzee forest candidate SNPs; both of which also underlie adaptations to malaria in humans (*102–105*).

**Fig. 6.**
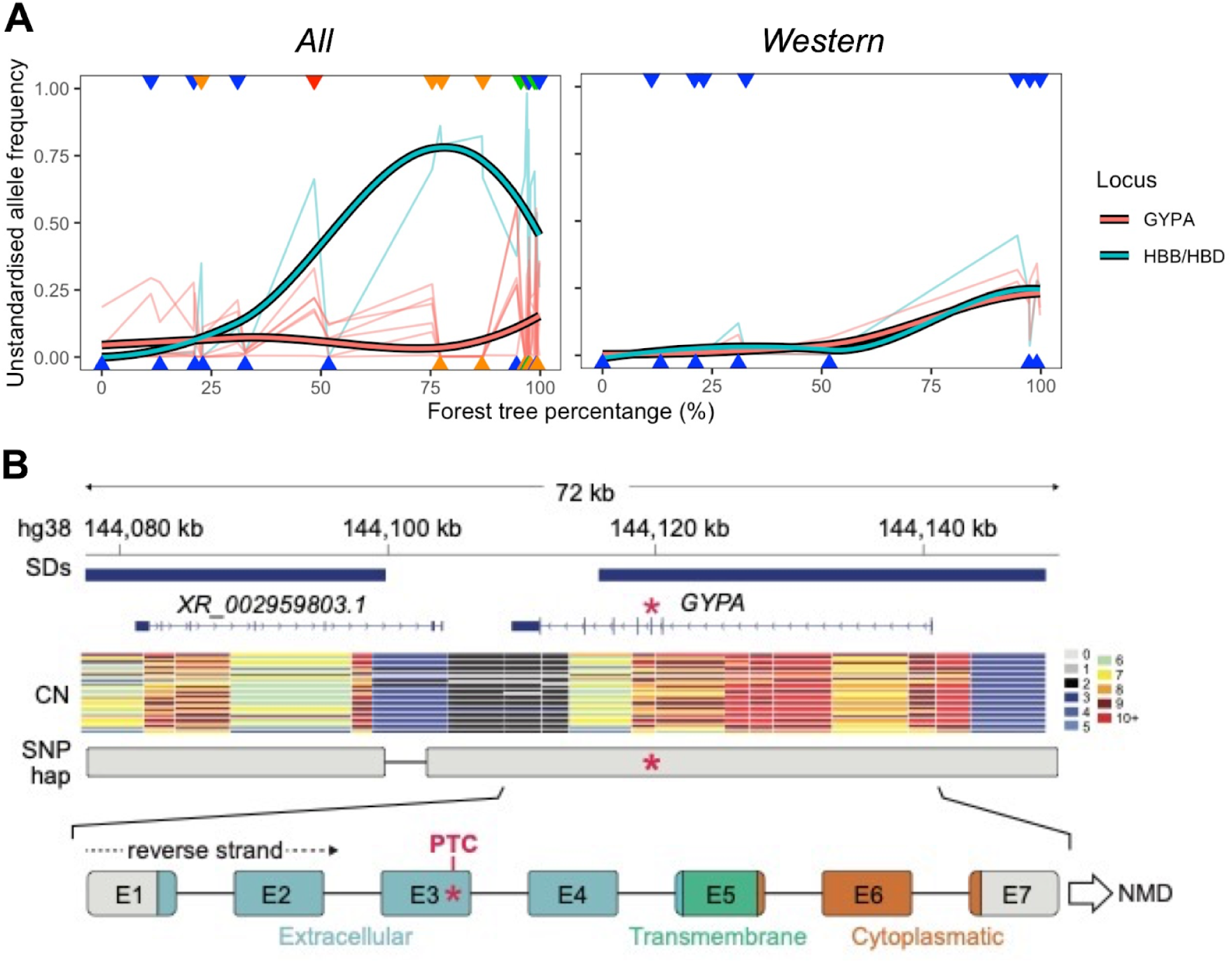
Key malaria-related forest candidate genes. (**A**) Derived allele frequencies of candidate SNPs at the *HBB*/*HBD* (green) and *GYPA* (red) loci plotted against forest-tree-percentage, with population values indicated with triangles coloured according to subspecies (green=central, orange=eastern, red=Nigeria-Cameroon, blue=western) arbitrarily assigned to the top or bottom of the graph to reduce overlap. Left for candidate SNPs and populations from *All*. Right for candidate SNPs and populations from *Western*. Thin lines represent the estimated population allele frequencies for each candidate SNP, thick lines show the smoothed pattern of all candidate SNPs per locus using LOESS. (**B**) Diagram of the *GYPA* locus in hg38 coordinates, including segmental duplications (SDs), copy numbers (CN) across captive chimpanzees, representative long-read sequencing haplotype containing candidate SNP C>A at chr4:145040845 in hg19 coordinates (red asterisk), and schematic representation of the candidate SNP location within *GYPA* exons (E1-7). PTC: premature termination codon. NMD: Nonsense-mediated mRNA decay. The PTC SNP is 210 exonic base pairs upstream of the last exon-exon junction (between exons 6 and 7), and therefore, likely to cause NMD according to the 50–55nt rule (*116*).

For HB, candidate SNP chr11:5254366 (*Western* 0.5% and *All* 0.05% tails) lies within an intron of Haemoglobin Subunit Delta (*HBD*), less than 5kb upstream of the adjacent paralogue Haemoglobin Subunit Beta (*HBB*). While mutations in *HBD* have little effect on malaria resistance due to low expression in adults (*106*), the HbS mutation in *HBB* is a classic example of balancing selection in humans, as heterozygotes are protected against severe malaria (*102*, *103*). Therefore, the signatures that we observe may reflect selection on a linked variant within *HBB*, the regulation of *HBB* or *HBD* itself. In any case, it is striking that this locus shows evidence of local adaptation in both chimpanzees and humans.

For GYP, two candidate SNPs, chr4:145040845 and chr4:145039806 (*Western* 0.5% and 0.05% tails respectively) lie within Glycophorin A (*GYPA*) (Ensemble hg19 also places them within an intron of Glycophorin B (*GYPB*) likely due to an annotation error, see Supplemental Note 8.2) (Fig. S52). The evidence for selection at this locus is strong with chr4:145039806 having the 23^rd^ highest BF in *Western* accounting for read depth (FPR≤2.91×10^-4^). In *All*, there are six forest candidate SNPs in *GYPA* including chr4:145040845 (0.05% tail) and chr4:145039806 (0.5% tail), and another *GYPA* SNP is a candidate in the genetics-only 0.5% tail (Fig. S52).

*GYPA* and *GYPB* encode glycophorins used by *Plasmodium falciparum* to enter erythrocytes (*107*). In humans, structural variants associated with this locus appear to mediate adaptation to malaria (*104*, *105*, *108–112*). It is therefore interesting to investigate structural variation in chimpanzees. Read depth in the PanAf exomes is not unusual at this locus, but low-coverage target capture data is not ideal for investigating structural variation. Copy-number (CN) estimates from high-coverage short-read (n=60) (*5*, *62*) and long-read data (n=2) (*113*, *114*) from captive chimpanzees confirm that, in addition to the full-length and likely ancestral *GYPA*, chimpanzees also carry 2–9 copies of truncated paralogues lacking the last two exons of *GYPA*, which encode for the cytoplasmic domain (*115*) (Supplemental Note 8.1, Fig. 6B, Fig. S53D). Thus, like in humans, structural variants contribute to the complexity of the locus in chimpanzees. We note that the *GYPA* candidate SNPs are present in both the long-read (*113*, *114*) and high-coverage short-read (*5*, *62*) data (Fig. S55), confirming them to be true polymorphisms; further, there is no evidence of an association between forest-tree-percentage and read-depth in the PanAf exomes at these SNPs (Fig. S54, details in Supplemental Note 8.1).

The long-read data (*113*, *114*) show the *GYPA* candidate SNPs residing in a single haplotype spanning the full-length gene (Fig. 6B, Fig. S53C). The candidate allele at chr4:145040845 introduces a premature stop gain in exon 3 of *GYPA* (E76X) predicted to result in degradation of the mRNA by nonsense-mediated decay (*116*); even if the truncated protein was translated, it would encode only a partial extracellular domain and be missing the remaining extracellular and entire transmembrane and cytoplasmic domains (*115*), resulting in non-functional GYPA. Thus, as suggested for *GYPA* deletions in humans (*109*), this SNP may be the target of natural selection, by preventing the expression of a key receptor protein used by the malaria parasite to enter erythrocytes (*107*).

## Conclusions

We present the largest population genomic study of natural selection in a non-human ape to date, capturing and sequencing the exome from non-invasive samples of hundreds of wild chimpanzees and integrating these exomes with previously published full chr21 sequences from the same samples (*3*). Even in the face of limitations from non-invasive sampling (Supplemental Note 1), this work demonstrates that population genomics can reveal the presence of local genetic adaptation in an endangered species.

The genotype-environment association analysis provides strong, genome-scale evidence of local adaptation to habitat in *All* and *Central-Eastern–*although not in *Western*, likely because of their small long-term *N_e_* (*5, 62*). This demonstrates the power of genotype-environment associations (GEA) to identify positive selection by revealing signatures of local adaptation in the form of subtle allele frequency changes correlating with a relevant covariable (Supplemental Note 6.4.4). Indeed, the GEA candidate SNPs differ consistently in allele frequency with respect to habitat but generally do not have large frequency differences between populations (Fig. S45A). This is consistent with local adaptation in chimpanzees being mostly polygenic and driven by soft sweeps as observed in humans (*32*, *117*, *118*), and suggests the presence of complex genetic adaptations even in the absence of fixed differences among populations.

Our findings suggest that while behaviours such as tool use (*16*, *119*) and thermoregulatory behaviours (*17*) are important in mitigating environmental stressors, selective pressures associated with habitat still appear to drive genetic adaptation in chimpanzees. Thus, both behavioural flexibility and genetic adaptation may explain how chimpanzees inhabit such a range of habitats. Far from replacing genetic adaptation, behavioural adaptations may drive genetic changes via gene-culture coevolution (*120*), as seen with human diets (*43*), whereby behavioural flexibility facilitates exposure to novel selection pressures that later drive genetic adaptations.

The evidence of genetic adaptation in forests demonstrates the importance of novel adaptations even in habitats with high availability of resources that support high population densities. This is perhaps because the struggle against the high pathogen load of lowland forests shapes the evolution of these populations. This is not surprising, as pathogens have been important selective pressures for chimpanzees over longer time scales (*65*, *66*). Today, infectious diseases are major causes of chimpanzee population decline(*1*) and recent increased exposure to humans has led to an increase in deadly outbreaks caused by cross-species transmission (*121*). Our findings highlight the importance of genetic adaptation in shaping infectious disease mortality in chimpanzees and suggest that individuals are adapted to the pathogens present in their local habitat, emphasising the dangers of displacement and environmental change.

Evidence of adaptation to malaria in forests is particularly interesting. A range of malaria parasites infect wild chimpanzees, including three *Laverania* species closely related to *P. falciparum*, which originated in gorillas (*57*, *122*) and is now responsible for 90% of global malaria mortality in humans (*123*). However, the fitness effects of malaria in wild chimpanzees are poorly understood (*124*). Its high prevalence in wild populations (*122*) and the few studies of captive chimpanzees suggest that severe effects are rare (*125–128*). However, fitness effects in the wild may be more severe than in captivity, as demonstrated by the largely asymptomatic SIV infections in captivity (*58*, *129*, *130*) but fitness effects in the wild (*29*, *131*). Young chimpanzees and pregnant mothers are particularly susceptible to malaria infection (*132*, *133*), which may lead to higher morbidity/mortality as observed in humans (*134*). Our findings indicate that malaria has been an important selection pressure in the recent past and may have important fitness effects in present-day wild populations. That adaptation appears to be mediated by the same few genes in chimpanzees and humans is striking from an evolutionary point of view; further, it demonstrates how understanding chimpanzee evolution can inform human medicine.

Chimpanzees also appear to have adapted genetically to savannah habitats, although identifying key selective pressures and adaptive traits is harder. Studying additional savannah populations would help address this question. This would provide insights into how our ancestors may have adapted to similar habitats, and have important implications for the conservation of wild chimpanzees as their habitats become hotter and more seasonal under climate change (*2*).

Just as previous studies highlighted the importance of conserving behavioural diversity (*16*, *135*, *136*), we emphasise the importance of conserving adaptive genetic diversity across chimpanzees’ ecological range to maintain their adaptive potential and ensure long-term survival in the wild (*46*, *74*). This is important because chimpanzee habitats are changing rapidly due to direct anthropogenic destruction (*1*, *137*, *138*), climate change (*139*) and disease transmission (*121*, *140*). We emphasise the need to protect all chimpanzee habitats and to consider local genetic adaptations when planning conservation efforts to ensure that individuals are adapted to the local environment. Finally, our study demonstrates the value and promise of non-invasive sampling to investigate genetic adaptation in endangered species.

## Acknowledgements

We would like to thank Laia Llovera for assistance in the laboratory. We thank Thierry Aebischer, Floris Aubert, Emmanuel Ayuk Ayimisin, Arcel Bamba, Matthieu Bonnet, Chloe Cipoletta, Katherine Corogenes, Bryan Curran, Lucy D’Auvergne, Jean Claude Dengui, Theophile Desarmeaux, Karsten Dierks, Emmanuel Dilambaka, Dervla Dowd, Andrew Dunn, Manasseh Eno-Nku, Theo Freeman, Annemarie Goedmakers, John Hart, Martijn Ter Heegde, Inaoyom Imong, Mohamed Kambi, Laura Kehoe, Vincent Lapeyre, Vera Leinert, Joshua M Linder, Sergio Marrocoli, Michael Masozera, Richard McElreath, Vianet Mihindou, Geoffrey Muhanguzi, Felix Mulindahabi, Mizuki Murai, Protais Niyigaba, Nadege Wangue Njomen, Nicolas Ntare, Bruno Perodeau, Alhaji Malikie Siaka, Alexander Tickle, Els Ton, Richard Tshombe, Hilde Vanleeuwe, Paula Álvarez Varona, Virginie Vergnes, Magloire Kambale Vyalengerera and Klaus Zuberbuehler for assistance in field site coordination and sample collection. We thank the field assistants and volunteers from the Jane Goodall Institute Spain in Senegal for their help with sample collection in Dindefelo. We also thank the Greater Mahale Ecosystem Research and Conservation (GMERC) field assistants who provide crucial support and data collection at the Issa Valley field site in Tanzania.

We thank the following government agencies for their support in conducting field research in their countries: Ministère de la Recherche Scientifique et de l’Innovation, Cameroon; Ministère des Forêts et de la Faune, Cameroon; Ministère des Eaux et Forêts, Cote d’Ivoire; Ministère de l’Enseignement Supérieur et de la Recherche Scientifique in Côte d’Ivoire; Institut Congolais pour la Conservation de la Nature, DR-Congo; Ministère de la Recherche Scientifique, DR-Congo; Agence Nationale des Parcs Nationaux, Gabon; Center National de la Recherche Scientifique (CENAREST), Gabon; Société Equatoriale d’Exploitation Forestière (SEEF), Gabon; Department of Wildlife and Range Management, Ghana; Forestry Commission, Ghana; Ministère de l’Agriculture de l’Elevage et des Eaux et Forets, Guinea; Instituto da Biodiversidade e das Áreas Protegidas (IBAP), Guinea-Bissau; Ministro da Agricultura e Desenvolvimento Rural, Guinea-Bissau; Forestry Development Authority, Liberia; Eaux et Forets, Mali; Ministre de l’Environnement et de l’Assainissement et du Developpement Durable du Mali; Conservation Society of Mbe Mountains (CAMM), Nigeria; National Park Service, Nigeria; Ministère de l’Economie Forestière, R-Congo; Ministère de le Recherche Scientifique et Technologique, R-Congo; Ministry of Education, Rwanda; Rwanda Development Board, Rwanda; Direction des Eaux, Forêts, et Chasses, Senegal; Reserve Naturelle Communautaire de Dindefelo, Senegal; Ministry of Agriculture, Forestry, and Food Security, Sierra Leone; National Protected Area Authority, Sierra Leone; Tanzania Commission for Science and Technology, Tanzania; Tanzania Wildlife Research Institute, Tanzania; Makerere University Biological Field Station (MUBFS), Uganda; Uganda National Council for Science and Technology (UNCST), Uganda; Uganda Wildlife Authority, Uganda; National Forestry Authority, Uganda; Agence Congolaise de la Faune et des Aires Protégées; and The Wild Chimpanzee Foundation (WCF).

We thank Dean Lavelle for assistance in re-base calling ONT data using guppy 5.0.11, as well as Agilent for their collaboration in the project. We thank Hernán Burbano, Max Reuter and all the members of the Andrés and Burbano groups for useful comments and suggestions. Finally, we thank Jinliang Wang for his support and supervision.

## Author contributions

Conceptualisation: AMA, TM-B, MA, CB

Formal Analysis: HJO, CF, DCS, AMA, MYD, TM-B

Methodology: HJO, AMA, CF, DCS, MYD, JMS, TM-B

Investigation:

Project design and planning: JMS, LV, CB, CF, EL, TM-B, MA, HSK, AMA

Bioinformatic processing: HJO, CF, DCS

Interpretation of results: HJO, AMA, CF, JMS, CDB, PG, AP, HSK, TM-B, MA, MYD, DCS

Supervision or execution of data collection: CF, EL, MA-E, PD, AA, SA, AKA, EB, DB, MB, GB, RC, HC, CC, ED, A-CG, JH, DH, VH, RAH-A, KJJ, SJ, JJ, PK, MK, AKK, Mbangi Kambere, IK, DK, KEL, JL, BL, AL, KCL, ML, GM, RM, AM, DM, EN, SN, Stuart Nixon, Emmanuelle Normand, CO, LJO, RO, LP, JP, SR, MMR, AR, CS, LS, VS, FS, NT, LRT, JvS, EV, ECW, JW, RMW, YGY, KKY, LV, AP, CB, HSK, MA

Visualization: HJO, DCS

Funding acquisition: TM-B, MA, AMA, MYD, HSK, CB

Project administration: AMA, MA, HSK, TM-B, CB

Supervision: AMA, MA, TM-B

Writing – original draft: HJO, AMA

Writing – review & editing: All co-authors

## Data and materials availability

All data and code will be made available on publication.

