## Supplementary Materials for "Local genetic adaptation to habitat in wild chimpanzees"

**The PDF file includes:**

Materials and Methods
Supplementary Text
Figs. S1 to S56
Tables S1 to S4
References

**Other Supplementary Materials for this manuscript include the following:**
N/A

### Materials and Methods

#### *Sampling, DNA extraction and sequencing*

Sampling, DNA extraction and identification of unique individuals from samples were performed as described in (*81*). These steps plus library preparation and pooling were performed as described in (*3*) since both studies include the exact same samples (an in-depth description of the methods used can be found in supplemental information from (*3*)). Briefly, faecal DNA was extracted from 5,397 PanAf samples and screened with microsatellite genotyping (*81*) to select samples with good-quality DNA while discarding repeated individuals and first-order relatives. 828 samples across all four subspecies were then sequenced, representing 147 central, 209 eastern, 86 Nigeria-Cameroon and 386 western chimpanzees, with a minimum of 20 individuals per sample site when possible, over 52 sample sites. Library preparation was performed on different days for random batches of 24–48 samples, with a unique double-inline barcoded library per sample following the BEST protocol with minor modifications (*77*, *141*). Pooling for capture was devised by host DNA content (fraction of chimpanzee DNA, relative to gut microbial and exogenous DNA), with 30 samples/pool (*3*). Each capture pool was divided into two main aliquots (one for chr21 (*3*) and one for the exome) and subsequently into several aliquots for hybridisations as in (*3*). Target hybridisation capture was performed separately to retrieve the non-repetitive regions of chr21 (*3*) and the exome (this study) using the SureSelect Human All Exon V6 RNA library baits from Agilent Technologies. Sequencing was done on 27 lanes of a HiSeq X, 2x150.

#### *Data processing and read filtering*

Demultiplexing, filtering and read mapping were done as in (*3*) (detailed information in (*3*) supplemental information) and is briefly described in Supplemental Note 1.1. Reads were mapped to the human genome hg19 (GRCh37, Feb.2009 (GCA_000001405.1)) because the high quality of hg19 and its extensive annotation make this a better option than mapping to the chimpanzee reference genome. Mapping to the human genome also avoids the potential risk of subspecies-specific reference biases, since the chimpanzee reference genome was generated from a single western individual. The resulting BAM files contained only reliable on-target reads which were used for all downstream analyses.

#### *Sample filtering*

Filtering followed (*3*), which is briefly described here and in detail in Supplemental Note 3. We identified genetic outliers to find problematic samples using PCAngsd v3 (*142*) with genotype likelihoods estimated in ANGSD v0.933 (*143*); PC1 outliers were manually removed (n=107). We excluded samples with >1% human contamination as estimated by HuConTest (*144*)) (n=188) and then those with mean per-base read depth <0.5x (n=66). Identical samples and first-degree relatives were removed using NgsRelate (*145*) (n=47). PCAngsd was run once more to remove samples that failed to cluster with their respective sample sites (n=5). The filtered exome dataset contains 415 samples from 44 sample sites across all four subspecies. To have a comparable dataset, we generated a chr21 dataset with all these samples except those with high contamination or erroneous PCA clustering using the chr21 dataset (n=3). The filtered chr21 dataset contains 412 samples from 44 sample sites across all four subspecies.

##### *Estimating derived allele counts*

We investigated population structure in the exome data using PCAngsd and NGSadmix (*146*) (Supplemental Note 4). Following (*3*, *81*), we combined sample sites less than 15 km apart because the frequent movement of females between nearby communities (i.e. social groups) over these distances means that they cannot be considered genetically distinct. Population structure analyses confirm these sample sites as very closely related (*3*) (Figs. S12-16). We, therefore, combined the five Comoé sample sites, the two Taï sample sites, and Bakoun with Sobory, all of which belong to the western subspecies. In addition, we combined two Nigeria-Cameroon sample sites, Korup and Mt. Cameroon, because PCAangsd and NGSadmix showed that the Mt. Cameroon samples lay within the variation of Korup (Figs. S12-16), and because chr21 analysis of identical by descent segments indicated very high connectivity until only ~600 years ago (*3*). We note that combining populations may limit the power to identify putative differential adaptations but never create false positives. The resulting dataset contains 37 genetic units that we refer to as 'populations'. Populations with sample sizes lower than 8 (n=7) were then excluded, resulting in a final filtered dataset of 388 samples (385 for chr21) from 30 populations (5 central, 9 eastern, 2 Nigeria-Cameroon and 14 western) (Fig. 3, Fig. S20).

We estimated the population minor allele frequency (MAFs) of each autosomal SNP in each population from genotype likelihoods using ANGSD (*143*) (Supplemental Note 5). MAFs were estimated only for genomic sites with at least one read in at least six samples or half the samples from that population, whichever was larger (i.e., more conservative). Sites with a MAF lower than 1/2N (N=number of individuals with at least one read at a given genomic site) or a p-value of a site being monomorphic within a population greater than $10^{-6}$ were considered monomorphic within that population. Minor allele counts (MACs) were obtained by multiplying the estimated allele frequencies by 2N. Alleles were polarised according to the ancestral state from the EPO alignment of six primate species (ftp://ftp.ensembl.org/pub/release-75/fasta/ancestral_alleles/homo_sapiens_ancestor_GRCh37_e71.tar.bz2), sites with missing ancestral state data were excluded. This information was used to generate estimated derived allele counts (DACs) for each population.

We created four 'subspecies-datasets'; *All*, *Central-Eastern*, *Nigeria-Cameroon* and *Western*. Sites with MAC lower than 2 in a subspecies-dataset were removed to discard sequencing errors and, in any case, we cannot identify signatures of positive selection in exceedingly rare variants. In fact, this filter greatly improves the shape of the $X^tX^*$ distribution to the expectation, likely because it removes sequencing errors. In each subspecies-dataset, sites with allele count data in less than 70% of populations were also removed. The same process was applied to the chr21 data. This chromosome was then filtered to obtain a 'non-genic-chr21' dataset containing only regions >1kb from a gene using BEDTools intersect v2.29.2 (*147*) and the hg19 annotation file downloaded from Ensembl (http://ftp.ensembl.org/pub/grch37/current/gtf/homo_sapiens/Homo_sapiens.GRCh37.87.chr.gtf.gz). The number of populations and SNPs per subspecies-dataset for the exome and non-genic-chr21 data is shown in Table S1 and Figs. S19-20.

##### *Environmental data*

Large-scale biogeographic analysis of Africa identifies forest, savannah and intermediate bistable biomes within the chimpanzee range (*87*). Forest and savannah are at the extreme ends of the chimpanzee habitat gradient and have very different tree species compositions (*87*). The percentage of trees identified as forest specialists was thus used to reflect habitat. Forest-tree-percentage was calculated as the number of forest tree specialists divided by the total number of forest specialist, generalist and savannah specialist trees as classified in (*87*). The proportion of trees that could not be assigned to one of these categories varies greatly between populations so unclassified trees were excluded to avoid introducing noise or biases in the habitat statistic (Supplementary Note 2). While a single variable cannot fully describe the nuances of chimpanzee habitats, forest-tree-percentage is a good proxy for many potential selection pressures and better represents the habitat gradient than the discrete categories used in previous studies (*12*). When sample sites were combined to form populations, we used the mean values. The variable was imputed in one missing eastern population (Chinko) using missForest (*148*) using all PanAf environmental data and publicly available environmental data (Supplemental Note 2). The imputed value for Chinko (75.36%) is consistent with its classification as a bistable forest (*87*).

##### *BayPass*

We identified signatures of natural selection using BayPass v2.2 (*83*) (Supplemental Note 6). The covariance matrices for each subspecies-dataset were first estimated under the core model with default parameters. Because this step requires no missing data, we used only SNPs with no missing allele count data in any population. This left ample SNPs to estimate the covariance matrix (Table S1).

To account for run-to-run variation, BayPass was run three times with different seeds using the -seed option. Estimated model hyperparameters were consistent across independent runs. Correlation matrices, hierarchical clustering trees and PCAs were calculated from the covariance matrices and PCAs run for visualisation (Supplementary Note 6.1).

To perform the genetics-only test, BayPass was run on each subspecies-dataset under the core model using the corresponding allele count dataset and estimated covariance matrix. This stage is robust to missing data (*83*, *85*), so we retained sites with missing data. BayPass was run three times with different seeds using the three covariance matrices previously estimated using the same seed. The median $X^tX^*$ value of each SNP across these runs was used to select candidates. $X^tX^*$ values are expected to follow a $\chi^2_J$ distribution ($J$=number of populations) under the core model; however, violations of the assumption of normally distributed allele frequencies can lead to a poor fit. We, therefore, used the $X^tX^*$ distribution from the non-genic-chr21 data as our null distribution because these regions are expected to evolve mostly neutrally, with positive selection mainly targeting the exomes and neighbouring genomic elements (Supplementary Note 6.2). We note that this is a conservative null as there may be some sites evolving under positive selection in non-genic regions of chr21. The distribution of $X^tX^*$ values from non-genic-chr21 was thus used to define $X^tX^*$ thresholds according to estimated false positive rates (FPRs). To account for any putative effect of read depth on signatures of selection, SNPs were divided into five depth bins and the FPR was calculated for each of these bins. Per-site depth was calculated as the total sequencing depth across all samples in the subspecies-dataset. Candidates were selected at three thresholds corresponding to estimated FPRs of 0.5%, 0.1% and 0.05%.

We performed a GEA by running BayPass on each subspecies-dataset under the AUX model using the corresponding allele count dataset, estimated covariance matrix and standardised population environmental data (using the -scalecov option) as input. As with the genetics-only test, BayPass was run three times with different seeds and median values across three independent runs were calculated and used to select candidates using the same method only using BF rather than $X^tX^*$.

To verify that BayPass correctly accounts for population structure, we investigated the allele frequency patterns in the candidate SNPs, both unstandardised (closely related to the observed allele frequencies with missing data imputed using the covariance matrix) and standardised (which account for neutral population structure). For both statistics, we calculated correlation matrices among populations and performed hierarchical clustering and k-medoids clustering in R, using the allele frequencies from a single BayPass run under the core model.

##### *Gene set enrichment*

We annotated SNPs using BEDTools intersect v2.29.2 (*147*) and the hg19 annotation file. SNPs were assigned to a gene if they lay within the gene coordinates ±5kb. Gowinda (*149*) was run to test for enrichment of gene categories in our candidate SNPs while accounting for gene length and overlapping genes (method details in Supplemental Note 7), in 'gene' mode to also account for linkage disequilibrium. We used the same Ensembl hg19 annotation file as above and restricted our analysis to only genes with 1-1 chimpanzee-human orthologs (*65*).

Hypothesis-free candidate gene enrichment tests were run using Gene Ontology (GO) categories (*150*), KEGG pathways (*151*), Reactome categories (*152*), human GWAS traits (*153*), Phenotype database traits (*154*) and tissue expression data from the Human Protein Atlas (*155*), where genes were considered associated with a tissue if expression level was 'high' and reliability was 'approved'. We hypothesised that pathogens and dehydration stress may be important selection pressures so we tested for enrichment of immunity genes (*93*, *94*), viral interacting proteins (VIPs) (*156*, *157*), other pathogen-related genes (SIV/HIV (*158–160*), malaria (*96–98*), influenza (*161*), SARS-Cov-2 (*162–166*), HSV-1 (*167*), anthrax (*168*) and ebola (*169*)), and genes involved in response to dehydration (*89*, *90*) (details in Supplemental Note 7). Gowinda accounts for multiple testing within each run; no additional corrections were performed across runs.

##### *GYPA structural variation analysis*

The human reference genome and panTro6 were retrieved from the UCSC Genome Browser FTP website. AG18354 primary and alternate assemblies were obtained from PRJNA916736 and PRJNA916737 (https://github.com/marbl/Primates). PacBio CLR data from Clint/panTro6 were obtained from PRJNA369439, ONT reads from AG18359 were obtained from PRJEB36949, and PacBio HiFi reads from AG18359 were downloaded from the GenomeArk of the Primates Telomere-to-Telomere Consortium (https://genomeark.github.io/t2t-all/Pan_troglodytes.html). AG18359 assembly was generated using previously published ONT data by re-calling bases using guppy5.0.11 and running the Shasta assembler with default parameters (*170*). Assemblies were mapped to hg38 using minimap2 with -x asm5 parameter, while long reads were mapped using minimap2 v2.26 with settings -x map-pb, -x map-hifi, -x map-ont, respectively, and default parameters otherwise. Contigs and reads mapped to hg38 were visually examined using the Integrative Genome Browser. Human gene annotations were obtained from Gencode v43, and litfed from hg38 to chimpanzee assemblies using liftoff (*171*), enabling the detection of additional gene copies with parameters: -copies -sc 0.9.

High-coverage short-read data from 60 chimpanzees were retrieved from ENA BioProject PRJEB15086 and PRJNA189439. Gene-family copy-number estimates were obtained in hg38 coordinates across 1-kbp windows using the fastCN pipeline (*172*), which utilizes MrsFast (*173*) to perform short-read multimapping. Copy-number genotyping was obtained using a custom Python script that uses pybedtools package to select 1-kbp windows intersecting regions of interest and calculates median copy number. SNP genotypes were called using the high-coverage short-read data aligned to hg19 with Genome Analysis Toolkit (GATK) v4.2.5.0 (*174*) following GATK best practices (*174*). Variants were first called for each sample separately using GATK HaplotypeCaller resulting in a GVCF per sample. GVCFs were consolidated into a VCF using GATK GenomicsDBImport and joint genotype calling was performed using GATK GenotypeGVCFs. Variants were filtered only for genotype quality of at least 30. wANNOVAR (*175*, *176*) was used to predict the functional consequences of the candidate SNPs.

#### Supplementary Text

#### 1. Genomic data

An unavoidable limitation of our study is the use of non-invasive sampling which is the only option available for obtaining genomic data from wild individuals for many protected species, including chimpanzees. However, our carefully designed methodology minimises the risk of false positives. There is also no reason to expect sequencing errors to result in allele frequencies correlating with habitat type or to occur more in genes with particular functions. Despite the inherent challenges of non-invasive sampling, we demonstrate that genomics can provide important insights into patterns of local adaptation in an endangered species where it is impossible to obtain invasive samples of wild individuals.

Coding regions are more likely to contain functional variants and so exome sequencing is an economical method for investigating genetic adaptation. However, exome sequencing does not cover non-coding regions, such as introns, enhancers or promoters which may alter gene expression. Nevertheless, recent selective events substantially increase linkage disequilibrium and thus the exome allows us to identify signatures of selection at neighbouring functional sites.

##### *1.1. Demultiplexing, filtering and read mapping*

We demultiplexed libraries belonging to the same hybridization pool using Sabre (https://github.com/najoshi/sabre). Illumina adaptors and bases with average quality scores <20 were removed with Trimmomatic (version 0.36) (*177*). Reads were mapped to the human genome hg19 (GRCh37, Feb.2009 (GCA_000001405.1)) using BWA (version 0.7.12) (*178*). Duplicates were removed using PicardTools (version 1.95) (http://broadinstitute.github.io/picard/) and further read filtering using samtools (version 1.5). Off-target reads were removed using BEDTools intersect v2.22.1 (*147*) and the Agilent Exome V6 target space bed file.

#### 2. Environmental data

To investigate potential selection pressures driving local adaptation, the spatial resolution of genetic and environmental data should ideally be the same. For every sample site with genetic data, we also have dozens of environmental variables recorded by field workers or remote sensing. This dataset is unprecedented in its scale, increasing our power to detect local adaptation in chimpanzees and aiding the identification of likely selection pressures.

We decided to use a single measure of habitat type for the genotype-environment association analysis (below). Using such a composite measure summarises a range of environmental variables and allows us to contribute to literature investigating adaptations to habitat types in chimpanzees (*2*, *48–51*). Two types of floristic habitat type were recorded by PanAf field workers at each sample site. The first measures the percentage of tropical forest, mosaic-forest or savannah habitat along a transect; this data was missing for 4 sample sites (Chinko, Gishwati,

Comoe2 and ComoeCNPN). The second measures the number of forest specialist, generalist or savannah specialist trees as defined in (*87*), this data was missing for 3 sample sites (Chinko, Comoe2 and ComoeCNPN).

In cases where sample sites were combined to form 'populations' (see below), the mean value for all sample sites within a population was used. Because three other Comoé sample sites had habitat data, only Chinko and Gishwati remained with missing habitat data. Missing data was imputed using missForest (*148*) using all the PanAf environmental data (http://panafrican.eva.mpg.de/english/approaches_and_methods.php) in addition to published and publically available environmental data (elevation (https://biogeo.ucdavis.edu/data/), percentage tree cover (Hansen et al., 2013; http://earthenginepartners.appspot.com/science-2013-global-forest/download_v1.7.html), human footprint (Global Human Footprint Dataset v.2; https://doi.org/10.7927/H4M61H5F), habitat stability measures (*13*) and 19 climatic variables at a resolution of 2.5 minutes from WorldClim (https://www.worldclim.org/)). Imputed values were consistent with the biomes classified by (*87*).

Tree species composition is highly divergent between forest and savannahs (*87*) and the vegetation data is more complete than the transect data, therefore, we decided to use either the percentage of forest or savannah specialist trees as a measure of habitat type. Unsurprisingly, these two measures are highly negatively correlated across all our sites (Pearson r=-0.97, $p=2.200\times10^{-16}$). After visual inspection, the percentage of forest tree specialists was chosen as it best separates sites by the biomes inferred from a large-scale biogeographic analysis (*87*) and every population bar one (Kayan) had some forest specialist trees (in contrast, six sites have no savannah specialist trees).

##### *2.1. Unclassified trees*

Only trees classified as being forest specialists, savannah specialists or generalist species in (*87*) were considered in the calculation of forest-tree-percentage, with unclassified trees considered discarded. This is because there is substantial variation in the number of unknown trees between populations (Fig. S1), with some populations having as much as 45% of trees unclassified, which brings noise and potential biases to the statistic. Indeed, running the BayPass analysis using a related measure of forest-tree-percentage that includes unknown trees (i.e. forest/(forest+generalist+savannah+unclassified)) results in no excess of sites with high BFs in the exome compared to non-genic-chr21. We note that highly noisy or erroneous habitat measures would be expected to generate a false negative result (with noise hiding a true correlation) but not a false positive in the GEA. This indicates that unknown trees can contribute substantially to the measure in this dataset and that including unknown trees results in a measure that does not correspond closely with selection pressures.

#### 3. Sample filtering

Non-invasive sampling is necessary for the genetic sampling of endangered, elusive, unhabituated wild populations. However, such sampling strategies result in additional challenges which may compromise the quality of the genetic data. Faecal samples contain low amounts of endogenous DNA often resulting in low coverage, our unfiltered samples have a median coverage of 1.51-fold (0.00- to 69.50-fold). Samples may also have been exposed to the elements for hours before being collected and the hot humid conditions of the tropics facilitate bacterial growth and DNA damage. (*3*) found damage patterns (increased T-to-C and A-to-G substitutions) similar to those in ancient DNA, but the error rates were an order of magnitude lower. Faecal samples are also susceptible to contamination from the individual's diet. Chimpanzees often prey on other primates so DNA from an individual's diet may be captured, sequenced and mapped to the reference genome (Hg19). There is also potential for sample misidentification at the moment of collection, resulting in the sampling of different species (*3*). These challenges mean sample filtering is particularly important to remove low-quality, misidentified and contaminated samples which could add noise to the analysis. We used the same filtering methodology as for chr21 data from the same samples (*3*) summarised in Fig. S7.

##### *3.1. PC1 outliers*

First, samples showing signatures of contamination were removed by excluding those which were outliers in a PCA. Because all endogenous DNA should belong to chimpanzees and non-endogenous to other species, PC1 is expected to discriminate contaminated samples from samples with very low levels of contamination. BAM files containing only reliable on-target reads were input into ANGSD v0.933 (*143*) to estimate genotype likelihoods. The following parameters were used every time ANGSD was run: -uniqueOnly 1 -remove_bads 1 -minMapQ 30 -only_proper_pairs 1 -C 50 -baq 1 -skipTriallelic 1 -GL 2 -doMajorMinor 1. These parameters, hereafter 'standard parameters', include only reads with a single best hit, mapping quality information, a minimum mapQ quality of 30 and pairs of reads with both mates mapped correctly. mapQ is adjusted for excessive mismatches, per-Base Alignment Quality is calculated, and genotype likelihoods were estimated using the original GATK model (-GL 2). In addition to the standard parameters, the following parameters were used: -minInd 15 -minMaf 0.01 -SNP_pval 1e-6 -doHWE 1 -minHWEpval 0.001 -doGlf 2. These 'additional parameters' retained only sites that were present in ≥ 15 individuals, biallelic, had a minor allele frequency (MAF) ≥ 0.01, a probability of being monomorphic < $10^{-6}$ and a probability of < $10^{-3}$ of deviating from Hardy-Weinberg equilibrium (HWE). Genotype likelihoods were outputted as beagle files. The PCA was performed using the resulting genotype likelihoods and PCAngsd v3 (*142*) with default parameters.

PC1 explains 15.34% of the genetic variation (Fig. S2). The density of PC1 values shows two clear peaks, one corresponding to western samples and another corresponding to non-western samples, with a long tail of samples which did not cluster with the rest of the chimpanzees (Fig.

S2). This density distribution was used to define the threshold of -0.01 to remove samples in this tail of the distribution which are likely to be contaminated resulting in 721 samples passing this filter. Of the 107 samples removed at this stage, 94 were also removed at this stage of filtering in (*3*) who found that many of these samples were contaminated with DNA mapping to monkeys (likely due to diet), gorillas (due to sample misidentification) and humans (likely due to contamination during handling and sample misidentification in one case). PC2 explains 9.34% of the genetic variation and begins to separate the four subspecies. This filtering step resulted in a dataset of 721 samples.

##### *3.2. Human contamination*

We also removed samples with over 1% human contamination estimated using HuConTest (*144*) as in (*3*). Briefly, human contamination was estimated by investigating genomic positions where humans and chimpanzees consistently differ based on diversity data from high-coverage chimpanzee genomes (*62*) and the 1000 genomes project (*179*). The proportion of chimpanzee alleles to human alleles provides a reliable estimate of the percentage of human contamination in the sample. We retained samples with < 1% human contamination (Fig. S3), consistent with (*3*), resulting in the exclusion of 188 samples. This step resulted in a dataset of 533 samples.

##### *3.3. Coverage*

The remaining 533 samples with low levels of contamination were filtered for coverage. Although we use genotype likelihood (GL) based methods to estimate allele frequencies, ultra-low coverage samples (< 0.5x) were filtered out because they would only increase noise (Fig. S4). Coverage was calculated as the average read depth per site in the on-target region (exome). This filter removed 66 individuals leaving 467 samples.

##### *3.4. Relatedness*

Samples were then filtered to remove those which were a first-order relative or from the same individual as another sample following (*3*). We used NgsRelate v2 (*145*) to estimate the coefficient of kinship (θ) (*180*). NgsRelate was run on each sample site individually to avoid confounding effects of population structure. ANSGD was run with the standard parameters (defined above) with the additional parameters -doMaf 1 -minMaf 0.05 -SNP_pval 1e-6 -doGlf 3 (consistent with (*3*)) to only include sites with a MAF ≥ 0.05 and a probability of being monomorphic < $10^{-6}$ and outputs the GLs in the appropriate format for NgsRelate. Samples from the same individual would be expected to result in θ=0.5, first-order relatives would be expected to result in θ=0.25 and so on. We considered any pair of samples with θ>0.1875 as first-order relatives to account for variation in the estimates of θ (0.1875 is the midpoint between first-order relatives (θ=0.25) and second-order relatives (θ=0.125)). Most cases were simple pairs of related samples with five groups of three related samples (Fig. S5). For every group of related samples, only the highest coverage sample was kept. This resulted in the exclusion of 46 samples leaving 421 samples.

##### *3.5. Samples and population structure*

The final stage of filtering consisted of running population structure analyses to identify samples which do not cluster with their correct subspecies and sample sites. A sample failing to cluster as expected using exome-wide data is likely a sign of a problem (e.g. borderline low coverage or mislabelling), and in any case, they cannot be considered as part of their labelled sample site. We ran ANGSD on the filtered samples with the standard parameters and additional parameters SNP_pval 1e-6 -minMaf 0.01 -doHWE 1 -minHWEpval 0.001 on all samples and for each of the four subspecies individually. GLs were then input to PCAngsd to perform PCA and the ANGSD tool NGSadmix (*146*) to estimate individual admixture proportions.

We identified one western sample (Fjn2-62) and two central samples (Con2-57 and GB-14-05) which failed to cluster with their respective subspecies and so they were removed (Fig. S6). Within westerns, two samples (Fjn3-24 and Gep2-41) failed to cluster with any sample site and so were also removed. These patterns could not be explained by recent migration between sample sites as samples were separated from the most likely potential sources of migration by large geographic distances (Fig. 1). These samples were not outliers for coverage or human contamination statistics and so it is unclear why they did not cluster as expected. Two mislabelled samples were identified; the sample reportedly from Campo Ma'an (CMNP1-24) clustered with the southern sample sites Conkouati and Loango when it would be expected to cluster with other northern sample sites; and a western sample reportedly from Sangaredi (Gco4-2) clearly clustered with samples from Mt Sangbe. CMNP1-24 likely belongs to Conkouati as analysis of rare alleles on the full chr21 found this to be the most likely true sample site for this sample; however, a relatedness analysis on Conkouati identified CMNP1-24 as a first-order relative of another Conkouati sample with higher coverage and CMNP1-24 was filtered out. Gco4-2 was reassigned to Mt Sangbe and is not a first-order relative of any other sample at this sample site. The final filtered dataset contained 415 samples from 44 sample sites across all four subspecies (Fig. S8) with a median coverage per sample of 4.96-fold (0.51- to 69.50-fold) in the exome target space.

##### *3.6. Chr21*

In order to have a comparable chr21 dataset, the chr21 data was filtered to only include samples which passed the exome filtering steps. In addition, we identified and removed three chr21 samples with evidence of contamination on the chr21, one sample with human contamination > 1% (Fig. S9) on the chr21 and two samples which did not cluster with their sample sites (Fig. S10) based on chr21. Four of the remaining samples had a mean coverage < 0.5x (Fig. S11), however, these were not removed in the chr21 dataset as they only narrowly missed the 0.5x threshold (>0.3x) and chr21 data is only used to generate null distributions rather than to identify sites under selection.

#### 4. Population structure

When running analyses to identify the signatures of local adaptation, it is important to understand population structure to avoid false positives and define genetic populations for estimating population allele frequencies. We also aimed to ensure that the population structure in the exomes corresponds closely with the reported genetic substructure in chr21 from the same samples (*3*). ANGSD, PCAngsd and NGSadmix were run as described above (Supplemental Note 3.5) using samples which passed all filtering stages.

##### *4.1. Exomes*

PCAs from the exome data agree closely with (*3*), including a similar proportion of the genetic variance being explained by the first two principal components, with slight differences likely due to small differences in sample filtering (Fig. S12). The PCA including all samples shows clear grouping into the 4 subspecies. PCA of centrals shows the northern and southern clades separated by the Ogooué river described by (*3*) along PC1. PC2 separates Mts de Cristal and Goualougo which lie at the far west and east of the northern clade distribution respectively but explains little variation in the southern clade. This is consistent with recent population connectivity in the southern clade described by (*3*) reducing population differentiation. PC1 for eastern populations shows a general north-south cline with Issa Valley being particularly distinct as Lake Tanganyika acts as a barrier to gene flow (*3*). Nigeria-Cameroon PC1 separates Gashaka from the other three sample sites. The three Mt. Cameroon samples lie fully with the diversity of Korup consistent with patterns of recent gene flow between these populations identified by (*3*). Western PCA shows less separation between sites with more overlap due partly to the higher sampling density in this subspecies. PC1 roughly reflects an east-west cline with Comoé sample sites as clear outliers.

Procrustes transformation of the first two principal components onto a map shows a pattern mostly consistent with isolation-by-distance within subspecies (Fig. S13) with notable exceptions such Korup lying within the variation of Mt. Cameroon despite being 90 km apart and more isolated populations such as Issa Valley and Mt. Sangbe. Sample sites in the southern central clade do not show isolation-by-distance using the first two principal components because PC1 separates the northern and southern clades while PC2 separates sample sites in the northern clade which has higher differentiation between sample sites (*3*).

##### *4.2. Chr21*

PCAs using chr21 agree very closely with results using the exome including the percentage of genetic variation explained by the first two principal components.

##### *4.3. Defining populations*

Investigating natural selection requires information about allele frequencies in the form of population allele counts. Informed by our demographic analyses and the results of (*3*) and (*81*), sample sites separated by small genetic and geographic distances were combined to better represent genetic units, which we refer to as 'populations'. Combining closely related sample sites increases the sample size per population leading to more accurate estimates of population allele frequency at the cost of reducing the resolution of the analysis.

We combined the five Comoé sample sites, the two Tai sample sites, and Bakoun with Sobory as done by (*3*) and (*81*). These sample sites are < 15 km apart and the frequent movement of females between nearby communities means they cannot be considered genetically distinct populations. These sample sites were also shown to be genetically similar in the population structure analysis (Figs. S12-16) and they inhabit almost identical habitats and so are likely subject to very similar selection pressures. In addition, we also combined the Korup and Mt. Cameroon sample sites together. Korup and Mt. Cameroon are 90 km apart and analyses of identical by descent genomic segments inferred very high connectivity between them until very recently (~600 years ago) (*3*). Exome demographic analyses further support combining them as the PCA and admixture analyses suggest that Mt Cameroon samples lay within the variation of Korup (Figs. S12-16). Using the WorldClim database (https://www.worldclim.org/data/index.html) and biomes from (*16*, *87*) we also confirmed that Korup and Mt. Cameroon have very similar tropical forest habitats. Any small differences in the environmental variables between these sites would be unlikely to have led to differential local adaptation due to the recent high levels of connectivity. The combined sample size of Korup and Mt Cameroon is 10 (Fig. 3 and S17) resulting in the presence of two Nigeria-Cameroon populations in the dataset, (our filter for sample size per population is eight individuals, Supplemental Note 5.1) allowing us to test for selection within this underrepresented subspecies.

#### 5. Estimating derived allele counts

##### *5.1. Estimating minor allele frequencies*

BAM files were inputted into ANGSD to estimate genotype likelihoods and minor allele frequencies (MAF) for each population. ANGSD was run population-by-population with the same parameters as used for the demography analyses with the following exceptions. Sites must have data for at least six individuals or 50% of the total sample size of a population, whichever is larger. A minimum of six was chosen as previous studies on simulated and empirical datasets suggest that this sample size is sufficient for GEA (*181*). We excluded populations which had sample sizes < 8 because population sample sizes close to the minimum required per site (6) resulted in large numbers of sites with missing data. 388 samples from 30 populations remain after this population filter. ANGSD was run with no minimum MAF filter and no filter based on the probability of a site being monomorphic (-SNP_pval 1) to retain sites which are monomorphic within a population as these locally monomorphic sites may prove to be globally polymorphic when compared with other populations. MAFs were estimated from genotype likelihoods by assuming the major allele is known (inferred from genotype likelihoods) and the minor allele is unknown (-doMaf 2). When the minor allele is very rare it is not always clear which base is the true minor allele and so we do not assume we know the true minor allele. The likelihood of a MAF is estimated by summing over the three possible minor alleles weighted by their probabilities and an expectation maximisation (EM) algorithm finds the MAF with the highest likelihood. Total sequencing depth per site was also calculated using -doSnpStat 1. An ancestral state file was also supplied to ANGSD so the ancestral allele was reported in the output and could be used to polarise the allele frequencies. We used the EPO alignment of six primate species aligned to Hg19 as the ancestral state file (ftp://ftp.ensembl.org/pub/release-75/fasta/ancestral_alleles/homo_sapiens_ancestor_GRCh37_e71.tar.bz2).

We conservatively restricted our analysis to the autosomes because our GL-based approach and the uncertain sex of the sampled individuals made it impossible to confidently estimate population allele counts for sex chromosomes. Sex chromosomes are commonly under particularly strong selection (*182*), particularly in species with a high reproductive skew such as chimpanzees (*63*, *64*, *183*) and so analysis of sex chromosomes in this dataset may be an interesting avenue for future research.

##### *5.2. Site frequency spectra (SFS)*

The unfolded site frequency spectra (SFS) conforms to expectations and shows no abnormalities which would indicate biases or errors in the allele frequency estimations (Fig. S18). For example, the *Western* SFS has relatively fewer mid-frequency alleles compared to the central SFS owing to western's lower effective population size (*5*). The exome also has relatively fewer mid-frequency alleles than non-genic-chr21 likely due to the stronger effect of purifying selection in the exome.

Running ANGSD without a HWE filter resulted in an excess of mid-frequency alleles likely due to mapping of chimpanzee paralogs to single genes in the human genome. These sites would be reported as having an excess of heterozygotes and therefore deviate strongly from HWE. Removing sites which deviate from HWE with a p-value≤0.001 removed the excess of mid-frequency alleles. Using the reference or ancestral allele as the major allele (running ANGSD -doMajorMinor 4 or 5) resulted in an excess of high-frequency derived alleles due to reference bias. Inferring the major and minor alleles from the GLs (-doMajorMinor 1) removed the reference bias and excess of high-frequency derived alleles.

### 6. BayPass

#### *6.1. Estimating population covariance matrices*

Visualising the covariance matrices as correlation matrices, hierarchical clustering trees and PCAs (Figs. S21, S22) show clear separation of subspecies, high correlation between western population allele frequencies due to a shared population bottleneck (*3*, *5*, *62*), and strong differentiation between central populations separated by the Ogooué river (*3*).

The covariance matrices from the exome and non-genic-chr21 regions correspond closely but differ likely due to stronger purifying selection and lower linkage disequilibrium in the exomes compared to non-genic regions of chr21. Förstner and Moonen distances (FMD) (*184*) between covariance matrices were computed using fmd.dist(), an R function included in BayPass. Mean FMD between independent runs for exome and non-genic-chr21 were 0.82 and 0.76 for *All*, 0.09 and 0.10 for *Central-Eastern*, 0.02 and 0.02 for *Nigeria-Cameroon*, and 0.03 and 0.05 for *Western*. As expected, the mean FMD between exome and the non-genic-chr21 covariance matrices are larger: 2.51 for *All*, 0.59 for *Central-Eastern*, 1.43 for *Nigeria-Cameroon* and 2.19 for *Western*. These values are in line with those reported in published BayPass analyses (*83*, *185–188*).

#### *6.2. Generating an appropriate null distribution*

In our analyses, SNPs with the highest selection statistics ($X^tX^*$ or Bayes factor) are those most likely to be under positive selection (details below). Candidate targets of selection can thus be chosen simply by selecting SNPs in the top tail of the empirical distribution. However, this method relies on the assumption that there are SNPs under positive selection in our dataset and does not allow the estimation of false positive rates (FPR). Using an appropriate null distribution, tail thresholds can be selected based on estimated FPRs, and are not reliant on the assumption that the dataset contains SNPs under selection. An excess of SNPs in the top tail of the empirical distribution compared to the null expectation would also provide evidence for the presence of positive selection on the genome-scale.

So how can we generate an appropriate null distribution? Statistical methods can in principle be used to estimate the probability that a site is under selection, however, these calculations can be incorrect if improperly calibrated and are reliant on strong assumptions. For example, BayPass assumes that allele frequencies are normally distributed which is often violated in practice, particularly when sample sizes are small. A second option is to simulate a null dataset under a neutral model and use statistics estimated from this simulated dataset as a null distribution. The BayPass function simulate.baypass() generates pseudo-observed data sets (PODs) under the core model for this purpose. However, such methods are also reliant on the assumptions of the model used to simulate the data such as normally distributed allele frequencies in the case of BayPass.

An alternative approach is to generate an empirical null distribution by calculating summary statistics on an empirical dataset that is not subject to natural selection. A key advantage of this approach is that it does not rely on model assumptions. A suitable empirical null distribution can also account for a range of potential confounding factors. This is difficult to do using the exome, but it can be generated using non-genic regions, which contain fewer functional sites thus their evolution is primarily driven by neutral processes. In our case, we have access to previously published whole chr21 data from the same samples (*3*) and could use the non-genic regions of chr21 (non-genic-chr21) to generate our null distributions. Note that this method is conservative because it is likely that the non-genic regions of chr21 do contain some sites under natural selection. The chr21 and exome data are identical in a range of important ways, from the individuals included to the filtering process. Thus, using this empirical distribution also allows us to account for a range of potential confounding factors due to demographic history or study design. Data processing was also identical with the same site filtering criteria and near identical sample filtering. The only difference between these datasets is the efficiency of the target capture step, which resulted in slight differences in the sequence coverage distribution (with chr21 capture data generally having slightly higher coverage) (Fig. S23). The difference is not substantial (median coverage per filtered sample across the whole target space: 4.96× for the exome capture and 5.23× for the chr21 capture) and by estimating allele frequencies from genotype-likelihoods we minimise the effect of this coverage discrepancy. Indeed, comparing the allele frequency estimates for chr21 exonic SNPs present in both datasets (the filtered exome capture and filtered chr21 capture data) shows that allele frequency estimates are highly correlated across the two datasets, with no evidence for biases or important differences among them (Fig. S24 and S25).

In addition, we note that in our analyses we account for coverage when selecting candidates by binning both the exome and non-genic-chr21 data according to coverage (details below). This step controls for differences in coverage across exonic SNPs and for potential differences in coverage between the chr21 and exonic SNPs.

##### *6.3. Genetics-only test*

Classical selection analysis based on allele frequency alone relies on identifying sites with high allele frequency differentiation between populations compared to a neutral model (*189*). High genetic differentiation is thus considered evidence of divergent positive selection between populations. However, false positives can occur as different populations may have very different allele frequencies (and therefore high divergence) due to independent drift in the absence of panmixia. This can be accounted for by correcting for neutral genetic structure.

BayPass (*83*) is a selection scan method which accounts for neutral population structure by implementing an extension of the method of Bayenv2 (*190*). BayPass takes population allele counts as input. The conditional distribution of population allele frequency given allele counts is binomial assuming Hardy-Weinberg equilibrium. For the core model, the prior distribution of the vector describing population allele frequencies for site $i$ across $J$ populations ($\alpha_i^*$) is a multivariate gaussian (equation 1). $\alpha_{ij}^*$ is an instrumental variable which may take a value <0 or >1 and relates to an allele frequency on the real line such that $\alpha_{ij} = min(1, max(0, \alpha_{ij}^*))$. Here we refer to $\alpha_{ij}^*$ as 'unstandardised allele frequencies'. The ancestral population allele frequency ($\pi$) is the mean reference allele frequency weighted by a prior distribution estimated from the data. The precision matrix ($\Lambda$) is the inverse of the scaled covariance matrix of the population allele frequencies ($\Omega$). Parameters $\alpha_i^*$, $\Lambda$ and $\pi_i$ are sampled at time $t$ from the MCMC at $T$ different times.

BayPass calculates 'standardised allele frequencies' ($\ddot{\alpha}$) which account for population structure by multiplying unstandardised allele frequencies by the inverse of the Cholesky decomposition of $\Omega$ ($\Gamma_\Omega^{-1}$) (equation 2). The posterior means of the standardised allele frequencies ($\hat{\ddot{\alpha}}_i$) are rescaled using the mean ($\mu_{\ddot{\alpha}}$) and variance ($\sigma_{\ddot{\alpha}}$) (equation 3) and these values are used to calculate $X^tX^*$ (equation 4) (*84*). $X^tX^*$ is analogous to global $F_{ST}$ (*191*). Sites with exceptionally high $X^tX^*$ values have highly differentiated allele frequencies and are candidate targets of positive selection.

$$\boldsymbol{\alpha}_i^* | \Lambda, \pi_i \sim N_J\left(\pi_i 1_J; \pi_i(1-\pi_i)\Lambda^{-1}\right) \tag{1}$$

$$\hat{\ddot{\alpha}}_i = \Gamma_\Omega^{-1} \left\{ \frac{\alpha_{ij} - \pi_i}{\sqrt{\pi_i(1-\pi_i)}} \right\}_{(1..j)} \tag{2}$$

$$\hat{\ddot{\alpha}}_i^* = \left\{ \frac{\hat{\ddot{\alpha}}_{ij} - \mu_{\ddot{\alpha}}}{\sigma_{\ddot{\alpha}}} \right\}_{(1..j)} \tag{3}$$

$$\widehat{X^t X_i^*} = \hat{\ddot{\alpha}}_i^{*\,t} \hat{\ddot{\alpha}}_i^* \tag{4}$$

*6.3.1. Standardised allele frequencies*

If BayPass accounts for population structure in our data effectively then we should be unable to recover neutral population structure from the standardised allele frequencies. To test this, we performed various cluster analyses to investigate structure in the allele frequencies. We look for clusters by plotting correlation matrices, making hierarchical clustering trees based on the average agglomeration method and k-medoids clustering. k-medoids aims to find k clusters which minimise the sum of dissimilarities from cluster centres, while k-means uses the average of all the points in a cluster as the centre, k-medoids uses an actual data point. k-medoids clustering is more robust to outliers than k-means. The optimum average silhouette width can be used to select the best value for k and a Duda-Hart test can be used to estimate if clusters are significant. This is all implemented in the pamk() function from the fpc package (*192*). We also produced biplots in order to reduce dimensionality and visualise clusters in the first two principal components and in the loading vectors using prcomp() from the stats package (*193*).

Applying these methods to a random sample of 5,000 SNPs from the exome data does not recover neutral population structure such as separation into subspecies, indicating that BayPass effectively accounts for this potential confounding factor.

*6.3.2. Selecting genetics-only candidates*

Under the core model $X^tX^*$ values would be expected to follow a $\chi^2$ distribution with $J$ degrees of freedom ($\chi^2_J$). In practice, $X^tX^*$ values may not fit the $\chi^2_J$ distribution well due to violation of the assumption of normally distributed allele frequencies. We, therefore, opted to generate an empirical null distribution using results from the non-genic regions of chr21 (further justification for using this null is found in Supplemental Note 6.2). Contrary to expectations under local adaptation, fewer SNPs in the exome had very high allele frequency differentiation compared to the non-genic-chr21 null.

We found that there was an overrepresentation of SNPs with lower coverage in the tail of the distribution. There is no reason to expect that low-coverage sites would be more likely to underlie local adaptations, instead, this is likely due to the higher uncertainty in population allele frequency estimates leading to greater allele frequency variation. We, therefore, performed a *post hoc* coverage correction. Coverage was calculated as the total number of reads across all samples and populations at a site. This was calculated using the -doSnpStat 1 flag while estimating MAFs per population in ANGSD which reports the sequencing depth at each site for each population. A custom python script was used to sum sequencing depth over all populations in each subspecies-dataset. Sites were separated into five coverage bins, the bounds of the lowest and highest bins were selected so each contains 10% of the exonic SNPs. The remaining three bins were selected to have an equal width (but not necessarily contain an equal number of SNPs) to give five bins in total. False positive rates (FPRs) were estimated within each bin using only non-genic-chr21 SNPs within the same coverage bin as a null, we are able to do this because the coverage distribution of the exome and non-genic-chr21 data is similar albeit slightly shifted towards higher coverage in chr21-non-genic (Fig. S29). This results in more stringent thresholds in lower coverage bins (Fig. S28) meaning that the candidate SNPs are no longer biassed towards low coverage sites (Fig. S29).

##### *6.3.3. Candidate distribution in the genome*

We find that candidate SNPs cluster in the genome resulting in peaks which can be seen in Manhattan plots (Fig. S30) and a higher ratio of SNPs to genes in the tails compared to the background (Fig. S31). This clustering is due to linkage disequilibrium, although this is not evidence of selection in itself, it demonstrates that our results do not represent random noise caused by processes such as sequencing errors. The patchy nature of exome data and the likely prevalence of soft sweeps may explain why there is a relatively small amount of candidate clustering in the genome.

##### *6.3.4. Candidate gene overlap between subspecies-datasets*

Focusing on the genes that contain these candidate SNPs (candidate genes), there is more overlap between candidate genes in *All* and subspecies-specific datasets than between the subspecies-specific datasets (Fig. S32) suggesting within subspecies patterns are contributing to patterns detected across subspecies. Although there is generally a greater overlap of SNPs in *All* and subspecies-specific datasets than between subspecies-specific datasets, 92% of genes in any dataset are found in all in all datasets and the above pattern remains when restricting to only SNPs present in all datasets (n=18,851).

##### *6.3.5. Allele frequency patterns at genetics-only candidate SNPs*

To identify populations driving candidate SNPs and to ensure that neutral population structure was not driving our candidates, we tested for structure in the standardised population allele frequencies by plotting correlation matrices, making hierarchical clustering trees and performing k-medoids clustering at candidate SNPs (Fig. S33) as described above. We do not recover neutral population structure again indicating that BayPass effectively accounts for this potential confounding factor. Although we find a small amount of structure, we do not find evidence for particular populations driving candidates more than others.

To check that our candidates are highly differentiated, we calculated the pairwise population allele frequency differences for both unstandardised and standardised allele frequencies for each FPR tail and a random sample of SNPs. We see a clear shift towards larger allele frequencies in the candidates compared to the background which is more pronounced at more stringent tails (Fig. S34). This pattern becomes even clearer when focusing only on the maximum allele frequency differences per SNP with a large proportion of candidates having fixed or nearly fixed differences between populations (Fig. S35). These distributions suggest that our candidates have exceptionally large allele frequency differences between populations as expected under local adaptation.

##### *6.4. Genotype-environment association analysis*

Genotype-environment association analyses (GEA), also known as environmental association analyses (EAA), test for correlations between allele frequencies and environmental variables. These tests are based on the expectation that the frequency of an allele underlying adaptation to a particular selection pressure should correlate with the selection pressure. GEAs have two main advantages over genetics-only tests. Firstly, the inclusion of predictors increases the power to detect selection resulting in subtle changes in allele frequencies, for example, due to polygenic selection. Polygenic adaptation is likely to be important in chimpanzees as it is known to be important in the evolution of humans (*32*, *117*, *118*), the *Pan* clade's closest living relatives. Secondly, GEAs help to link potential selective pressures to genomic loci and allow specific hypotheses to be tested.

BayPass allows the integration of environmental variables to perform a GEA using a Bayesian hierarchical model to test for a linear relationship between standardised allele frequencies and environmental variables. BayPass offers two choices of alternative models: the standard (STD) and the auxiliary (AUX) models. The STD model (equation 5) extends the core model (equation 1) by introducing the vector of environmental variables (Z) of length J (number of populations) and correlation coefficient ($\beta_i$). The Bayes factor can then be computed using an importance sampling algorithm which samples MCMC runs under the core model. The AUX model (equation 6) also introduces the binary auxiliary variable ($\delta_i$) which can equal 1 or 0 indicating that a particular environmental variable is or is not associated with SNP i respectively. The posterior mean of $\delta_i$ can therefore be interpreted as the posterior probability of an association between a SNP and an environmental variable. The Bayes factor ($BF_{mc}$) can be computed from this model by multiplying the estimated posterior odds by the prior odds ($b_p/a_p$) (equation 7). We decided to use the AUX model for our analyses because it explicitly accounts for multiple testing by using prior odds which assume only a small proportion of the genome will be under selection (~1%).

$$\boldsymbol{\alpha}_i^* | \Lambda, \beta_i, \pi_i \sim N_J\left(\pi_i 1_J + \beta_i \mathbf{Z}; \pi_i(1-\pi_i)\Lambda^{-1}\right) \tag{5}$$

$$\boldsymbol{\alpha}_i^* | \Lambda, \beta_i, \delta_i, \pi_i \sim N_J\left(\pi_i 1_J + \delta_i \beta_i \mathbf{Z}; \pi_i(1-\pi_i)\Lambda^{-1}\right) \tag{6}$$

$$\mathrm{BF_{mc}} = \frac{\widehat{\mu(\delta_i)}}{1-\widehat{\mu(\delta_i)}} \frac{b_p}{a_p} \tag{7}$$

*6.4.1. Selecting GEA candidates*

Candidates were selected using the same method as for the genetics-only test; using results from chr21 as a null distribution to select candidates at BF thresholds corresponding to FPRs of 0.5%, 0.1% and 0.05% (justification for using this null is found above in Supplemental Note 6.2).

Similarly to the genetics-only analysis, we found that there was an overrepresentation of SNPs with lower coverage in the tail of the BF distribution likely due to the higher uncertainty in population allele frequency estimates leading to greater allele frequency variation. We, therefore, performed a *post hoc* coverage correction, using the same method as described for the genetics-only test (above). As seen in the genetics-only analysis, this generally results in more stringent thresholds in lower coverage bins (Fig. S38) meaning that the candidate SNPs are no longer biassed towards low coverage sites, instead, the candidate coverage distributions resemble the whole exome distribution (Fig. S39).

*6.4.2. Candidate distribution in the genome*

As in the genetics-only results, we find that GEA candidate SNPs cluster in the genome resulting in peaks which can be seen in Manhattan plots (Fig. S40) and a higher ratio of SNPs to genes in the tails compared to randomly sampled SNPs (Fig. S41) in *All* and *Central-Eastern* but not in *Western*. This clustering is due to linkage disequilibrium, although this is not evidence of selection in itself, it demonstrates that our results do not represent random noise caused by processes such as sequencing errors. The patchy nature of exome data and the likely prevalence of soft sweeps may explain why there is a relatively small amount of candidate clustering in the genome. The lack of evidence of selection in *Western* on the genome-scale may explain why we see little evidence of candidates clustering in the genome (Fig. S41).

*6.4.3. Candidate gene overlap between subspecies-datasets*

As found in the genetics-only test, there is more overlap between candidate genes in *All* and subspecies-specific datasets than between the subspecies-specific datasets (Fig. S43) suggesting within subspecies patterns contribute to patterns detected across subspecies. This pattern remains when accounting for the fact that there is generally a greater overlap of SNPs in *All* and subspecies-specific datasets than between subspecies-specific datasets.

*6.4.4. Allele frequency patterns at GEA candidate SNPs*

As for the genetics-only analysis, we tested for structure in the standardised population allele frequencies (calculated under the core model) at candidate SNPs. Unlike the genetics-only candidates, the GEA candidate SNPs show clear structure. This structure does not correspond to neutral population structure but instead to the habitat covariable (forest-tree-percentage) demonstrating that BayPass effectively accounts for neutral population structure and identifies SNPs associated with the habitat covariable (Fig. S44).

To independently verify that GEA candidate SNPs correlate with the habitat covariable, we calculated the Pearson correlation coefficient (r) of allele frequency (either standardised or unstandardised calculated under the core model) against forest-tree-percentage for savannah and forest candidate SNPs separately. When all candidates are analysed together, using either standardised or unstandardised allele frequencies, correlations are highly significant ($p<10^{-62}$) (mostly due to the large number of SNPs), r always has the correct sign and the absolute value of r increases at more stringent tails (Fig. S45). When correlations are tested for each SNP independently, p-values are not always significant due to the small number of sample points, however, the distribution of p-values shifts further towards 0 at more stringent tails (Fig. S46). The value of r always has the correct sign, with the exception of a few outliers in *All* and two *Central-Eastern* candidates, and the distribution of r values shifts further from 0 at more stringent tails (Fig. S46). These results independently confirm that BayPass effectively identifies candidate SNPs which correlate with the habitat covariable.

Local adaptation is likely to be mostly polygenic and driven by soft sweeps on standing genetic variation, resulting in moderate allele frequency differences among populations, as observed in humans (*32*, *117*, *118*). Our study design and stringent SNP filtering criteria also mean that many selected SNPs may not be included in the final dataset, with the evidence of local adaptation on these SNPs identified through signatures at linked variants that are more likely to show subtle allele frequency changes. Indeed, the GEA candidate SNPs differ consistently in allele frequency with respect to habitat type but do not necessarily have large frequency differences between populations (Fig. S45A).

##### *6.4.5. Accounting for population structure*

In our dataset, a general pattern of isolation by distance in chimpanzees (*3*) and environmental spatial autocorrelation results in a correlation between ancestry and habitat within subspecies, such population structure can lead to spurious signals of selection if not properly accounted for. BayPass has been used effectively on a range of biological systems (e.g. (*84*, *188*, *194–198*)) and has been shown to account for population structure under a range of simulated demographic histories (*83*). In our analyses, the estimated covariance matrices agree with previous population structure analyses (*3*) and the standardised allele frequencies showed no population structure overall indicating that BayPass does indeed correct for neutral population structure in our data. We further ensure against the confounding effect of population structure by using non-genic-chr21 (which has a near identical demographic history to the exome) to generate empirical null distributions.

##### *6.4.6. Issa Valley*

Issa Valley is the only savannah-like population in *Central-Eastern* with a forest-tree-percentage of 22.72%, the next lowest value in *Central-Eastern* is Chinko with 75.36% (Fig. 3). This makes Issa Valley a valuable data point representing a savannah-like habitat in the central-eastern clade.

To determine to what extent Issa Valley drives the *Central-Eastern* candidate SNPs, we ran BayPass with Issa Valley removed. Fig. S47 plots the distribution of BFs from this analysis showing the distribution for all SNPs and for SNPs identified as candidates in the full *Central-Eastern* analysis (i.e. with Issa Valley included). The candidate distributions are clearly shifted to lower FPRs and higher BF values in the analysis excluding Issa Valley also. This effect becomes more pronounced at more stringent tails indicating that Issa Valley alone does not drive the *Central-Eastern* GEA candidates. Unsurprisingly, this shift is far less pronounced for savannah candidates than forest candidates. Thus, signatures of selection identified in the *Central-Eastern* GEA candidates are generally not driven by Issa Valley alone, however, many savannah candidates identified in *Central-Eastern* may represent positive selection in this single population.

#### 7. Gene set enrichment

Genomic analyses of chimpanzees benefit from the detailed functional annotation of the human genome which can be confidently translated to chimpanzees due to low divergence between the species. The functions of specific genic or regulatory candidate SNPs can be inferred from the genome annotation and gene set enrichment analysis can be applied to test which biological traits are overrepresented in the candidate loci. Gowinda (*149*) is a program which performs gene set enrichment while accounting for potential confounding factors such as gene length and overlapping genes. Gowinda works by randomly sampling SNPs from all SNPs in the analysis and records the overlapping genes according to a user-provided genome annotation file. Gowinda can be run in ‘gene’ mode where multiple SNPs in the same gene are counted once (this is based on the assumption that SNPs within genes are in complete linkage) or in ‘SNP’ mode where multiple SNPs in a gene are counted multiple times (this relies on the assumption that SNPs are in linkage equilibrium). Running in ‘gene’ mode is advisable as it partially accounts for linkage disequilibrium. In ‘gene’ mode, SNPs are randomly sampled until the same number of genes are sampled as overlap with the candidate SNPs. Empirical p values for each gene set are calculated as the proportion of resampling runs which contain the same number of genes or more than observed in the candidate list. Multiple testing is accounted for by calculating the false discovery rate (FDR) for each p by dividing the number of gene sets expected to have a p-value $\leq$ p by the number observed to have a p-value $\leq$ p.

We tested for enrichment of hypothesis-free datasets and hypothesis-driven datasets. The hypothesis-free datasets tested were Gene Ontology (GO) categories (*150*), KEGG pathways (*151*), Reactome categories (*152*), human GWAS traits (*153*), phenotype database traits (*154*) and tissue expression data from the Human Protein Atlas (*155*) where genes were considered associated with a tissue if expression level was ‘high’ and reliability was ‘approved’.

Pathogens, particularly SIVcpz, are major drivers of differential adaptations among the four subspecies of chimpanzees (*65*, *66*) and there is local variation in the prevalence of pathogens within subspecies (*14*, *92*, *199–201*). To test whether SIV and other pathogens also drive local adaptation within subspecies, we tested for enrichment of pathogen- and immunity-related genes among the candidate SNPs. We tested for enrichment of two lists of immunity genes, the first being a manually curated list of 356 genes associated with immune function from (*93*) and the second being a manually curated list of 1,553 innate immunity genes from (*94*). Manually curated lists of genes that encode host proteins which physically interact with viral proteins, viral DNA or viral RNA, known as viral interacting proteins (VIPs), were also tested (*156*, *157*).

We also tested for enrichment of lists of genes related to a variety of pathogens similar to those infecting wild chimpanzees, we collectively refer to these as 'other pathogen-related genes'. SIV response genes are those which are differentially expressed between experimentally infected natural (vervet monkeys) and naïve (macaques) host CD4+ T lymphocytes (*158–160*). Malaria genes consist of a manually curated list of genes associated with malaria used to identify polygenic adaptation in humans (*96*), 23 red blood cell genes with strong links to malaria in the literature collated in (*97*) and 295 conserved mammalian genes (thus excluding *HBB*, *HBD*, *GYPA* or *GYPB*) related to *P. reichenowi*, *P. vivax* or *P. falciparum* (close relatives of parasites infecting wild chimpanzees (*57*)) identified in the literature by (*98*). Influenza-related genes are those which show significantly different expression between infected and control trials with biologically significant effect sizes in a meta-analysis of 18 studies (*161*). SARS-Cov-2 GWAS genes are those associated with SARS-Cov-2 in the COVID-19 Host Genetics Initiative (round 4 alpha September 30, 2020) (*202*), SARS-Cov-2 interacting genes are those which encode proteins which physically interact with SARS-Cov-2 identified in (*163*, *165*), and genes which are differentially expressed upon infection with SARS-Cov-2 are those identified in (*164*, *203*) and genes with $p<0.05$ in the differential expression analysis done by (*162*). Anthrax-related genes are the top 25 upregulated genes in human cells 4 hours and 24 hours (49 unique genes in total) after exposure to *Bacillus anthracis* spores (*168*), and 17 genes reported to be associated with anthrax using MalaCards (www.malacards.org) (*204*). Ebola-related genes are those identified as being related to ebola in an extensive literature search (*169*). Herpes Simplex Virus 1 (HSV-1) associated genes are those which were significantly upregulated in infected cells and showed viral gene expression compared to cells which were not exposed to the virus and cells which were exposed but did not show any viral expression (*167*).

We also hypothesised that dehydration stress may drive adaptation in savannah habitats (*30*) and so tested for enrichment of 'dehydration response genes'. Dehydration response genes are those which are significantly differentially expressed between dehydrated and hydrated Cactus mice (a desert-adapted species) (*89*) and 133 'classical water conservation pathway' genes collated in (*90*).

##### *7.1. Genetics-only candidates*

Although the genetics-only test did not find evidence of selection on the genome-scale, this does not exclude the possibility that some exonic SNPs have evolved under local adaptation, and the most highly differentiated SNPs in the exome (which are not driven by population substructure, Fig. S33) are the best candidates. To identify potential selection pressures driving selection at these sites, we tested for enrichment of the gene categories described above.

The hypothesis-free enrichment analysis found a significant enrichment (FDR<0.05) of lung-related GWAS categories in the *All* 0.1% tail (Airway Obstruction, FDR=0.001; Tobacco Use Disorder, FDR=0.018; Bronchodilator Agents, FDR=0.044; Forced Expiratory Volume, FDR=0.044) driven by 6 candidate genes (Fig. S48). No enrichment for these categories was observed in the subspecies-specific databases, however, standardised allele frequencies show that subspecies differences do not drive these candidates in *All*. Convergent evolution could instead explain this pattern, however, there is no clear consistent trend in population allele frequencies in these genes which would help identify potential selection pressures. Significant enrichment of GWAS genes related to the aorta is also observed in the *Western* 0.05% tail (FDR=0.046) driven by only two candidate genes. There is also a significant enrichment of genes highly expressed in the spleen in the 0.1% and 0.05% *Western* tails (FDR=0.023 and FDR=0.048) driven by eight and six candidate genes, respectively. Without clear prior hypotheses, it is difficult to speculate on the selection pressures which might drive selection for lung or aorta-related traits, however, the spleen plays an important role in immune system functioning and so it is possible that pathogen-mediated selection may explain this signal.

The hypothesis-driven enrichment tests revealed some limited evidence for pathogen-mediated selection. General immunity-related genes (*93*) are significantly enriched in the 0.05% *Nigeria-Cameroon* tail (FDR=0.035) and genes which change expression on SARS-CoV-2 infection (*162*, *164*, *203*) are nominally enriched (FDR=0.065) in the 0.05% *Western* tail (Fig. S48). There is generally a greater signal of enrichment for pathogen-related genes in *Western*. Nominal enrichment of genes related to SARS-CoV-2 does not reflect adaptation to this virus (which arose years after sampling) but may represent similar viruses. Wild chimpanzees are known to suffer from ‘flu-like’ diseases caused by respiratory viruses such as coronaviruses (*26*, *27*, *205–209*) although some of these outbreaks will be the result of exposure to humans in the very recent past (*121*, *140*). Viruses, particularly RNA viruses such as coronaviruses, are known to be strong selection pressures driving local adaptation in humans (*157*, *210–212*). Overall, the genetics-only analysis provides some evidence for viruses driving adaptation in the recent past as they are known to have been over longer time scales in chimpanzee evolution (*65*, *66*).

##### *7.2. GEA candidates*

*7.2.1. The effect of montane ecosystems on Central-Eastern forest candidates*

In the forest candidates, there is generally a greater enrichment for pathogen-related genes in *All* and *Western*, however, we do not find a similar pattern in *Central-Eastern.* Differences between montane and lowland ecosystems likely explain this pattern. *Central-Eastern* includes five eastern populations, Budongo, Bwindi, Gishwati, Ngogo and Nyungwe, which inhabit Albertine Rift montane forests at an elevation >1,000m, resulting in lower mean annual temperatures and therefore lower levels of vector-borne diseases such as malaria compared to lowland forests (*14*). Bwindi, Gishwati and Nyungwe in particular are at >2,000m elevation resulting in mean annual temperatures lower than 17°C which is over 3°C lower than any other population and over 6°C lower than any non-eastern population in this study. As mentioned above, Issa Valley is the only representative of a savannah-like habitat in *Central-Eastern*, this population also lies in the Albertine Rift with an elevation of 1494m resulting in a mean annual temperature of 20.4°C. This means that while there is a negative correlation between forest-tree-percentage and mean annual temperature for *All* (Pearson's $r = -0.24$, $p = 0.185$) and *Western* (Pearson's $r = -0.40$, $p = 0.159$), there is a positive correlation in *Central-Eastern* (Pearson's $r = 0.43$, $p = 0.125$). Although mean annual temperature represents only one of a myriad of possible selection pressures driving adaptation to habitat type, it indicates that the range of elevations represented in *Central-Eastern* may explain why candidate gene functions differ from the other subspecies-datasets.

*7.2.2. Viral-interacting proteins (VIPs)*

Although we find evidence of local adaptation to pathogens, we observed no significant enrichment of viral interacting protein (VIP) categories. This is initially surprising given that VIPs are known to be under strong natural selection across mammals (*156*), in humans (*157*, *210*), and in chimpanzees (*65*, *66*). It is possible that, as proposed for SIV (*65*, *66*) selection pressures on VIPs are strongest immediately after initial exposure to a novel virus while subsequent adaptation is driven by changes in other genes such as immunity genes. Our analysis investigates selection in the very recent past which may not encompass the emergence of novel geographically restricted pathogenic viruses that also had time to lead to host adaptation.

#### 8. Malaria candidates

##### *8.1. GYPA structural variants*

Considering that the glycophorin gene cluster is a well-established hotspot of structural variation in humans (*104*, *213*) and other great apes (*112*), we searched for associations between candidate alleles (A>T at hg19: chr4:145,039,806/hg38: chr4:144,118,653; and C>A at hg19: chr4:145,040,845/hg38: chr4:144,119,692) and gene copy number and/or linkage with nearby structural variants. We first examined this locus in genome assemblies, comparing the human reference genome to the chimpanzee reference genome panTro6/Clint_PTRv2 (*114*), a recently published chimpanzee diploid assembly AG18354 (*214*), and a new chimpanzee assembly made from previously published ONT reads from AG18359 (*113*), all derived from western chimpanzees (Fig. S53-A). After visual comparison between chimpanzee contigs and the human reference build hg38, we noticed that none of the assemblies carried the derived allele at the candidate alleles (Fig. S53-B), which can be explained by either true absence of the SNPs in the donor individuals or collapsed/missing gene copies in the assembly. To better assess assembly errors, we directly examined the total number of glycophorin gene copies by lifting over human gene annotations. Both the human reference genomes hg19 and hg38 contain three gene family members in chromosome 4q31.21, *GYPE*, *GYPB* and *GYPA* (Fig. S53-A). Similarly, panTro6 displayed three copies in synteny. The AG18354 diploid assembly differed, with both haplotypes showing an additional copy of *GYPE*, and one haplotype also showing an additional copy of *GYPB*. The new AG18359 assembly also contained an additional copy of *GYPE*. In contrast, previously published experimental copy-number assays in a chimpanzee individual using fibre-FISH supported the presence of three *GYPE* genes (*112*), suggesting that this region might be incorrectly assembled, likely due to technological limitations reconstructing structurally variant loci.

To overcome the limitations of the assemblies, we directly investigated three chimpanzee individuals for which long-read sequencing data exists, including PacBio CCS long-reads from Clint (the main donor of panTro6) (*114*), PacBio high-fidelity long-reads from AG18354 (*214*), and ONT reads from AG18359 (*113*). SNP genotyping from Illumina high-coverage data showed Clint as homozygous reference (C/C), while AG18354 and AG18359 as heterozygous (C/A) for the C>A substitution at chr4:145,040,845 in hg19. Inspection of Clint long-read data mapping to *GYPA* locus revealed only a small proportion of reads carrying the A allele (0.03 allele ratio) in line with the expected sequencing error rate of PacBio CCS reads. AG18354 and AG18359 were both heterozygous (albeit with skewed allele ratios) at the candidate SNPs sites (Fig. S53-C). Further examination of the reads mapping to *GYPA* locus in these two individuals, showed the presence of distinct molecularly phased haplotypes mapping to this locus, including one carrying a 16 kbp deletion ablating the last two exons of *GYPA* with a lower mapping quality score likely representing an alternative paralog, as well as one spanning full-length *GYPA* exclusively harbouring the candidate SNPs (Fig. S53-C). Clint, on the other hand, lacked reads carrying the 16 kbp deletion altogether suggesting this paralog does not exist, in line with its lower copy number.

Considering that chimpanzee population-level long-read sequencing data remains limited, we leveraged previously published short-read sequencing chimpanzee data (n=60) (*5*, *62*) to refine long-read sequencing findings. We estimated the copy number of all members of the glycophorin gene cluster using fastCN (*172*), which calculates copy number across 1 kbp windows based on read-depth of multi-mapping reads, thus providing gene-family copy-number estimates (Fig.

S53-D). We observed that windows overlapping glycophorins A-B-C in Clint displayed copy numbers >8, corroborating collapsed/missing copies in panTro6. To examine the association between the candidate SNPs and copy-number variation, we zoomed in on the *GYPA* locus (Fig. S53-E). In the human reference genome, *GYPA* comprises a duplicated portion, demarcated by a segmental duplication overlapping the first five exons, and a unique region, containing exons six and seven. To reduce methodological noise, we genotyped the copy numbers of the duplicated and unique portions of *GYPA* as the median copy number across 1kbp windows (Fig. S53-F). We corroborated the unique portion of *GYPA* as diploid copy-number two, with only nine individuals showing larger deletion/duplication events spanning the unique space downstream *GYPA*. The duplicated portion of *GYPA* showed copy-number polymorphism, with most individuals carrying eight to nine gene copies of the glycophorin genes. To detect possible SNP association with copy-number variation, we genotyped the candidate SNPs in the short-read sequencing data. We found only two individuals carrying the A>T substitution at chr4:145,039,806 and 17 carrying the C>A substitution at chr4:145,040,845 in hg19 coordinates. Focusing on the latter, we did not find significant associations between copy-number variation and the presence of the derived allele (Fig. S53-E), suggesting that this candidate SNP is tagging the ancestral *GYPA* full-length version independently of copy-number variation.

Both SNPs are present in the long-read data, confirming them to be true polymorphisms, and their allele frequencies in the high-coverage short-read data correspond to those in the PanAf exomes (Fig. S55). Although skewed allele balance in *GYPA* may result in an underestimation of derived allele frequencies, read depth does not correlate with habitat type (Fig. S54) or allele frequencies in the PanAf exomes at this locus, suggesting that copy number variation does not explain the strong genotype-environment association. We note that copy number variation is virtually impossible to study in genetic data from non-invasive samples, therefore, resolving this interesting locus will require further work.

##### ***8.2. Overlapping genes***

One of the Western forest candidate SNPs lies within HBD less than 5kb from HBB and thus was assigned to both HBB and HBD (both genes belong to the 'Malaria' gene set and HBB belongs to the 'Malaria (erythrocyte genes)' gene set), and two candidate SNPs lie within the GYPA and GYPB coordinates (both genes are in the 'Malaria' and 'Malaria (erythrocyte genes)'). Although gowinda is designed to account for overlapping genes and has been shown to do this effectively (*149*), for extra prudence, we checked whether the signal of enrichment for malaria-related categories in the western forest candidates remained if one of the two overlapping genes is removed from the enrichment analysis i.e. we re-ran the analysis assigning the HBB/HBD and GYPA/GYPB candidate SNPs to only one gene each. This was done by simply removing one of the overlapping genes from the annotation file passed to gowinda. Although removing genes removes significant enrichment (FDR<0.05) for any category, malaria categories still consistently show the strongest signal of enrichment in the pathogen response dataset at every tail and gene combination (Table S3).

We note that GYPA and GYPB appear as neighbouring in the Havana, NCBI and Ensembl annotations of hg38 (release 110) but their models overlap in the Ensembl hg19 annotation due to a single outlying GYPB exon. This is likely an annotation error in Ensembl. Re-running the analysis with this exon removed from the annotation still results in significant enrichment (FDR<0.05) of malaria-related genes (*96*) in the 0.5% Western tail and one of the three malaria categories has the strongest signal of enrichment in all tails (Table S4).

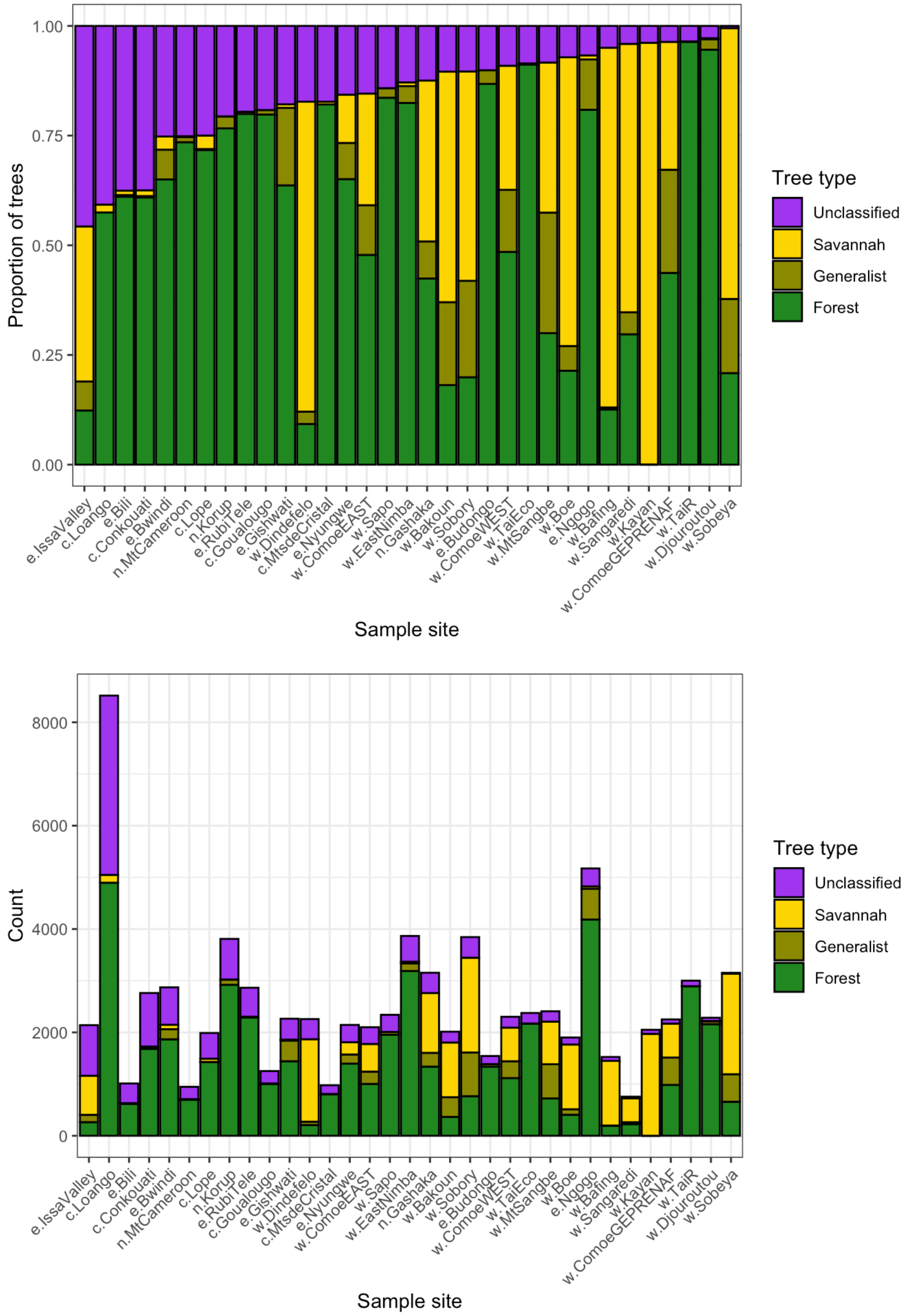


**Fig. S1. Data used to calculate forest-tree-percentage.** Top: The proportion of trees in each sample site classified as being a forest specialist, generalist, savannah specialist or unclassified according to (*87*). Bottom: the number of trees belonging to each category per sample site.

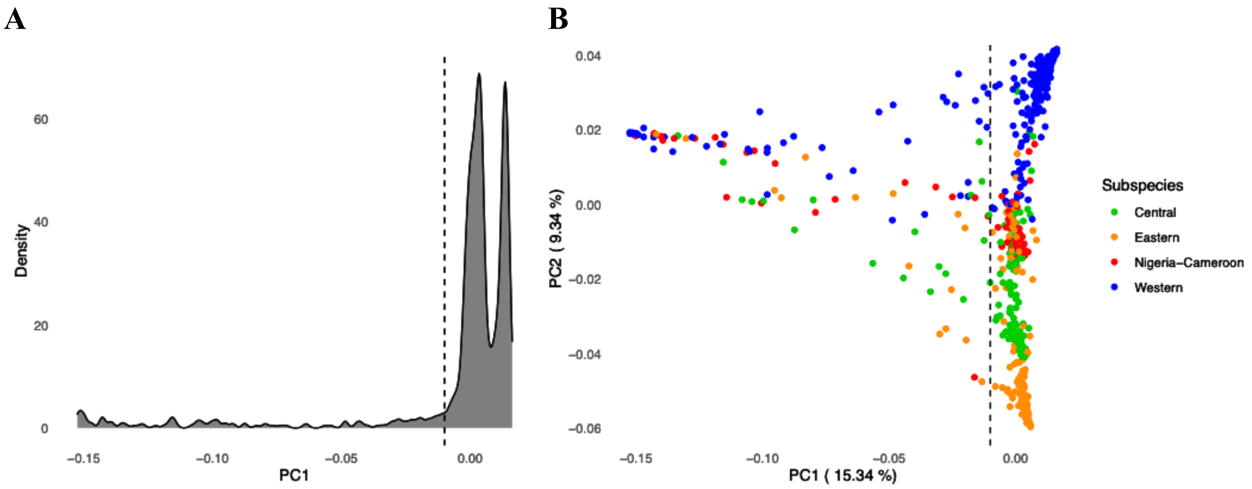


**Fig. S2. PCA of all samples (unfiltered exomes). A** Density of samples across PC1. The threshold of -0.01 (dotted line) was chosen as the point where the density of samples rises sharply. **B** PCA plot with PC1 and PC2 values for each sample coloured by subspecies and the threshold of -0.01 (dotted line). This plot reveals that the two peaks observed in A correspond to western and non-western samples.

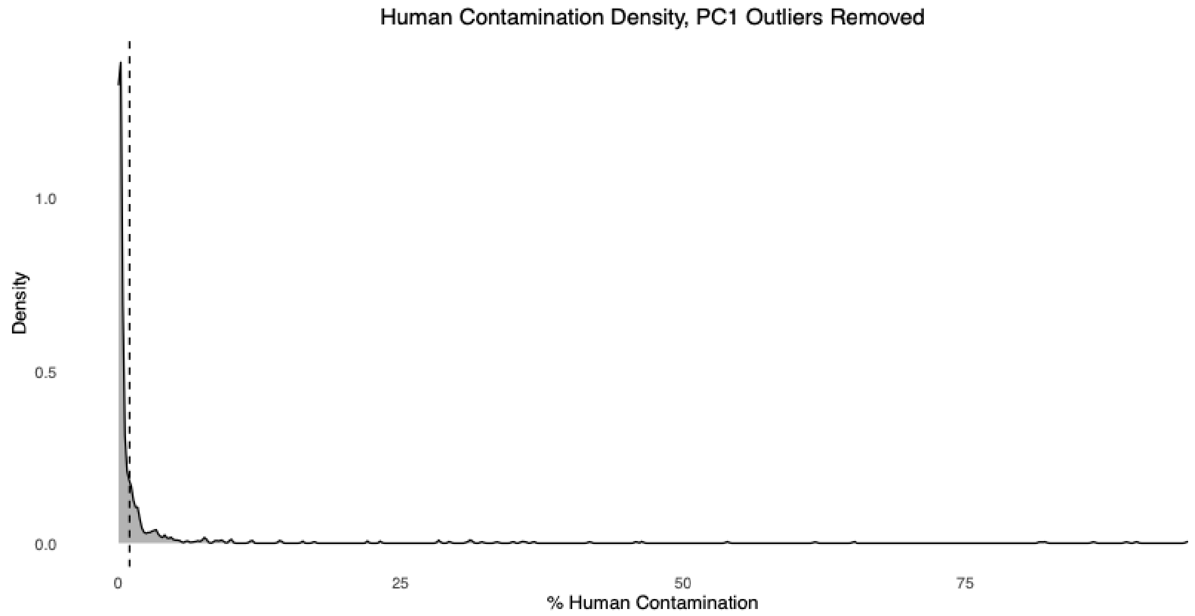


**Fig. S3. Distribution of human contamination estimates from HuConTest (*144*) which passed the PC1 filter (exome).** The 1% threshold is shown as a dotted line. The distribution is heavily skewed towards very low levels of human contamination.

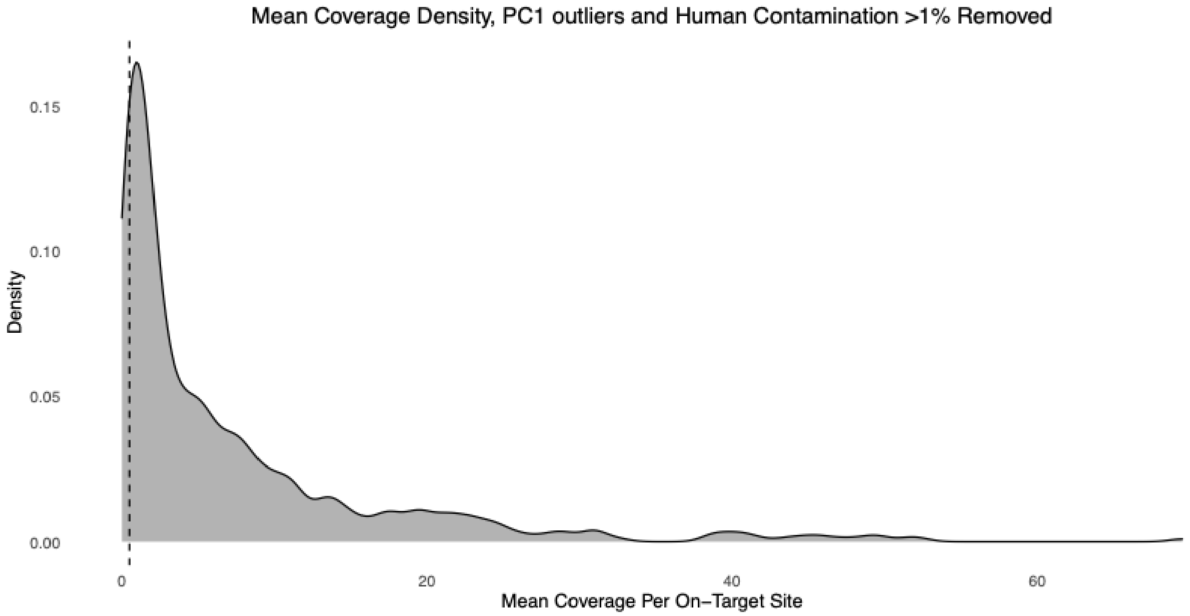


**Fig. S4. Density distribution of coverage for samples which passed contamination filtering (exome).** 0.5x threshold is shown by the dotted line. All samples below this threshold were filtered out.

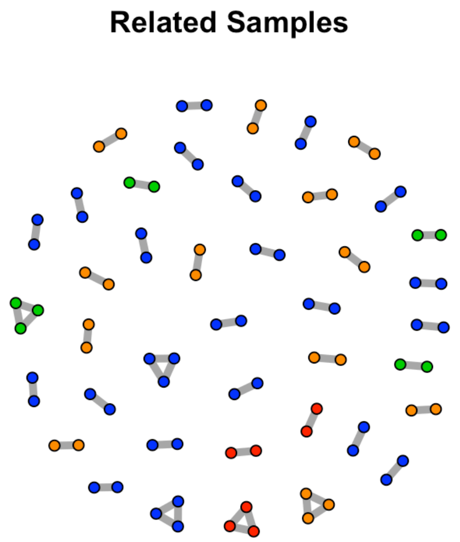


**Fig. S5. Graph showing samples (nodes) linked to related samples by edges for relationships where θ>0.1875.** Nodes are coloured according to subspecies (green=central, orange=eastern, red=Nigeria-Cameroon, blue=western).

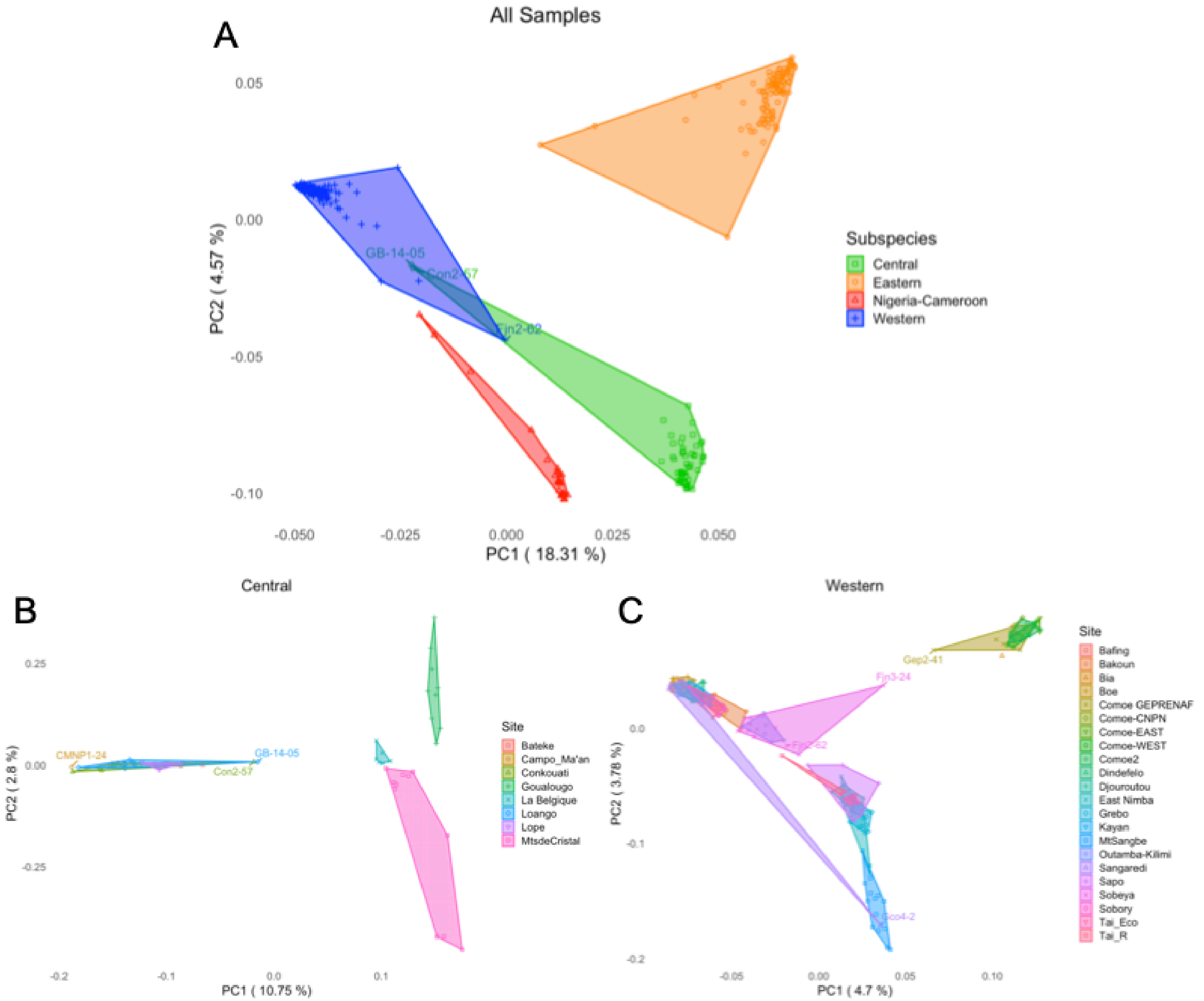


**Fig. S6. PCA plots generated from exomes for all samples (A), central samples (B) and western samples (C) with outlier samples labelled.** Polygons group samples labelled as the same subspecies (A) or sample site (B and C). Outlier samples are labelled.

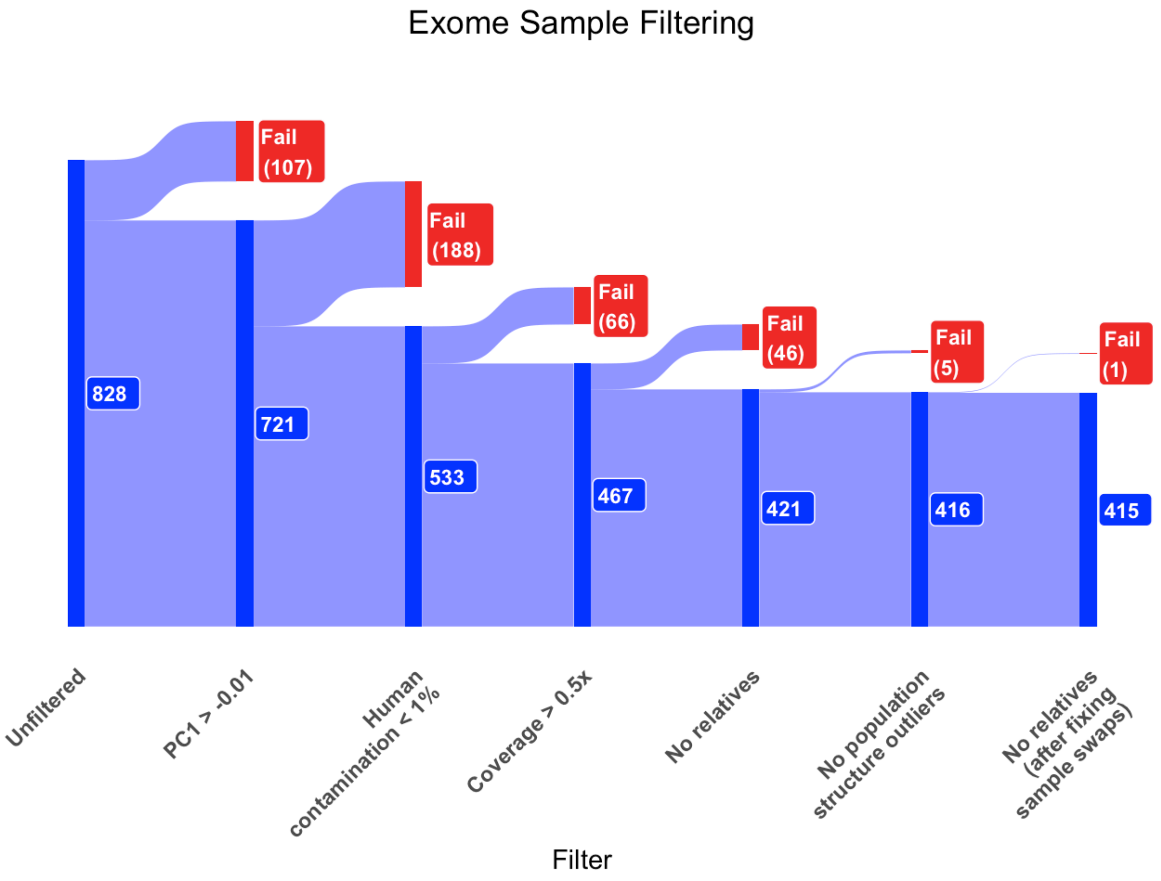


**Fig. S7. Sankey diagram showing the proportion of samples which pass or fail each stage of exome filtering.**

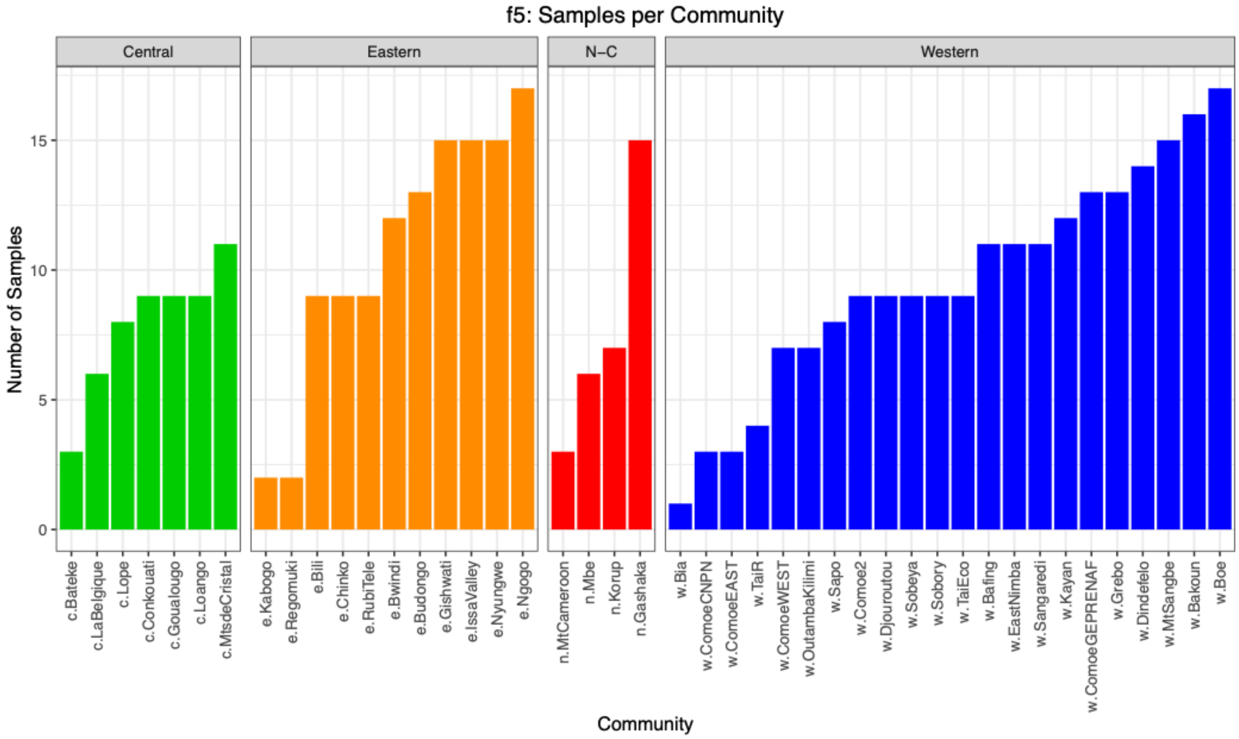


**Fig. S8. Number of samples per sample site after filtering (exome).**

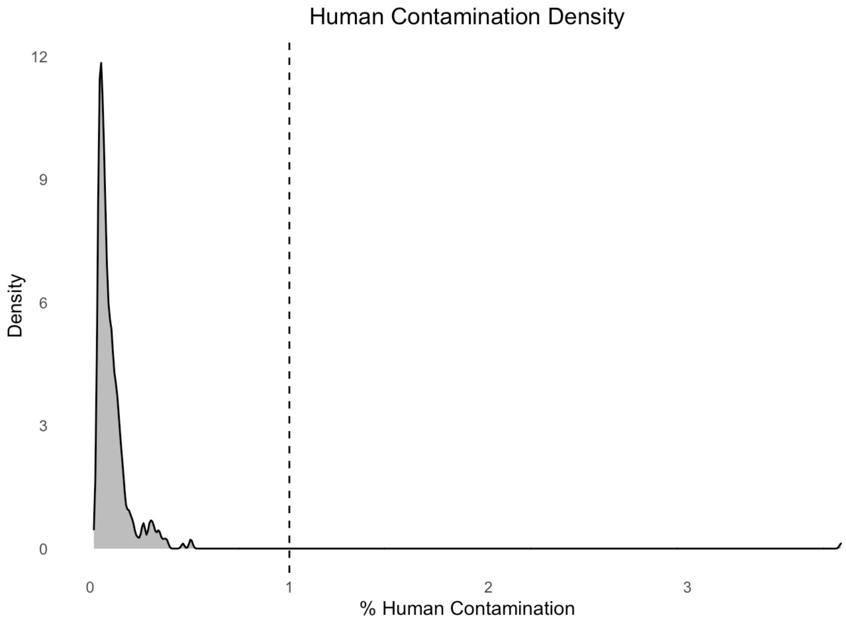


**Fig. S9. Distribution of human contamination estimates from HuConTest (*144*) for chr21 data from samples which passed filtering (exome).** The vertical dotted line represents a value of 1%, a single sample exceeds this threshold and was therefore removed.

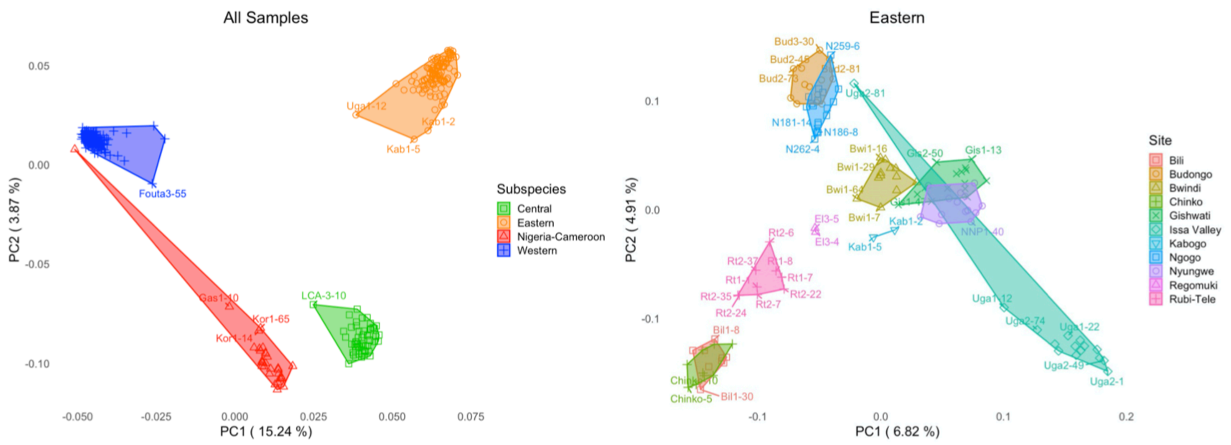


**Fig. S10. PCA plots generated from chr21 for all samples (left) and eastern samples (right) with outlier samples labelled.** Polygons group samples labelled as the same subspecies (when all samples are analysed) or sample site (when only the eastern subspecies is analysed separately). Left: the Nigeria-Cameroon outlier can be seen clustering near westerns. Right: The Issa Valley outlier can be seen clustering near Ngogo and Budongo. Both these outliers were filtered out of the chr21 data. All other samples cluster correctly.

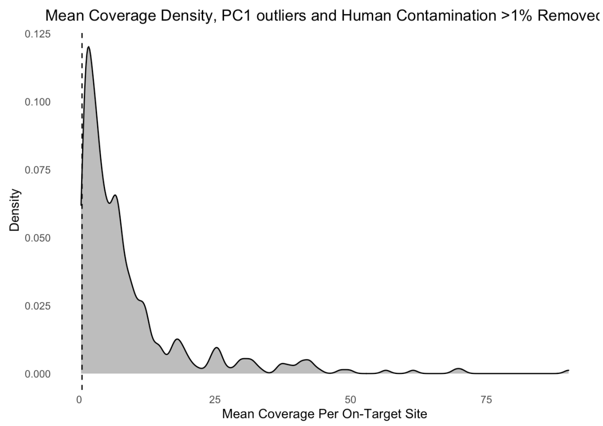


**Fig. S11. Coverage distribution of data after filtering (chr21).** The vertical dotted line represents coverage of 0.5x. Four samples have coverage < 0.5x but were not removed.

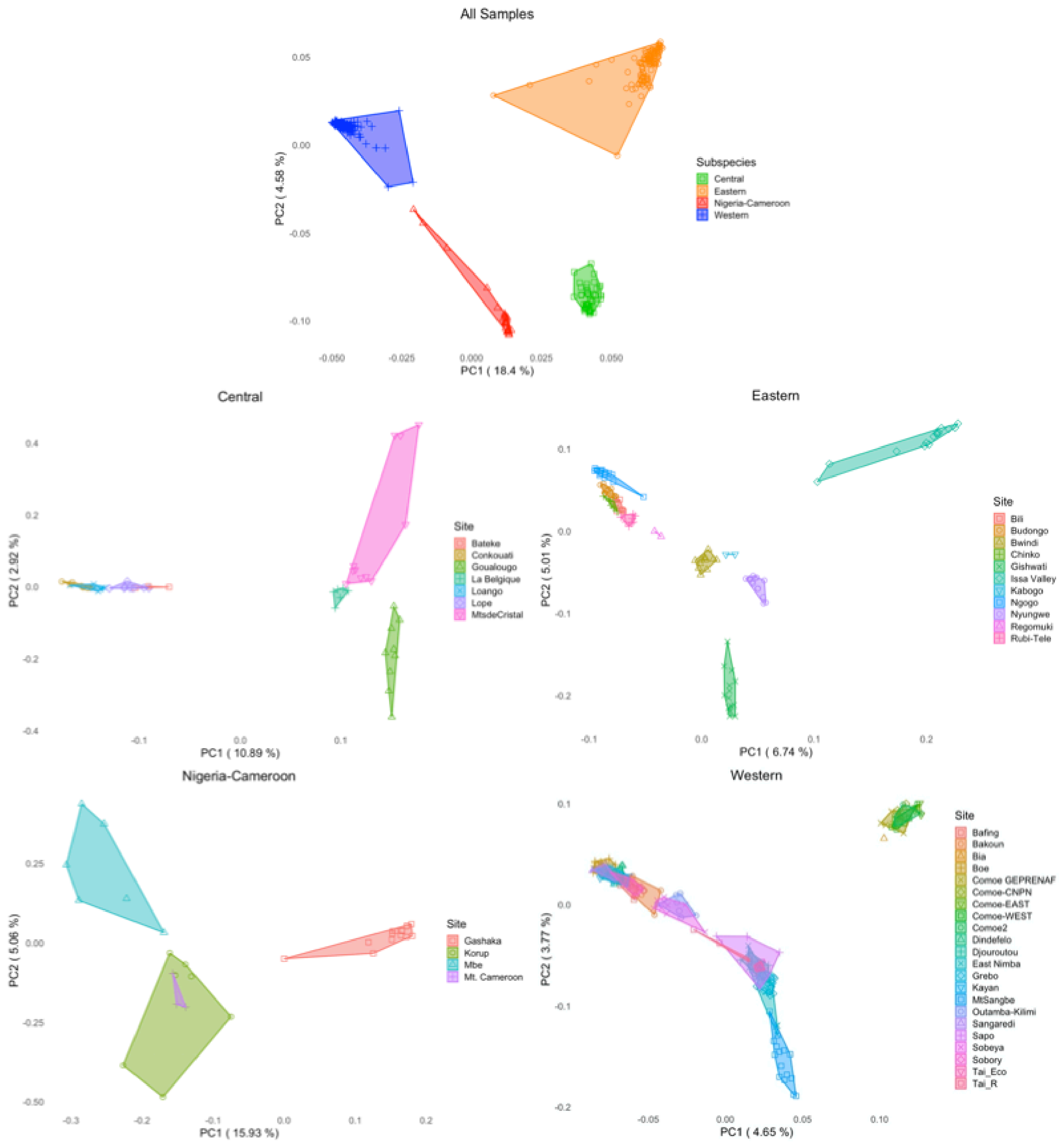


**Fig. S12. PCAs from running PCAngsd on the filtered exome data for all samples together and each subspecies separately.** Each subspecies is plotted separately to increase the resolution of the analysis. Points represent individuals, polygons represent subspecies in the all samples plot and sample sites in the subspecies-specific plots.

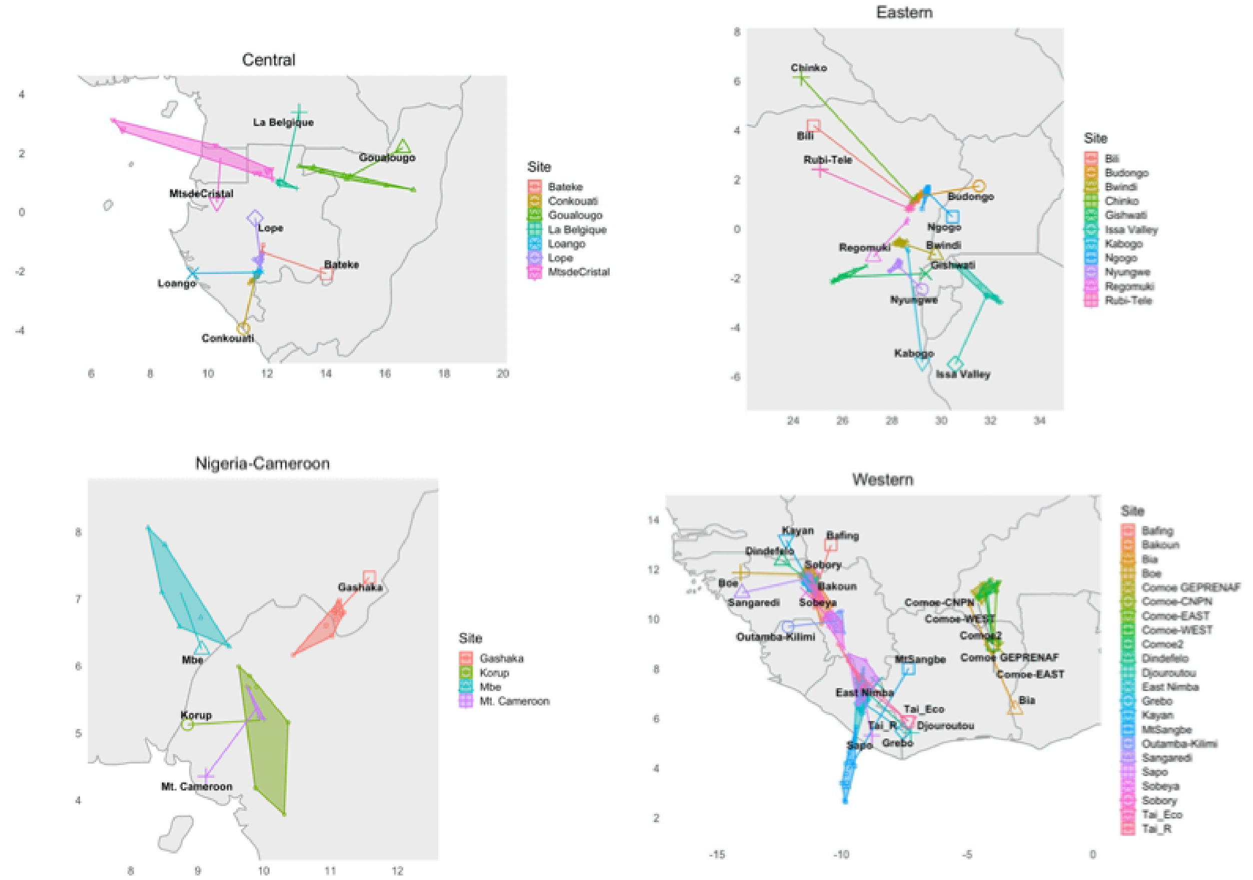


**Fig. S13. PCAs from PCAngsd on the filtered exome data for each subspecies separately Procrustes transformed onto a map.** Points represent individuals, polygons represent sample sites and lines link the centre of polygons to the geographical location of that sample site on the map.

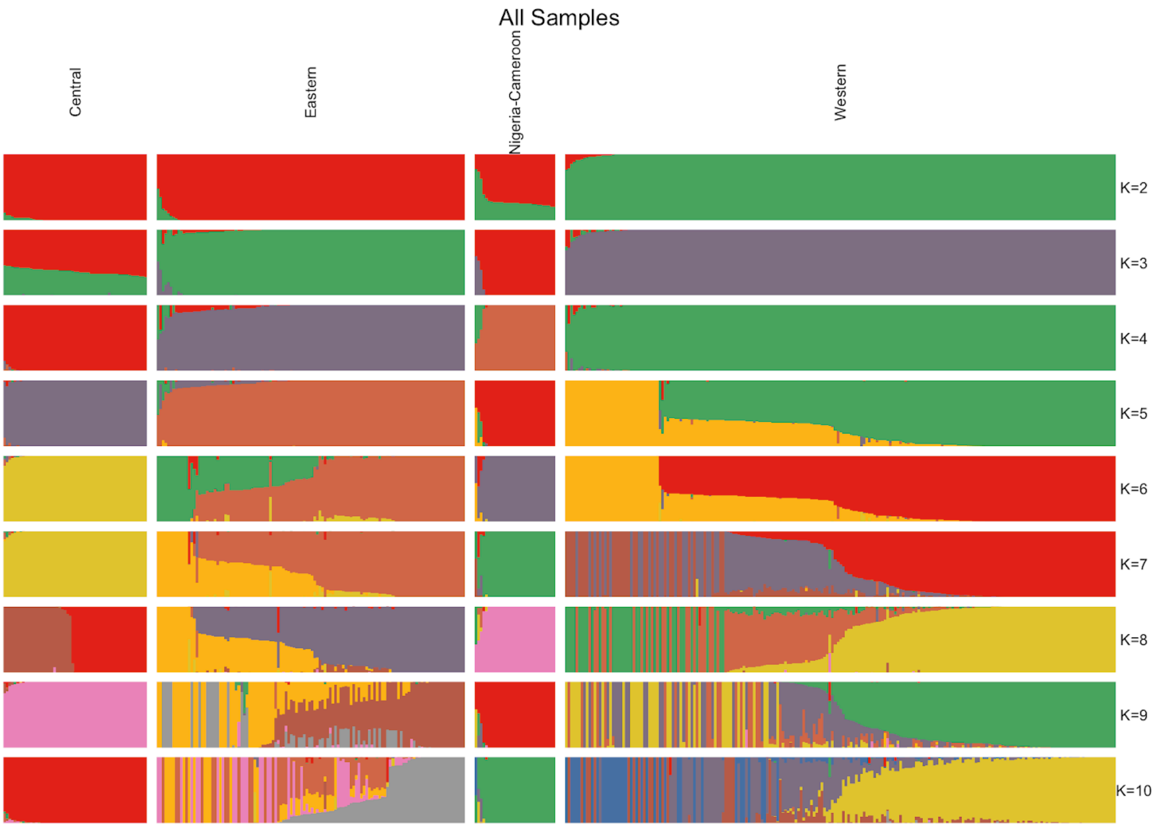
All Samples
Central
Eastern
Nigeria-Cameroon
Western
K=2
K=3
K=4
K=5
K=6
K=7
K=8
K=9
K=10

1

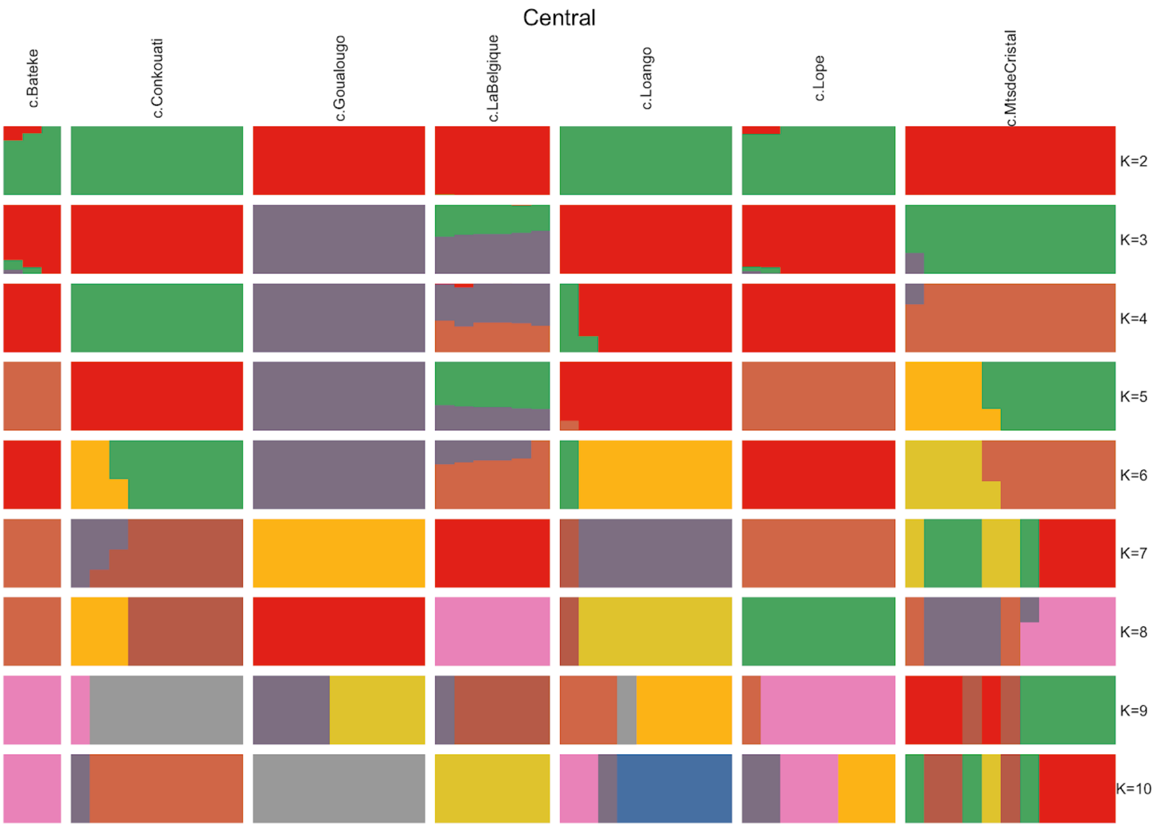
Central
c.Bateke
c.Conkouati
c.Goualougo
c.LaBelgique
c.Loango
c.Lope
c.MtsdeCristal
K=2
K=3
K=4
K=5
K=6
K=7
K=8
K=9
K=10

1

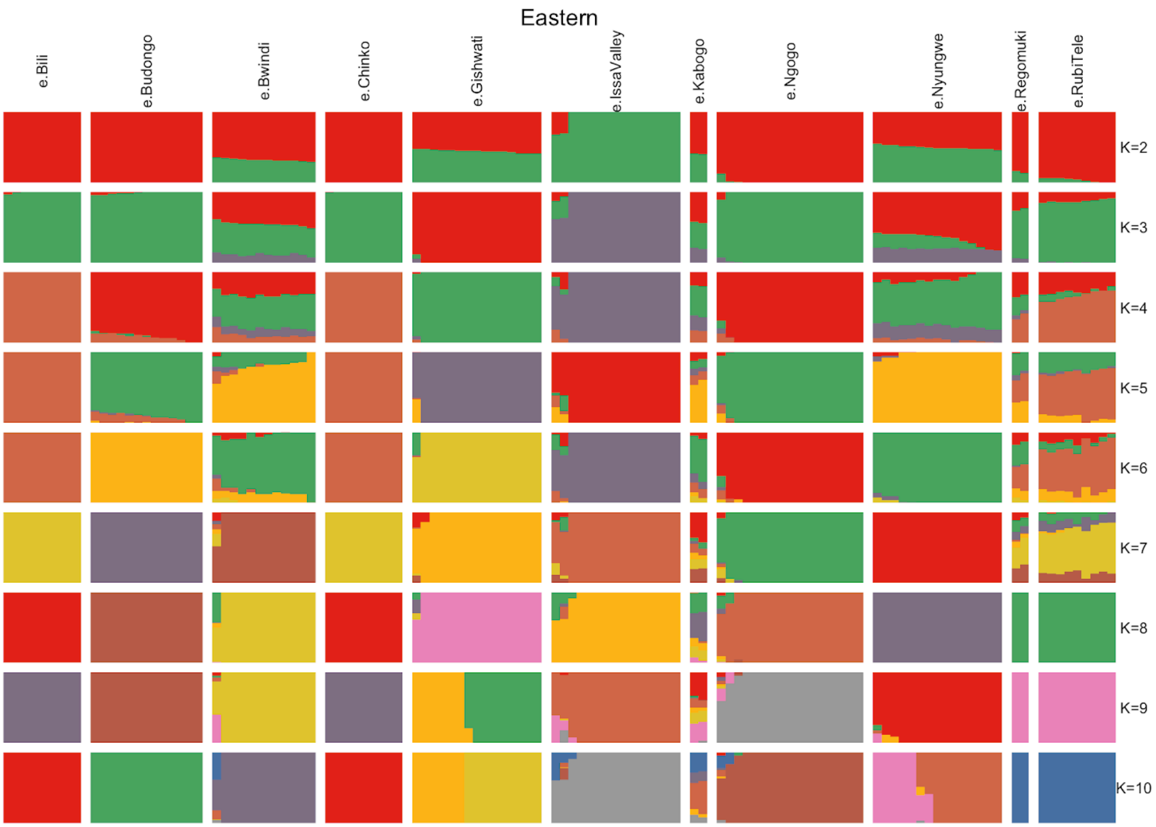

Eastern
e.Bili
e.Budongo
e.Bwindi
e.Chinko
e.Gishwati
e.IssaValley
e.Kabogo
e.Ngogo
e.Nyungwe
e.Regomuki
e.RubiTele
K=2
K=3
K=4
K=5
K=6
K=7
K=8
K=9
K=10

1

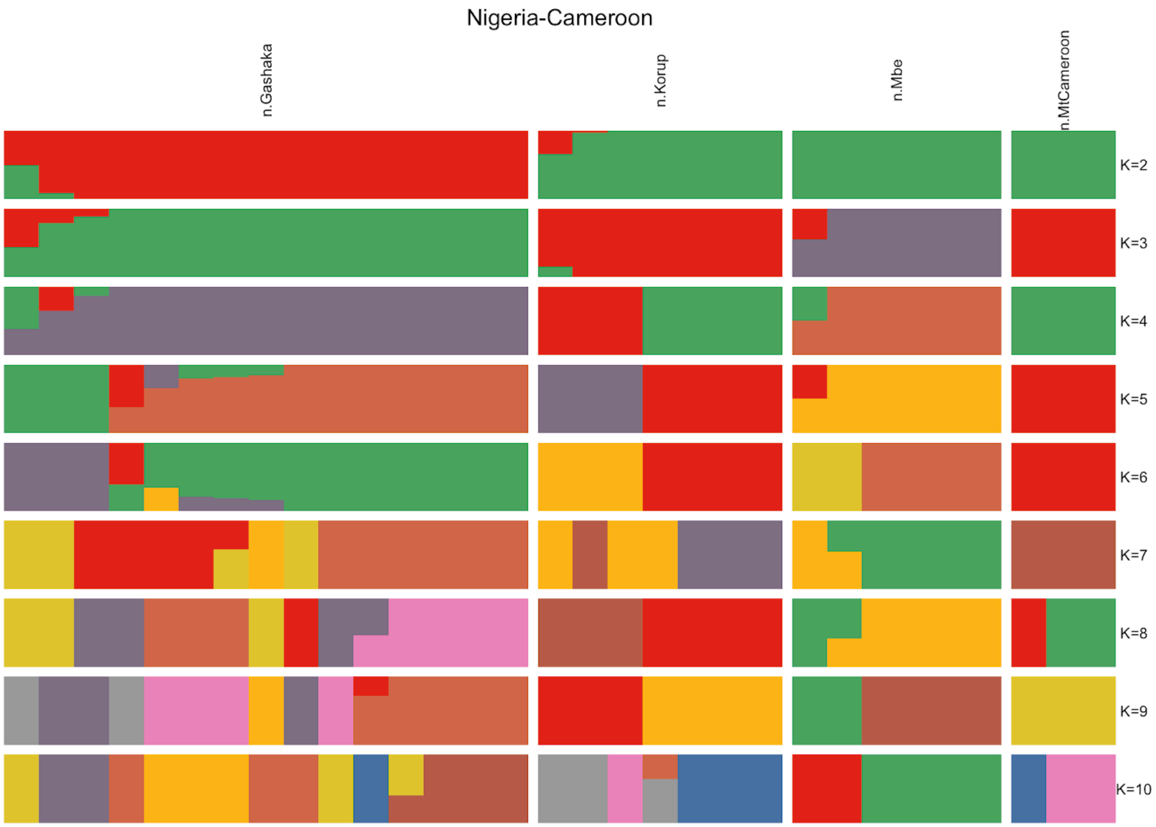
Nigeria-Cameroon
n.Gashaka
n.Korup
n.Mbe
n.MtCameroon
K=2
K=3
K=4
K=5
K=6
K=7
K=8
K=9
K=10

1

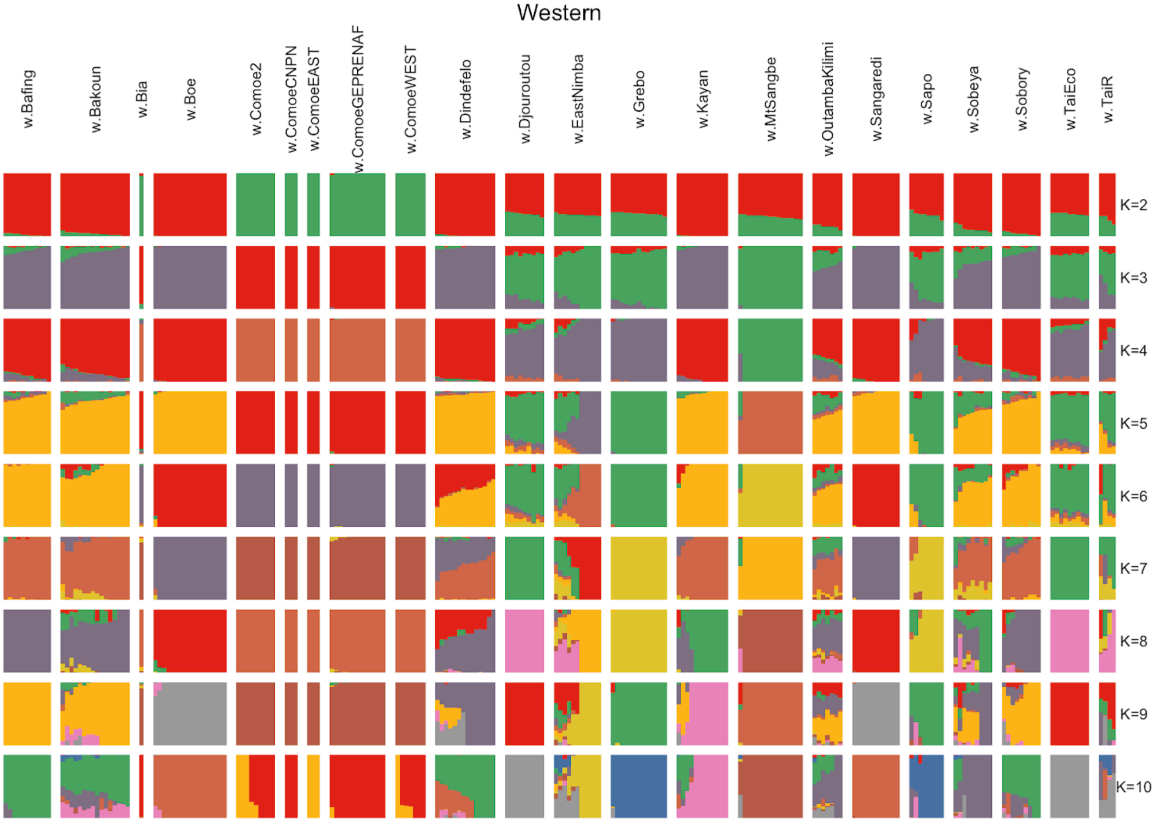


**Fig. S14. NGSadmix results showing ancestry compositions from K=2 to K=10 for all samples combined and each subspecies separately (exome).** Individuals (i.e. columns) are grouped according to subspecies when all samples are analysed together and by sample site where each subspecies is analysed separately. Individuals are ordered within groups according to the proportion of ancestry components.

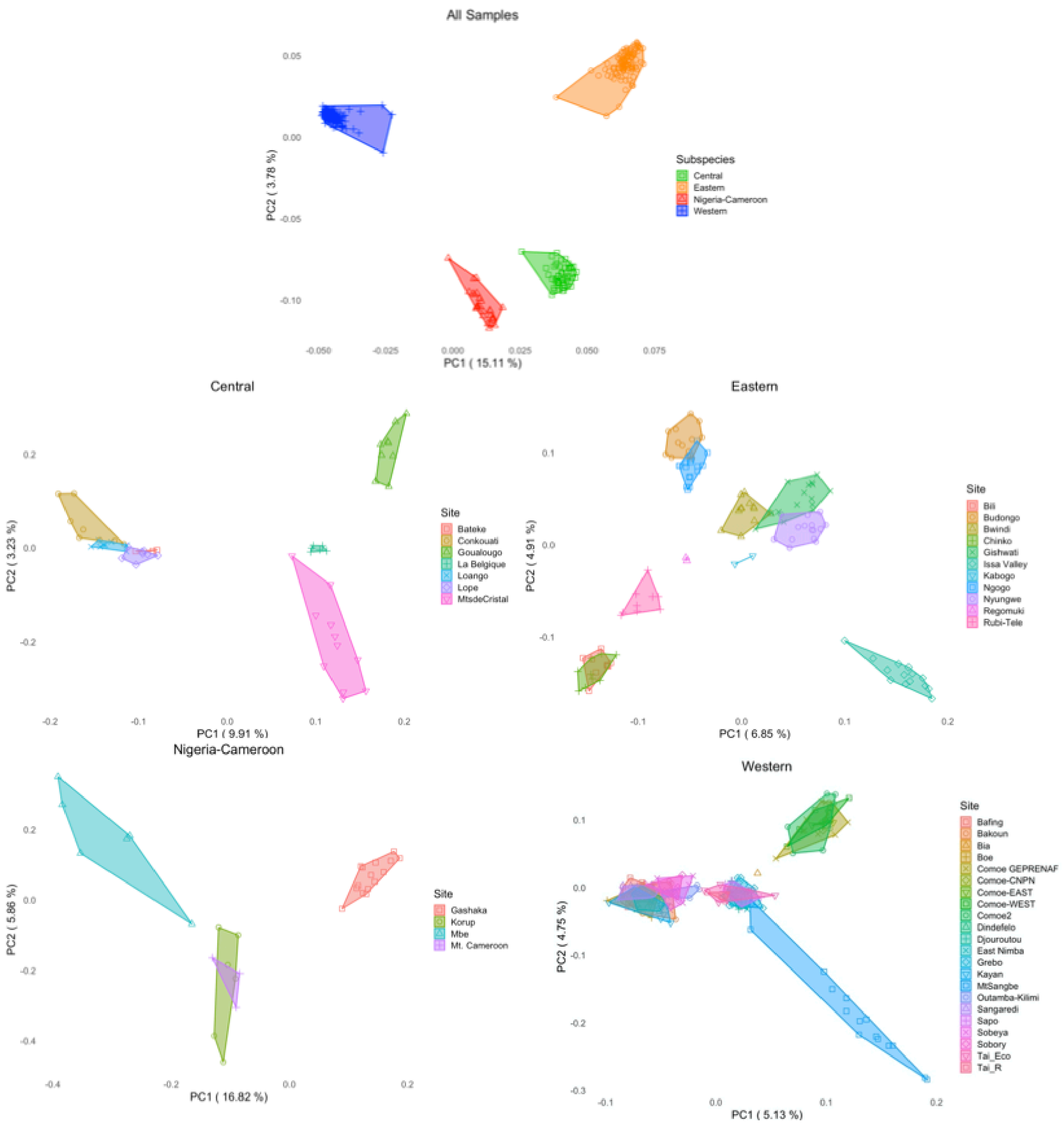


**Fig. S15. PCAs from running PCAngsd on the filtered chr21 for all samples together and each subspecies separately.** Points represent individuals, polygons represent subspecies in the all samples plot and sample sites in the subspecies-specific plots.

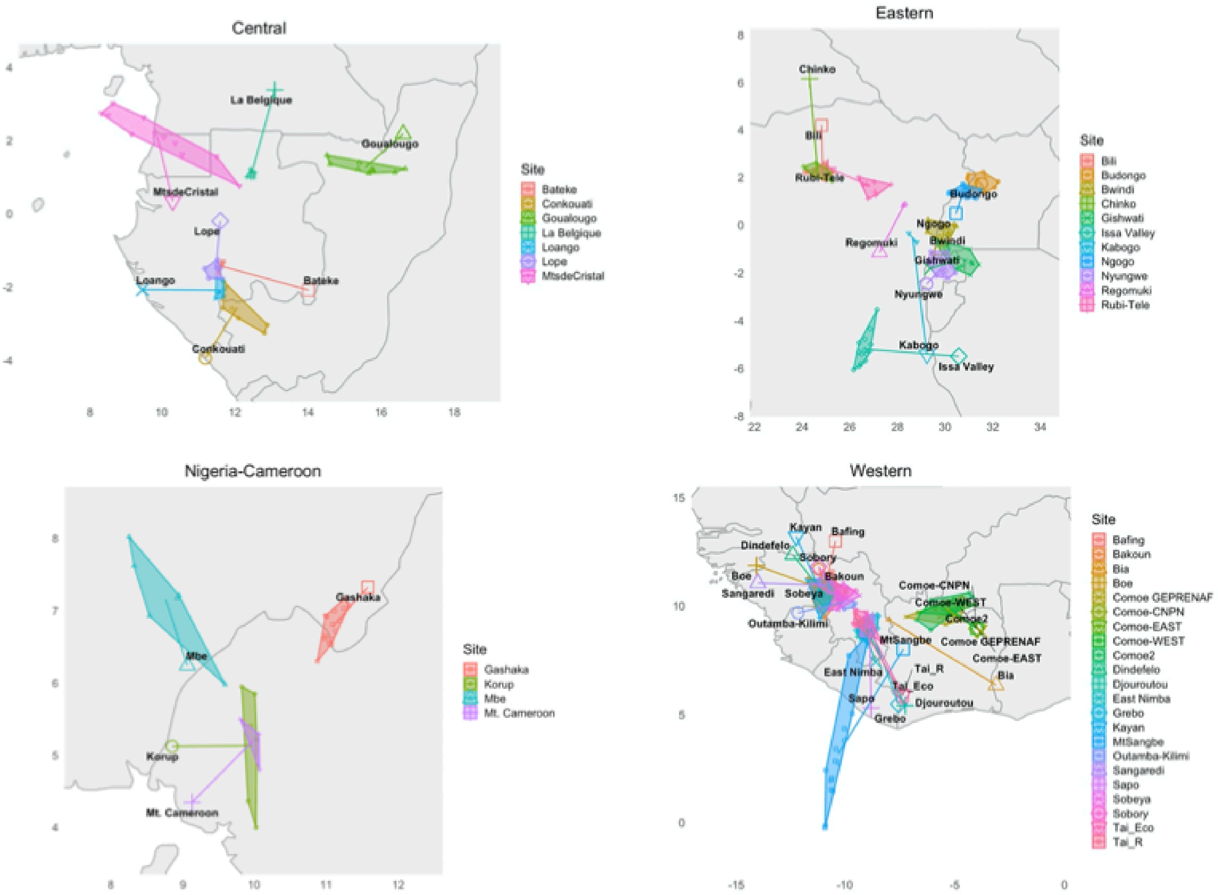


**Fig. S16. PCAs from running PCAngsd on the filtered chr21 for each subspecies separately Procrustes transformed onto a map.** Points represent individuals, polygons represent sample sites and lines link the centre of polygons to the geographical location of that sample site on the map.

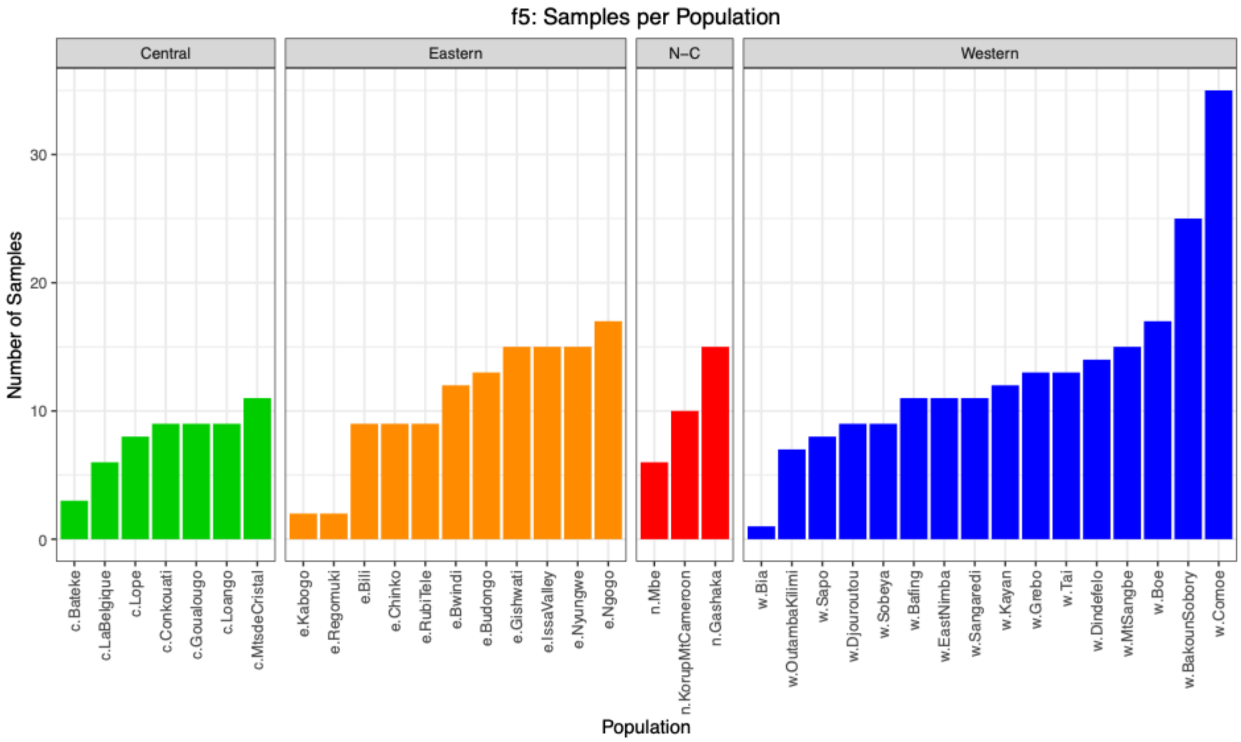


**Fig. S17. The number of samples per population after sample filtering and combining sample sites (exome).**

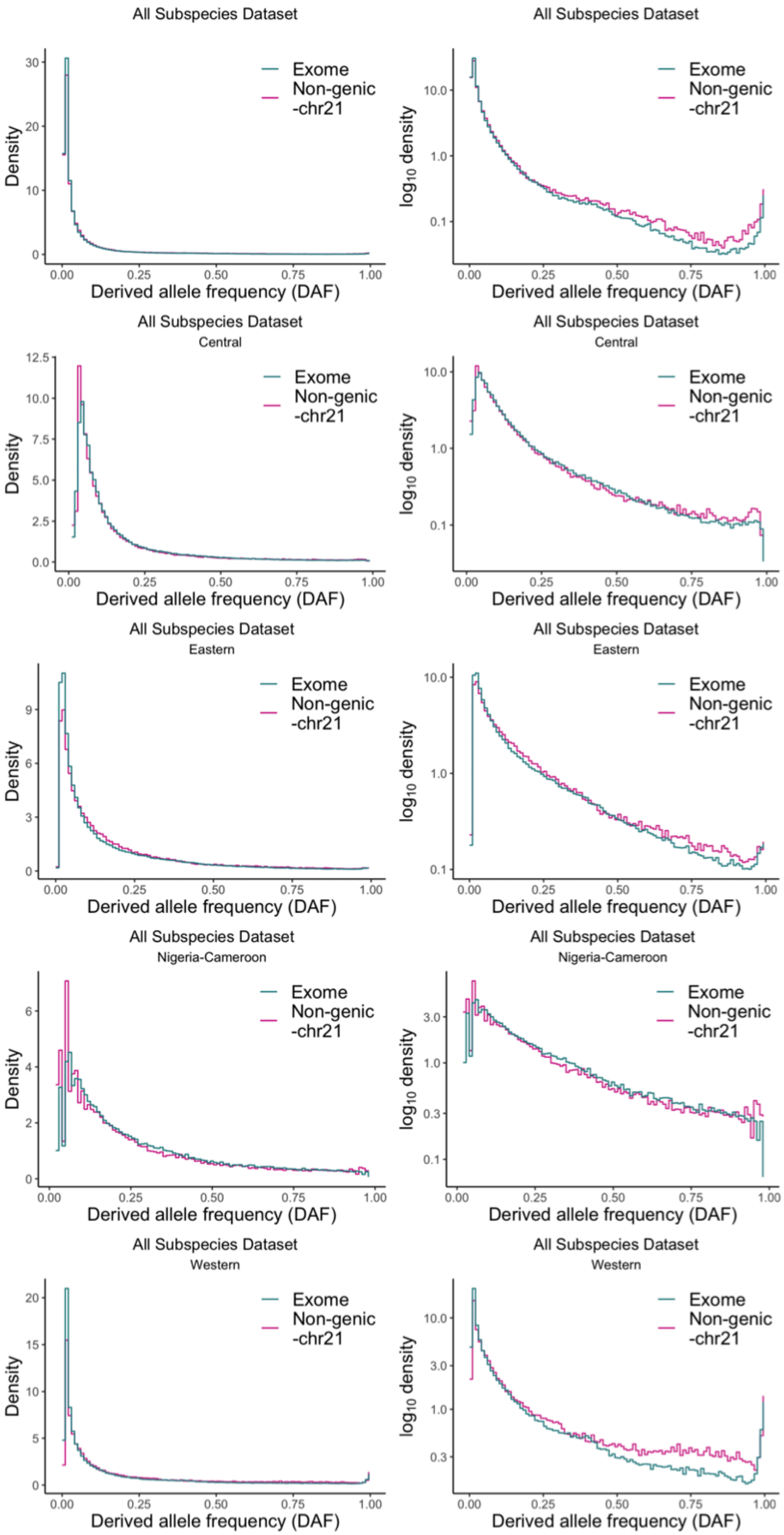
All Subspecies Dataset
Exome
Non-genic
-chr21
Density
Derived allele frequency (DAF)
All Subspecies Dataset
Exome
Non-genic
-chr21
log10 density
Derived allele frequency (DAF)
All Subspecies Dataset
Central
Exome
Non-genic
-chr21
Density
Derived allele frequency (DAF)
All Subspecies Dataset
Central
Exome
Non-genic
-chr21
log10 density
Derived allele frequency (DAF)
All Subspecies Dataset
Eastern
Exome
Non-genic
-chr21
Density
Derived allele frequency (DAF)
All Subspecies Dataset
Eastern
Exome
Non-genic
-chr21
log10 density
Derived allele frequency (DAF)
All Subspecies Dataset
Nigeria-Cameroon
Exome
Non-genic
-chr21
Density
Derived allele frequency (DAF)
All Subspecies Dataset
Nigeria-Cameroon
Exome
Non-genic
-chr21
log10 density
Derived allele frequency (DAF)
All Subspecies Dataset
Western
Exome
Non-genic
-chr21
Density
Derived allele frequency (DAF)
All Subspecies Dataset
Western
Exome
Non-genic
-chr21
log10 density
Derived allele frequency (DAF)

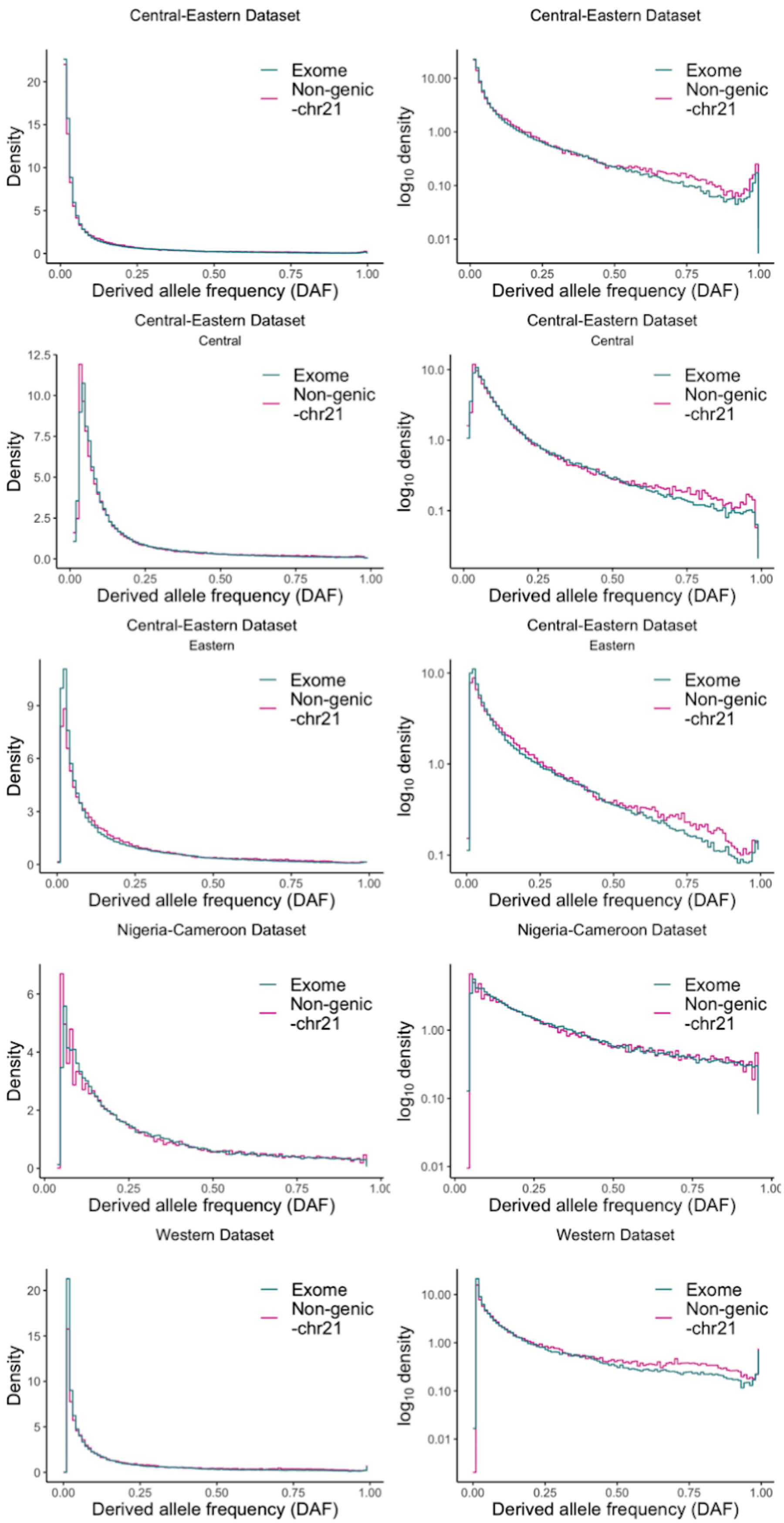
Central-Eastern Dataset
Exome
Non-genic
-chr21
Density
Derived allele frequency (DAF)
Central-Eastern Dataset
log10 density
Central-Eastern Dataset
Central
Central-Eastern Dataset
Eastern
Nigeria-Cameroon Dataset
Western Dataset

**Fig. S18. SFS for each subspecies-dataset.** Left shows the density distribution and the right shows the $\log_{10}$ transformed density. Datasets which contain multiple subspecies (i.e. *All* and *Central-Eastern*) are plotted once using all populations in the dataset and then for each subspecies separately as indicated by the plot subtitle. Fixed sites are not included.

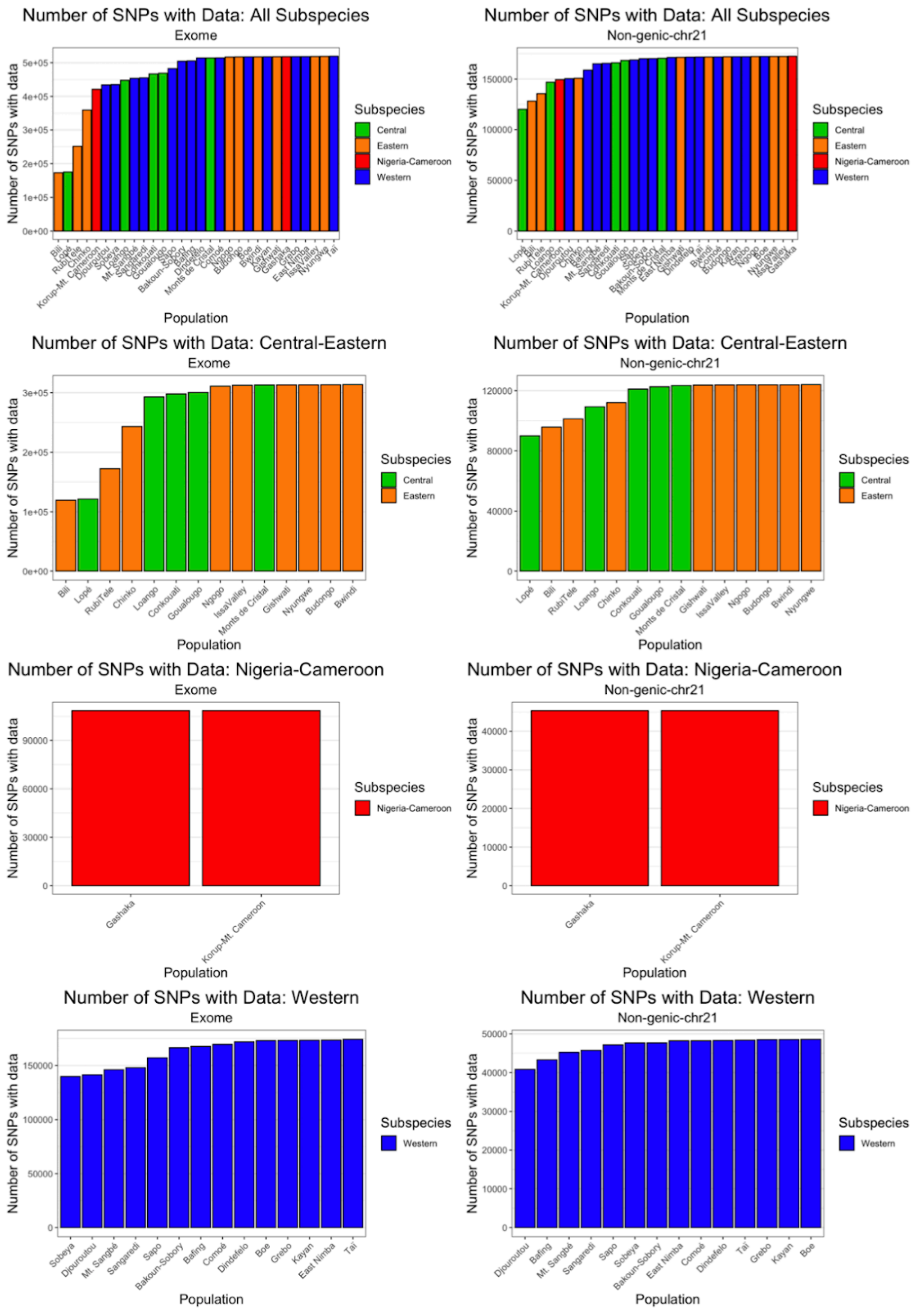


**Fig. S19. The number of sites with allele count data in each population.** Figures plotted for each subspecies-dataset (rows) and either exome (left) or non-genic-chr21 (right) data. Population bars are coloured according to subspecies (central: green, eastern: orange, Nigeria-Cameroon: red, western: blue).

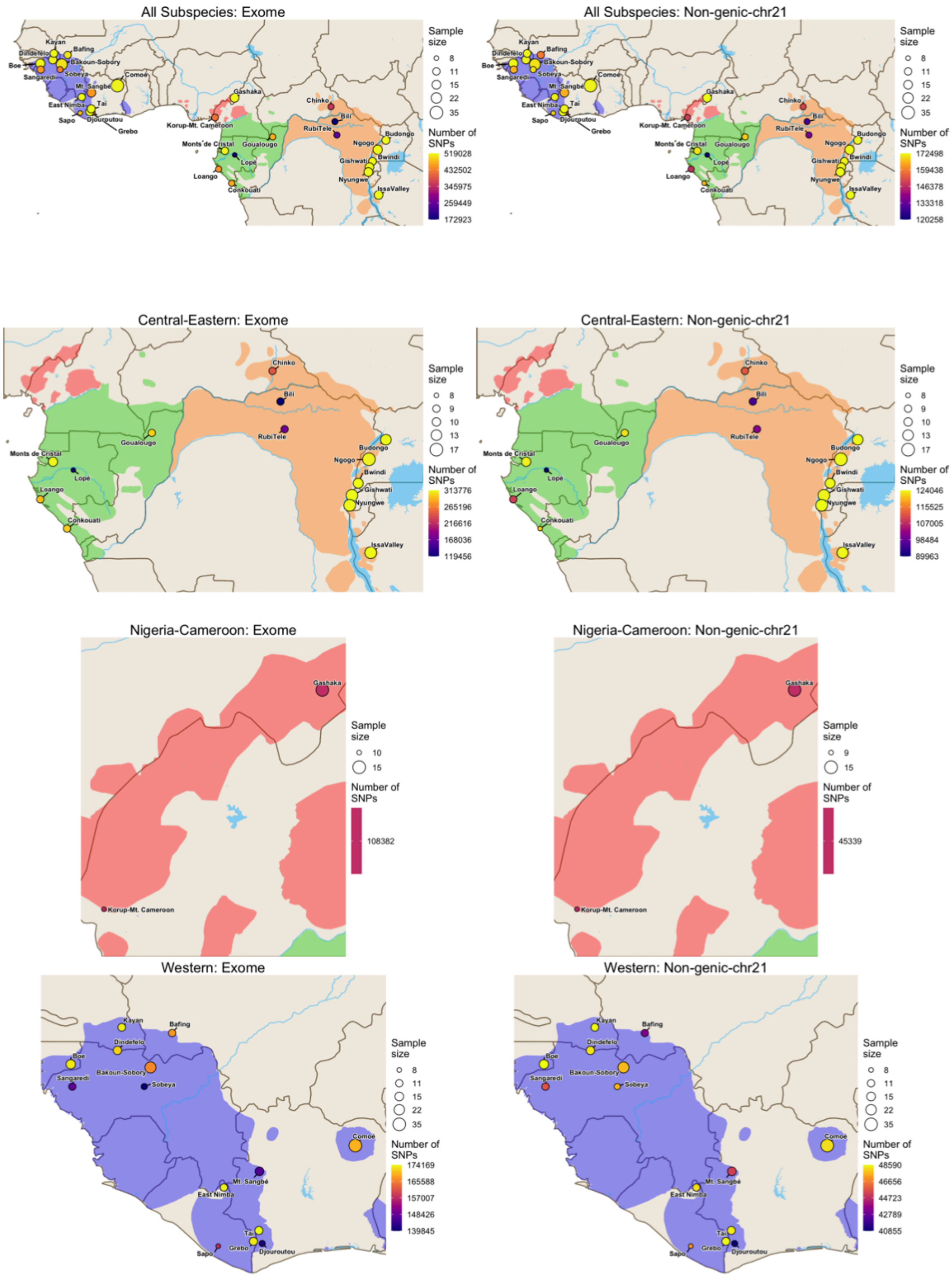


**Fig. S20. Total sample sizes and number of SNPs with allele count data in each population plotted on a map.** Figures plotted for each subspecies-dataset (rows) and either exome (left) or non-genic-chr21 (right) data. Distributions of the four subspecies are indicated with unique colours (green=central, orange=eastern, red=Nigeria-Cameroon, blue=western) and major rivers and lakes are indicated in light blue.

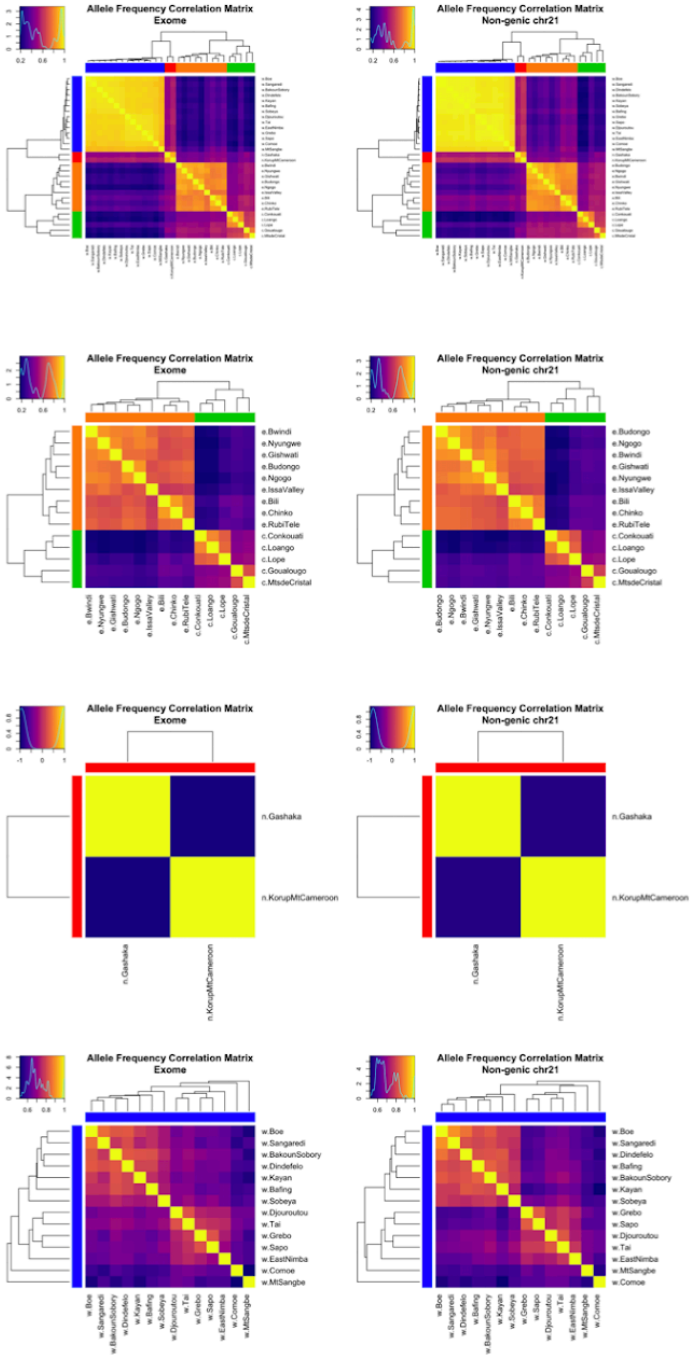


**Fig. S21. Population allele frequency covariance matrices estimated under the BayPass core model visualised as correlation matrices.** Figures plotted for each subspecies-dataset (rows) and either exome (left) or non-genic-chr21 (right). Lighter colours indicate larger correlation coefficients and the density plot in the top left of each plot shows the distribution of these values. Rows and columns are ordered according to a hierarchical clustering tree generated from the correlation matrix using the average agglomeration method. The tips of the tree are coloured according to subspecies (green=central, orange=eastern, red=Nigeria-Cameroon and blue=western).

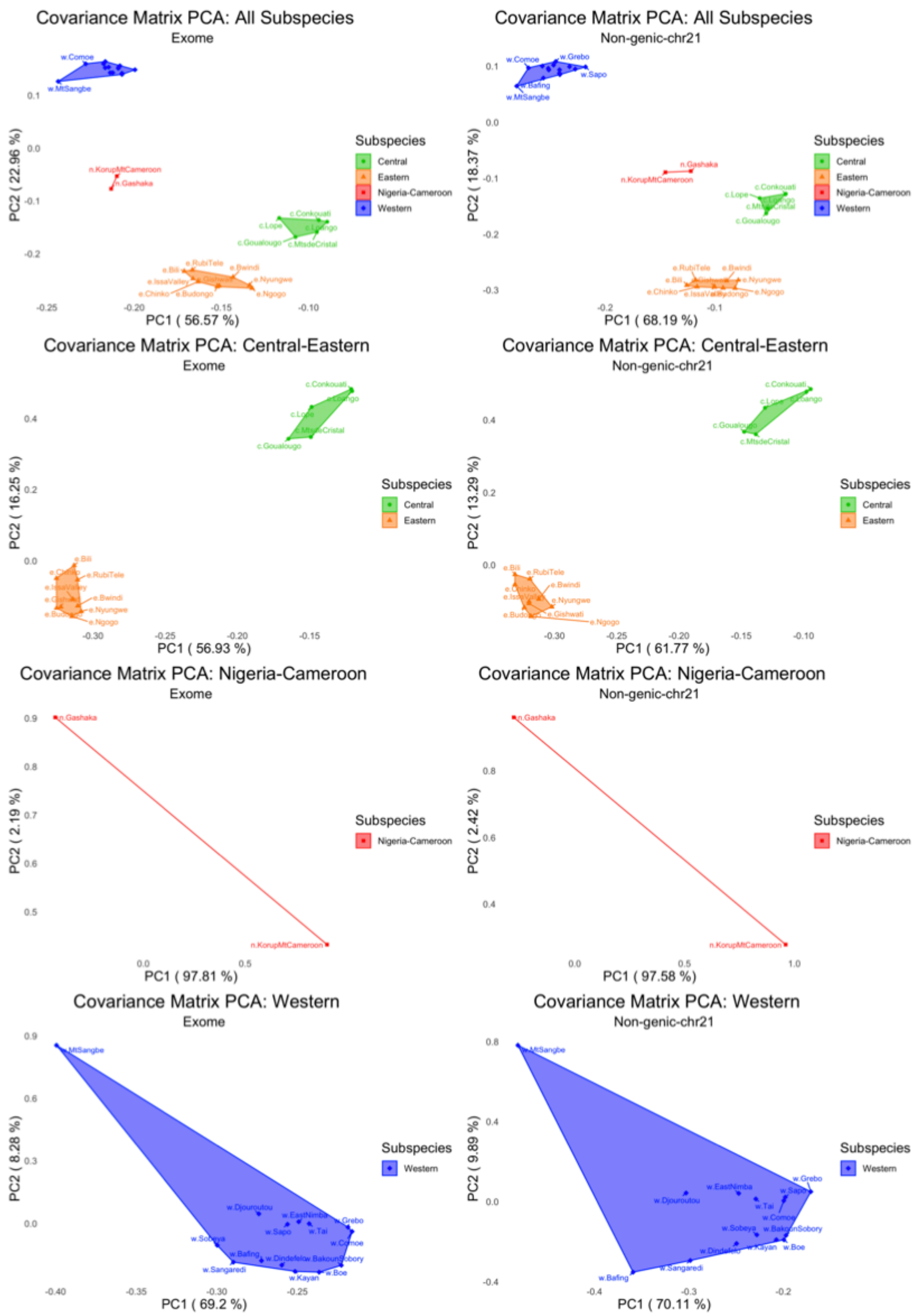


**Fig. S22. PCA plots showing the first two principal components generated from the population allele frequency covariance matrices estimated under the BayPass core model.** Figures plotted for each subspecies-dataset (rows) and either exome (left) or non-genic-chr21 (right) data. Points represent populations which are linked in a polygon coloured according to subspecies (green=central, orange=eastern, red=Nigeria-Cameroon and blue=western).

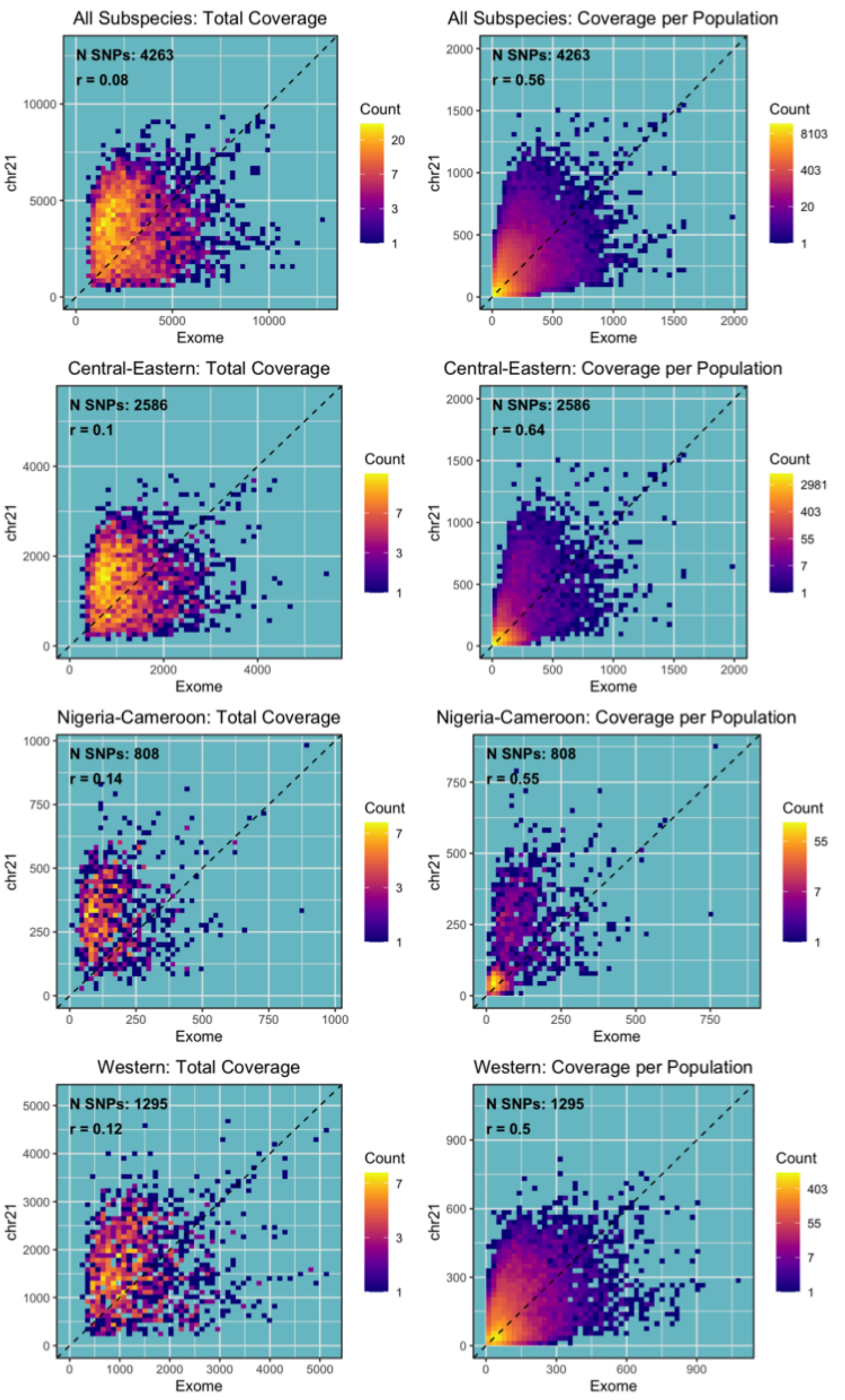


**Fig. S23. 2D density plots showing the coverage for exonic chr21 SNPs in the filtered exome and chr21 capture data.** The left side plots the density distribution of the total coverage across all samples in the dataset while the right plots the density distribution of the total coverage per population for the same SNPs. x=y is shown as a dashed line. The number of SNPs and Pearson correlation coefficient (r) is reported in the top left. We see a slight bias towards higher coverage values in the chr21 capture data compared to the exome capture data for the same SNPs.

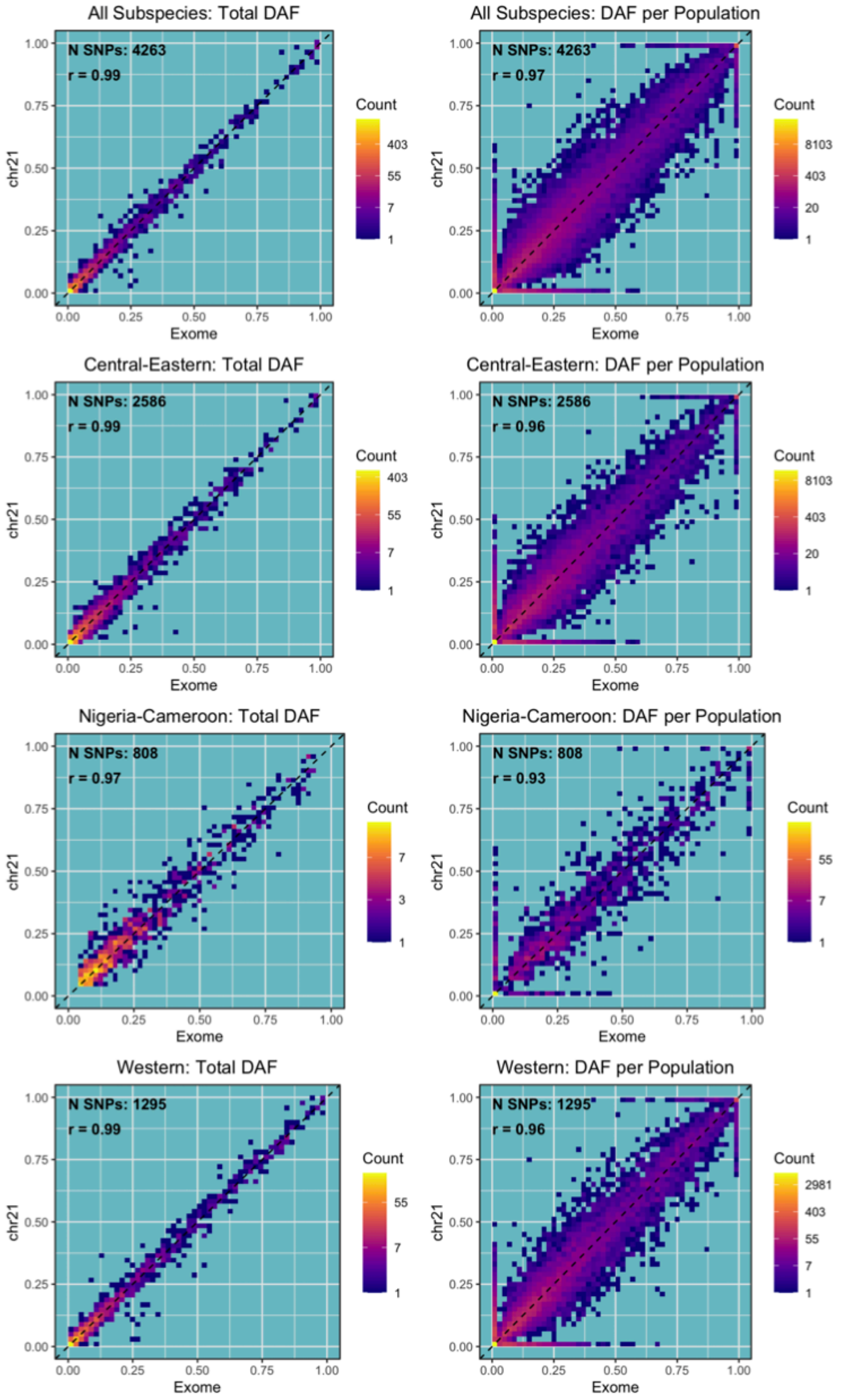


**Fig. S24. 2D density plots showing the allele frequencies estimated for exonic chr21 SNPs in the filtered exome and chr21 capture data.** The left side plots the density distribution of the allele frequencies across all populations in the dataset while the right plots the density distribution of the per-population allele frequencies for the same SNPs. x=y is shown as a dashed line. The number of SNPs and Pearson correlation coefficient (r) is reported in the top left. We see a strong correlation between the two and no clear bias caused by the different capture methods.

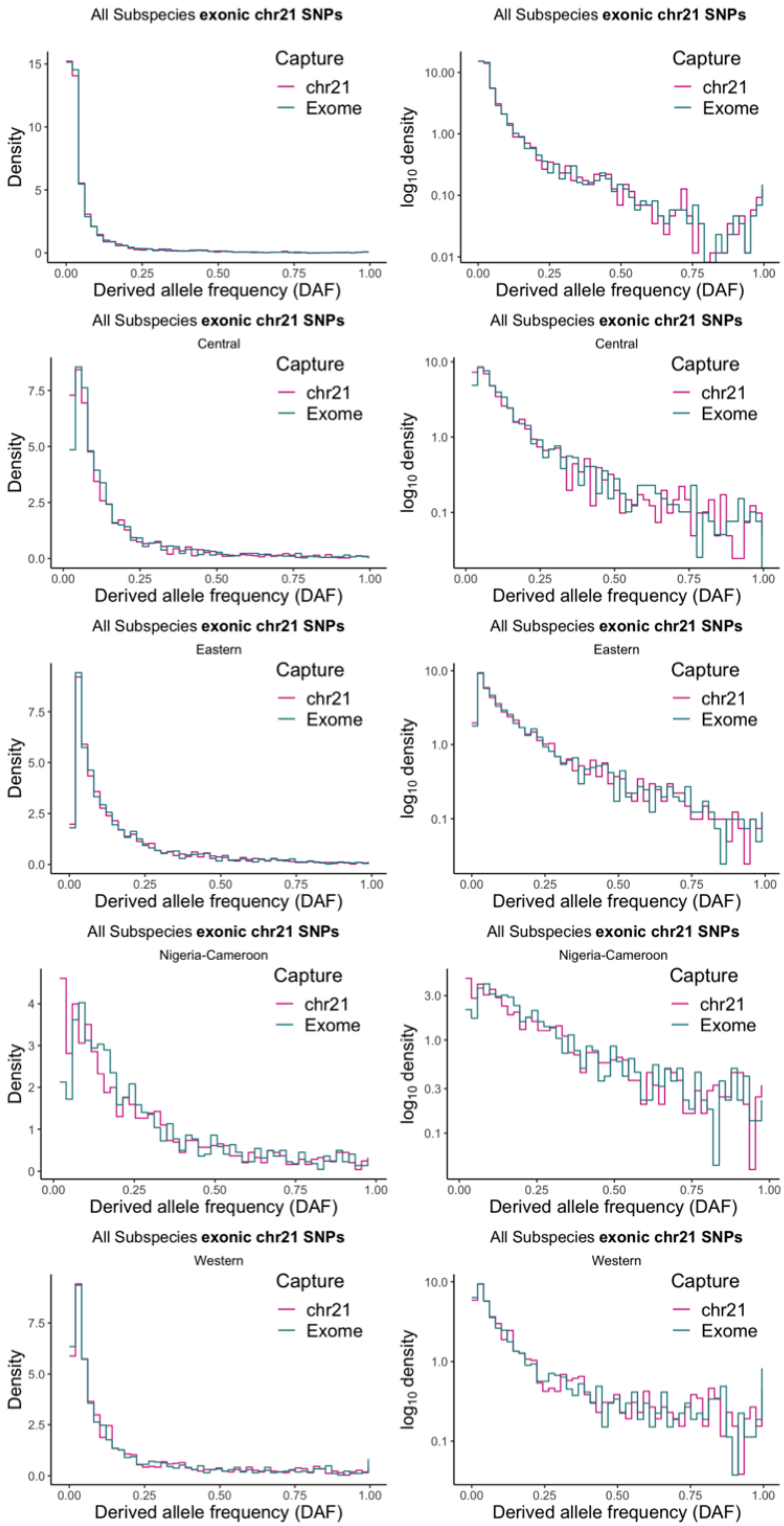
All Subspecies exonic chr21 SNPs
Capture
chr21
Exome
Density
Derived allele frequency (DAF)
All Subspecies exonic chr21 SNPs
Capture
chr21
Exome
log10 density
Derived allele frequency (DAF)
All Subspecies exonic chr21 SNPs
Central
Capture
chr21
Exome
Density
Derived allele frequency (DAF)
All Subspecies exonic chr21 SNPs
Central
Capture
chr21
Exome
log10 density
Derived allele frequency (DAF)
All Subspecies exonic chr21 SNPs
Eastern
Capture
chr21
Exome
Density
Derived allele frequency (DAF)
All Subspecies exonic chr21 SNPs
Eastern
Capture
chr21
Exome
log10 density
Derived allele frequency (DAF)
All Subspecies exonic chr21 SNPs
Nigeria-Cameroon
Capture
chr21
Exome
Density
Derived allele frequency (DAF)
All Subspecies exonic chr21 SNPs
Nigeria-Cameroon
Capture
chr21
Exome
log10 density
Derived allele frequency (DAF)
All Subspecies exonic chr21 SNPs
Western
Capture
chr21
Exome
Density
Derived allele frequency (DAF)
All Subspecies exonic chr21 SNPs
Western
Capture
chr21
Exome
log10 density
Derived allele frequency (DAF)

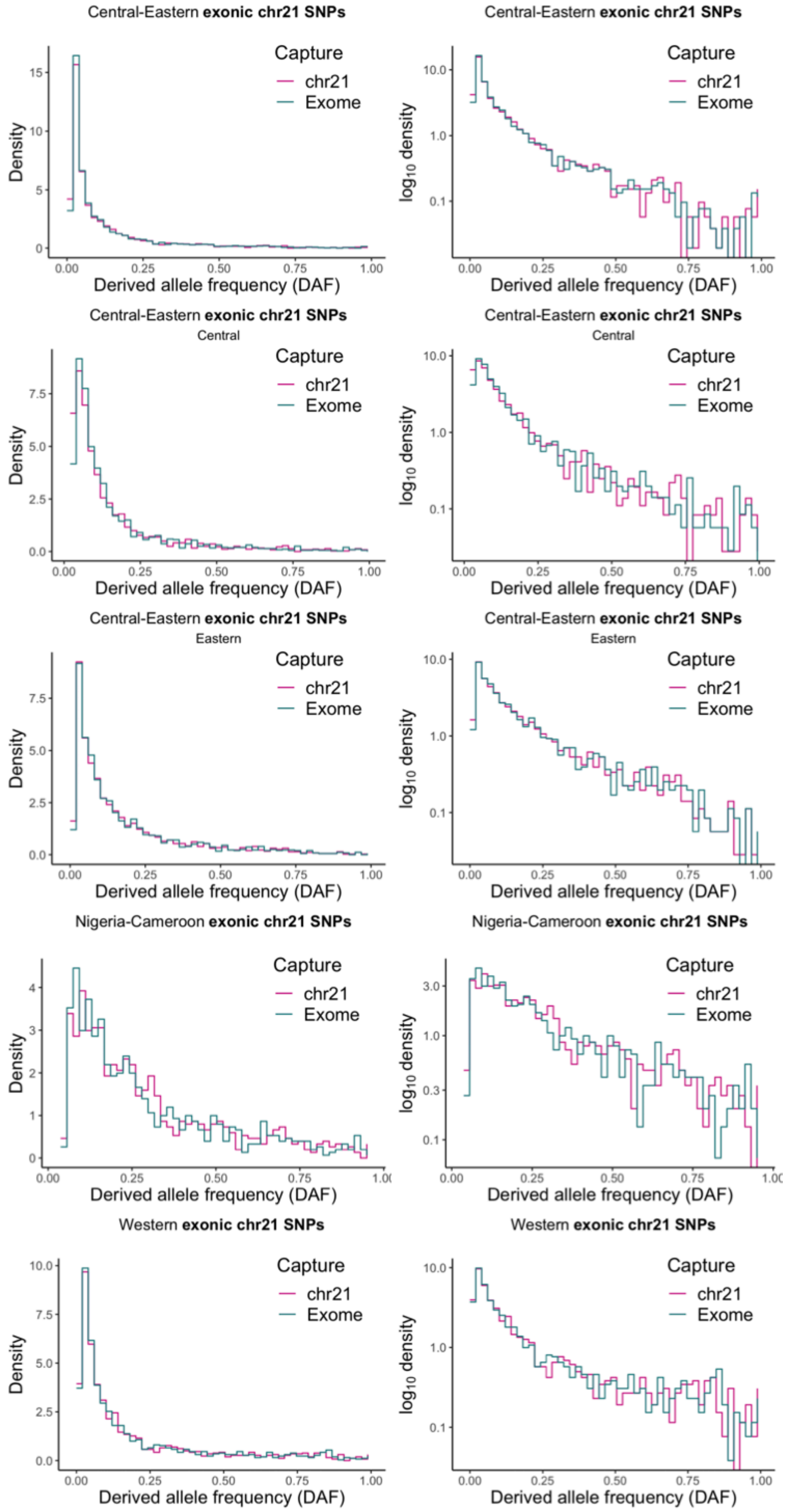
Central-Eastern **exonic chr21 SNPs**
Central
Eastern
Nigeria-Cameroon **exonic chr21 SNPs**
Western **exonic chr21 SNPs**
Capture
chr21
Exome
Density
$\log_{10}$ density
Derived allele frequency (DAF)

**Fig. S25. SFS for each subspecies-dataset for exonic chr21 SNPs present in both the filtered exome capture and chr21 capture data.** SFS for the exome (green) and chr21 capture data (purple) correspond very closely. Left shows the density distribution and the right shows the $\log_{10}$ transformed density. Datasets which contain multiple subspecies (i.e. *All* and *Central-Eastern*) are plotted once using all populations in the dataset and then for each subspecies separately as indicated by the plot subtitle. Fixed sites are not included.

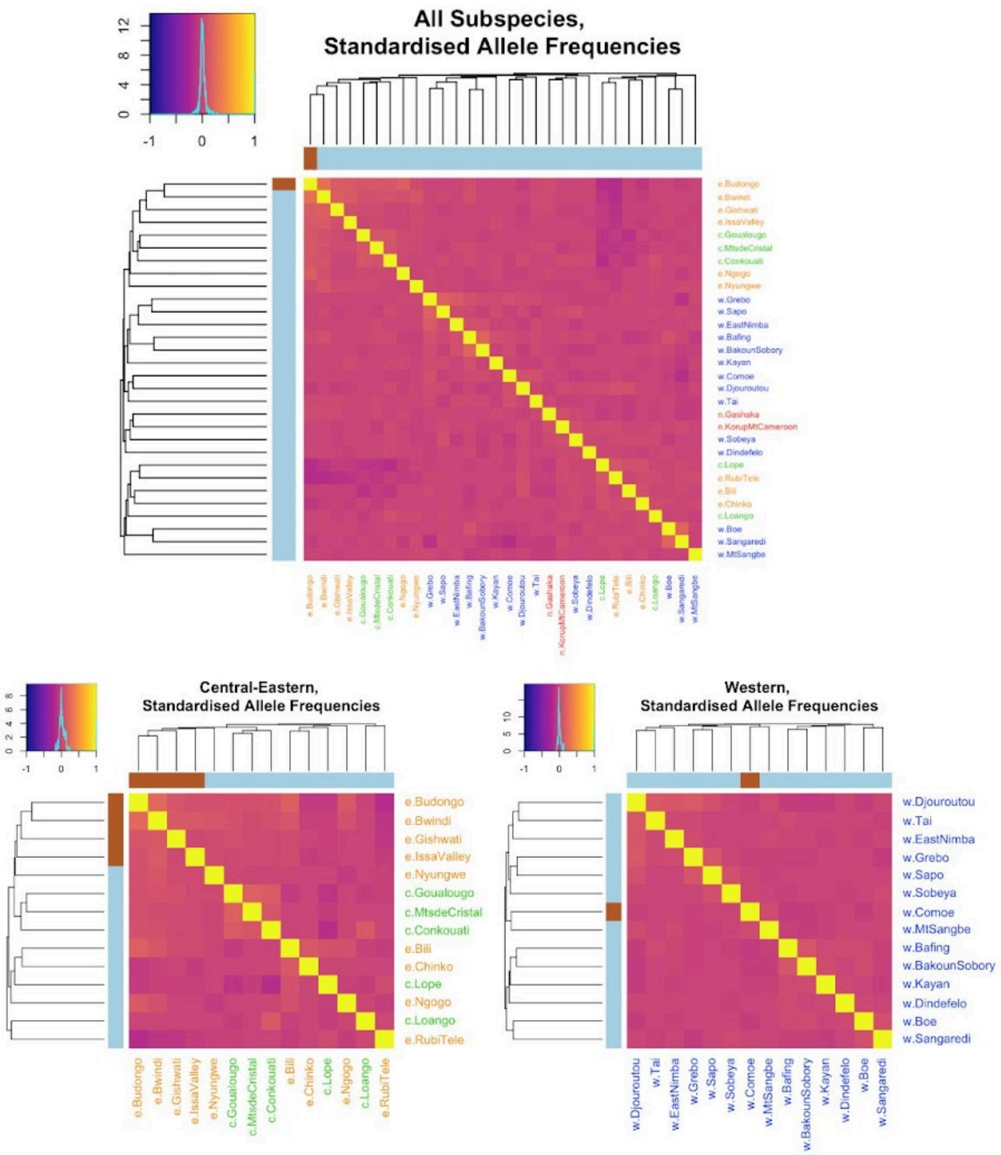


**Fig. S26. Correlation matrices calculated from the standardised allele frequencies calculated under the BayPass core model of 5,000 randomly sampled exonic SNPs.** Results for *Nigeria-Cameroon* are not shown as the correlation matrix formed from only two populations is uninformative. Lighter colours indicate larger correlation coefficients and the density plot in the top left of each plot shows the distribution of these values. Rows and columns are ordered according to a hierarchical clustering tree generated from the correlation matrix using the average agglomeration method. The tips of the tree are coloured according to clusters assigned by k-medoids clustering for the estimated best k. Population names are written in colours corresponding to their subspecies (green=central, orange=eastern, red=Nigeria-Cameroon and blue=western). We can see that unlike the correlation matrices and PCAs estimated with the raw allele frequencies (Fig. S21 and Fig. S22), there is no population structure observed in the standardised allele frequencies (unlike for raw allele frequencies shown in Fig. S21) indicating that BayPass does account for population structure.

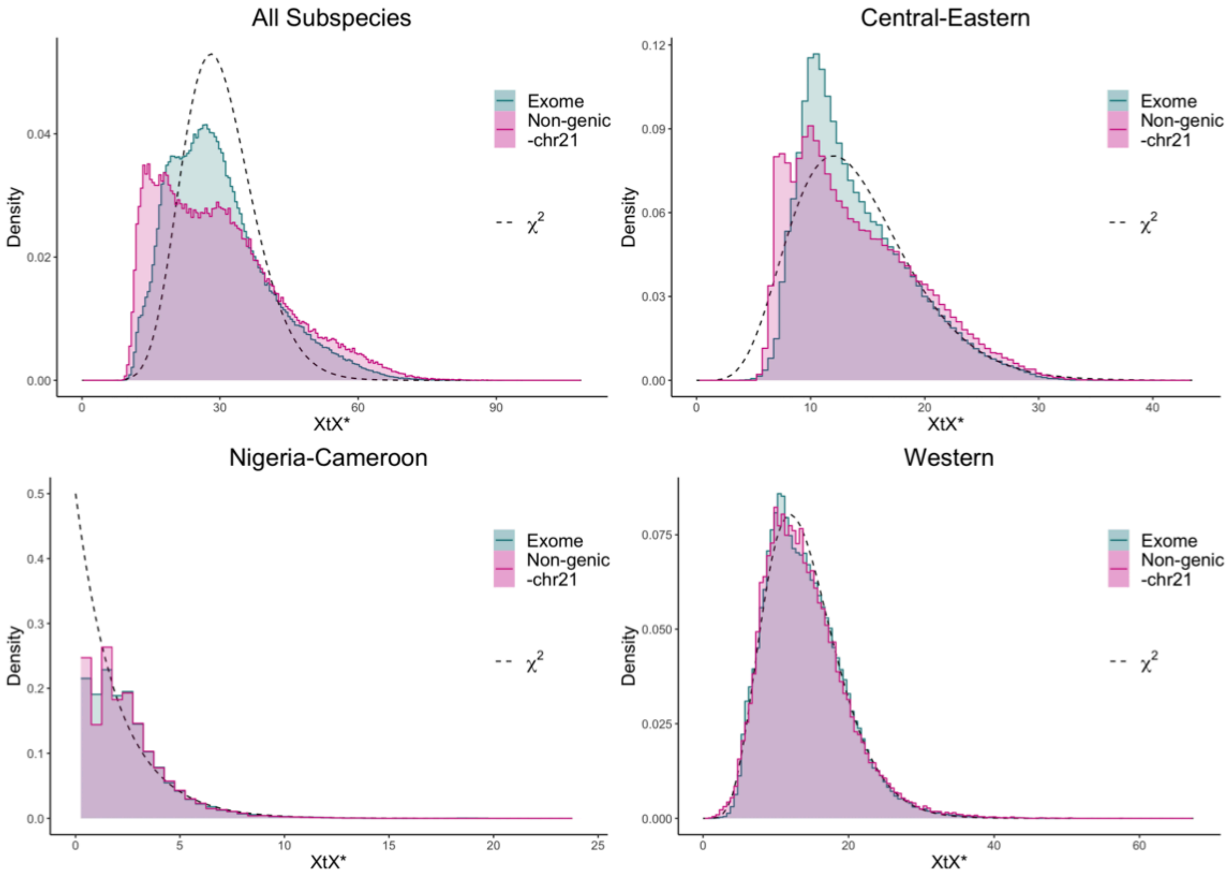


**Fig. S27. X$^t$X* distributions of exome and non-genic-chr21 for each of the subspecies-datasets compared to the $\chi^2_J$ distribution (*J*=N populations).**

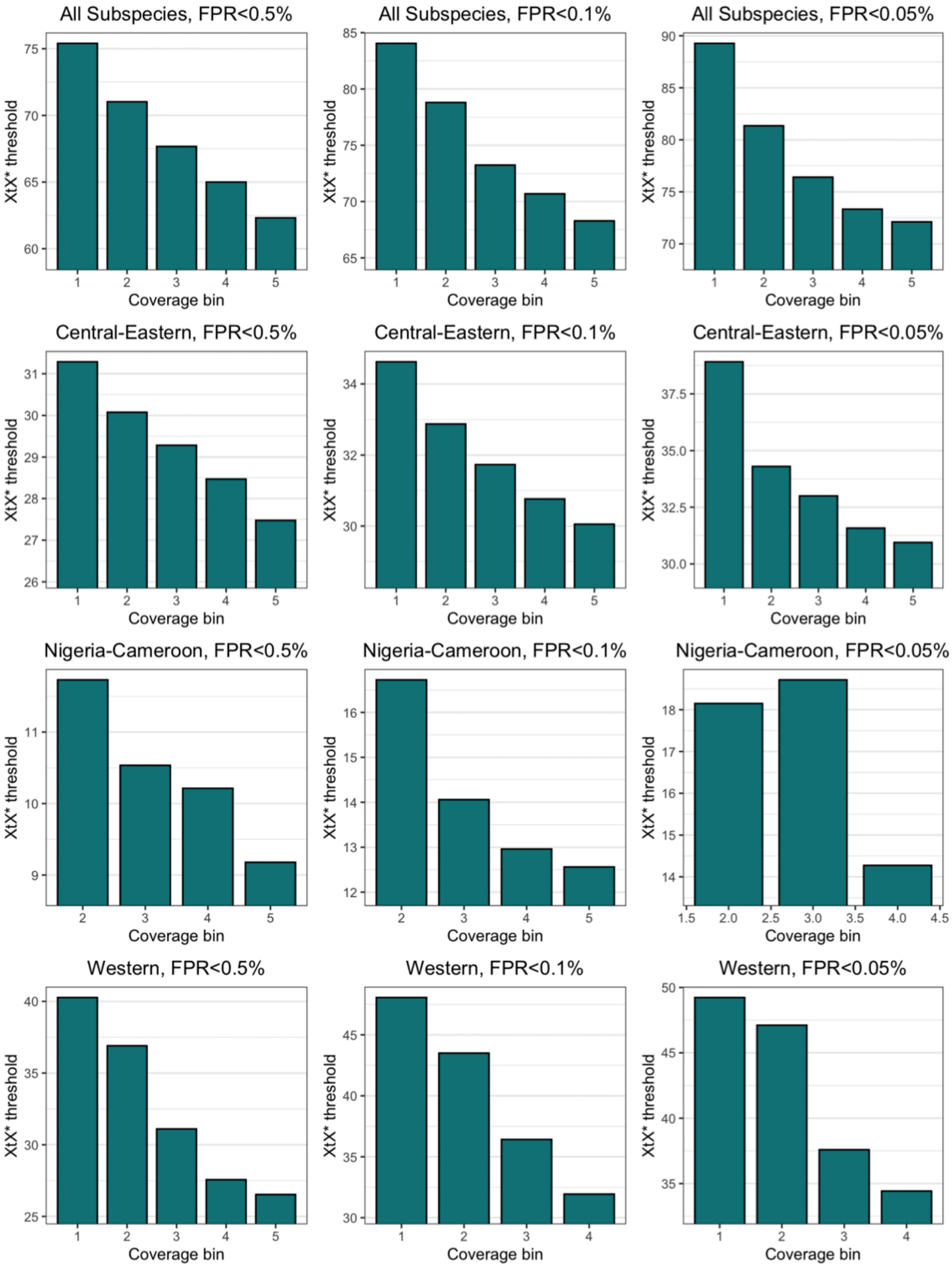


**Fig. S28. $X^tX^*$ thresholds for each coverage bin for each FPR tail (columns) and subspecies-dataset (rows).**

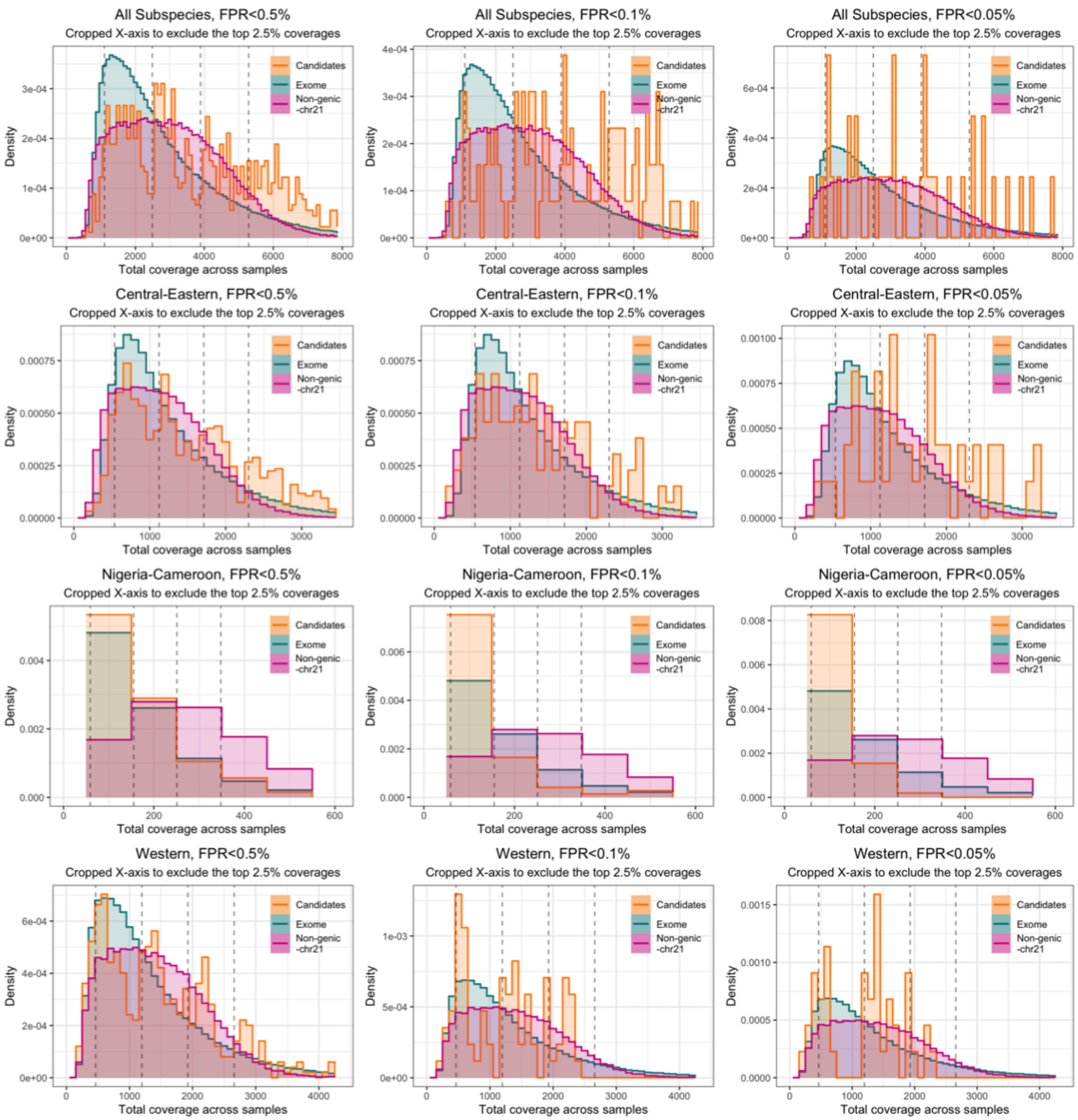


**Fig. S29. Distribution of total coverage across samples for the whole exome (green), non-genic-chr21 (purple) and the genetics-only candidates (orange).** Results shown for each FPR tail and each subspecies-dataset. The limits of the 5 coverage bins are shown as vertical dashed lines.

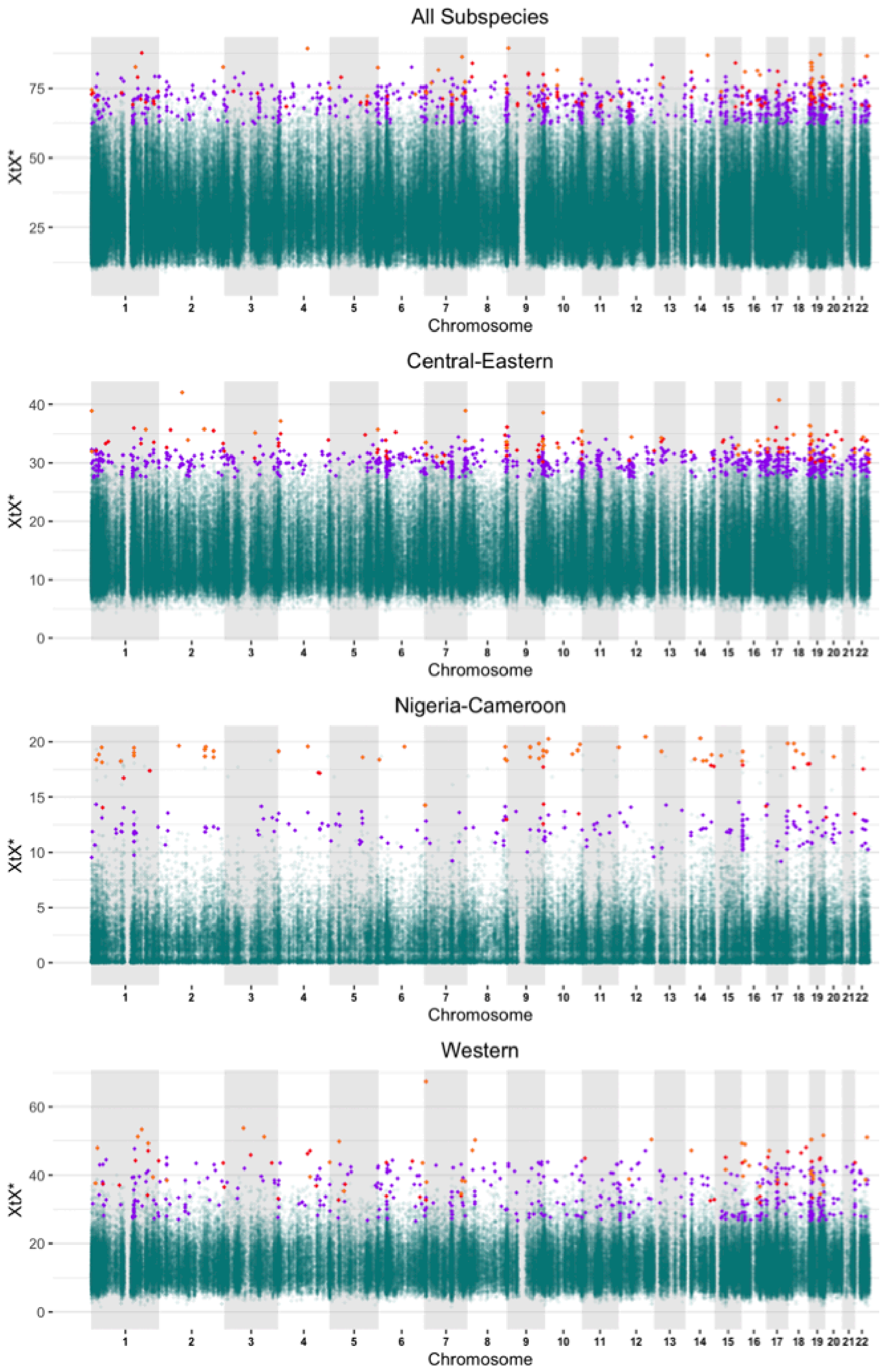


**Fig. S30. Manhattan plots showing the $X^tX^*$ values for each dataset.** Points coloured purple, red and orange are in the 0.5%, 0.1% and 0.05% FPR tails respectively.

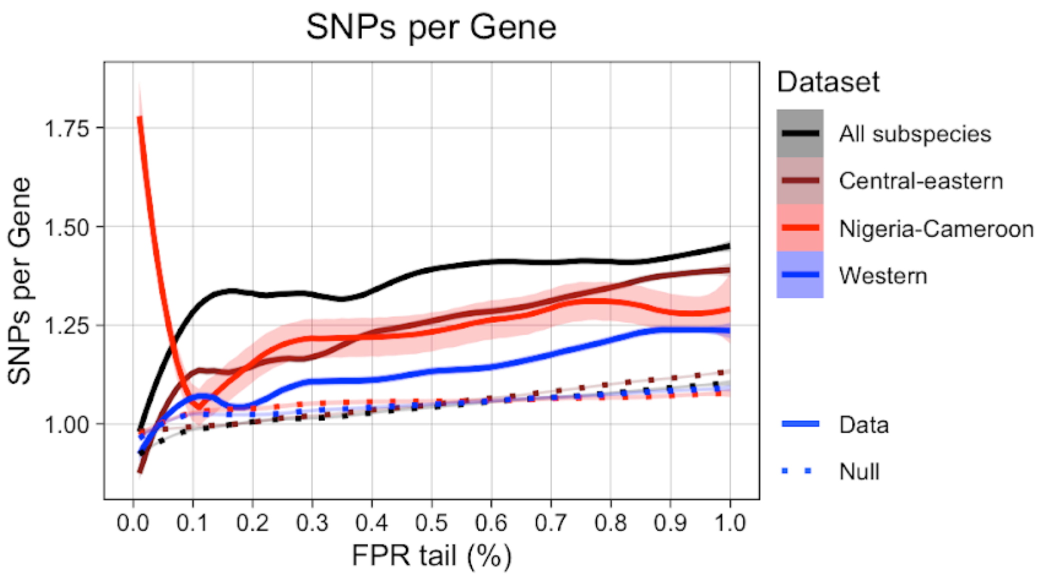


**Fig. S31. Number of SNPs per gene across FPR tails for the four subspecies-datasets analyses in the genetic-only test.** Solid lines represent the real data sampled at 0.01% intervals and dotted lines represent null expectations generated by randomly sampling the same number of SNPs 50 times at each 0.01% interval. Shaded areas represent the 95% confidence intervals. *Nigeria-Cameroon* contains the fewest SNPs and therefore there are very few SNPs at very stringent FPR tails resulting in increased stochasticity which explains why the line rises sharply at stringent thresholds.

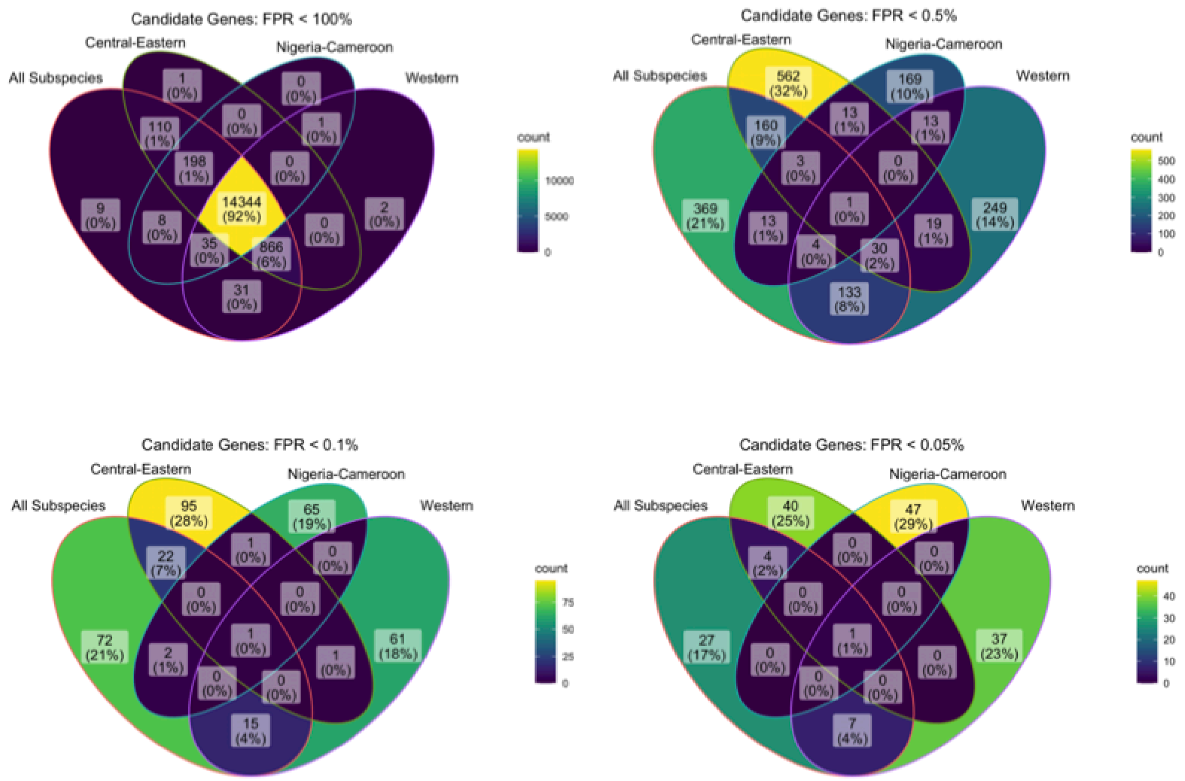


**Fig. S32. Overlap in SNPs between different subspecies-datasets in all SNPs (i.e. FPR<100%) and the 0.5%, 0.1% and 0.05% FPR tails from the genetics-only analysis.**

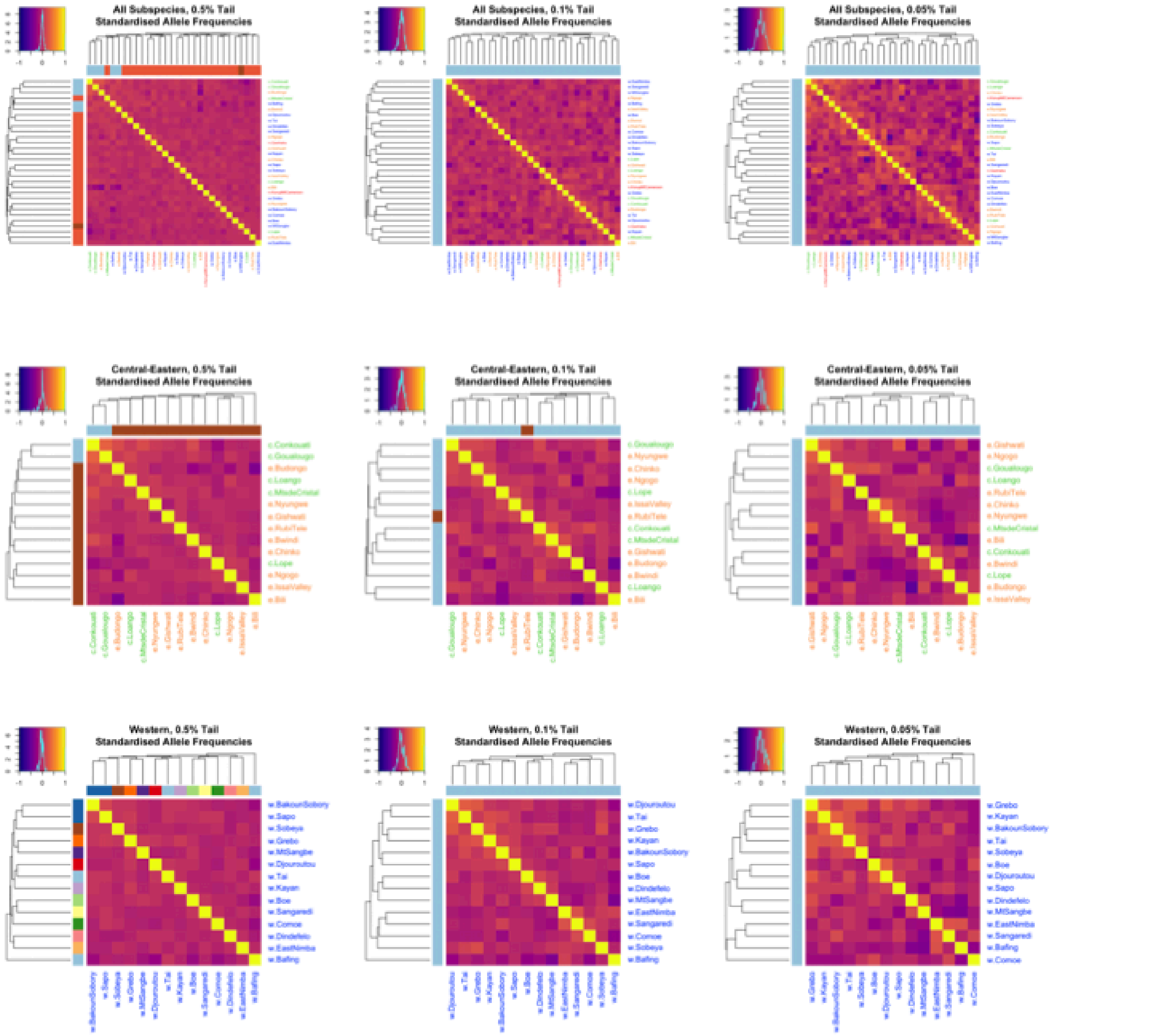


**Fig. S33. Correlation matrices calculated from the exome standardised allele frequencies calculated under the BayPass core model.** Figures are shown for *All*, *Central-Eastern* and *Western* for the 0.5%, 0.1% and 0.05% FPR tails. Results for *Nigeria-Cameroon* are not shown as correlation matrix formed from only two populations is uninformative. Lighter colours indicate larger correlation coefficients and the density plot in the top left of each plot shows the distribution of these values. Populations are ordered according to a dendrogram calculated using the average agglomeration method and the tips are coloured according to clusters assigned by k-medoids clustering for the estimated best k. Population names are coloured according to subspecies (central: green, eastern: orange, Nigeria-Cameroon: red, western: blue). The fact that there is no correspondence with the neutral population structure shown in Figs. S21 and S22 further shows that BayPass is correcting for neutral population structure effectively.

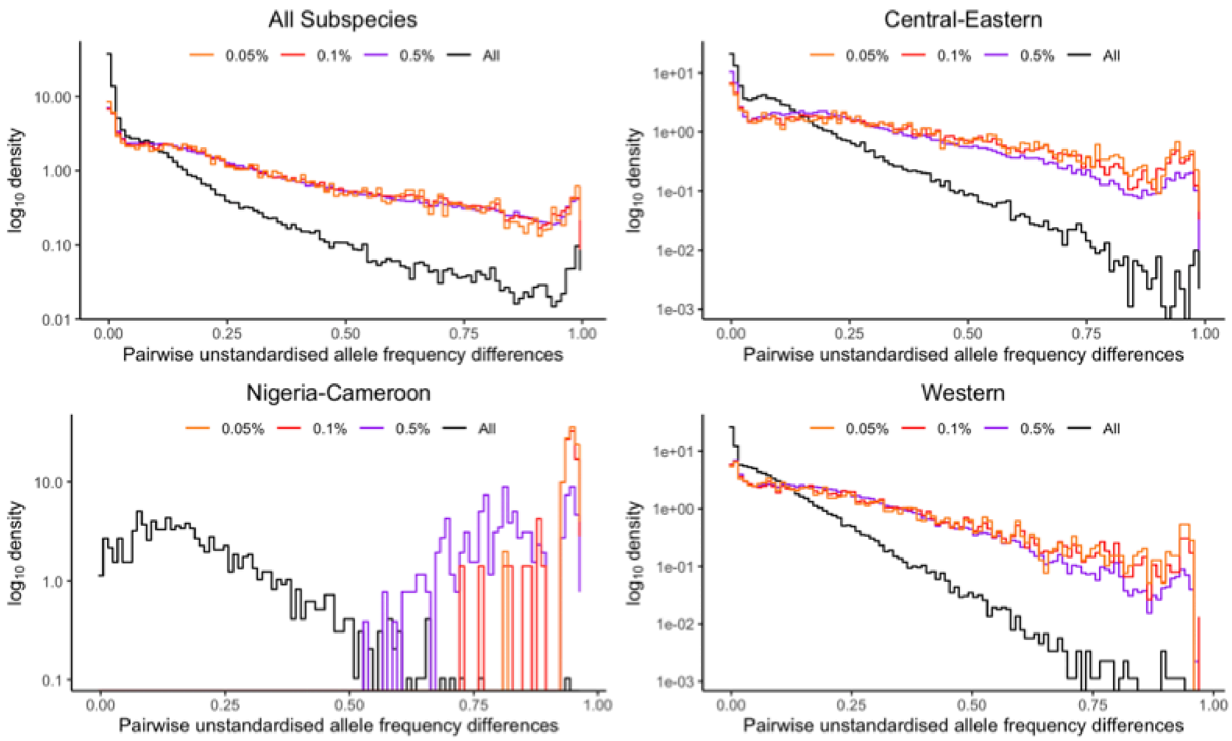


**Fig. S34. Distribution of absolute pairwise population unstandardised allele frequency differences at candidate SNPs.** Results shown for the 0.5%, 0.1% and 0.05% FPR tails compared to a random sample of 1,000 SNPs (All). The y-axis is log-transformed.

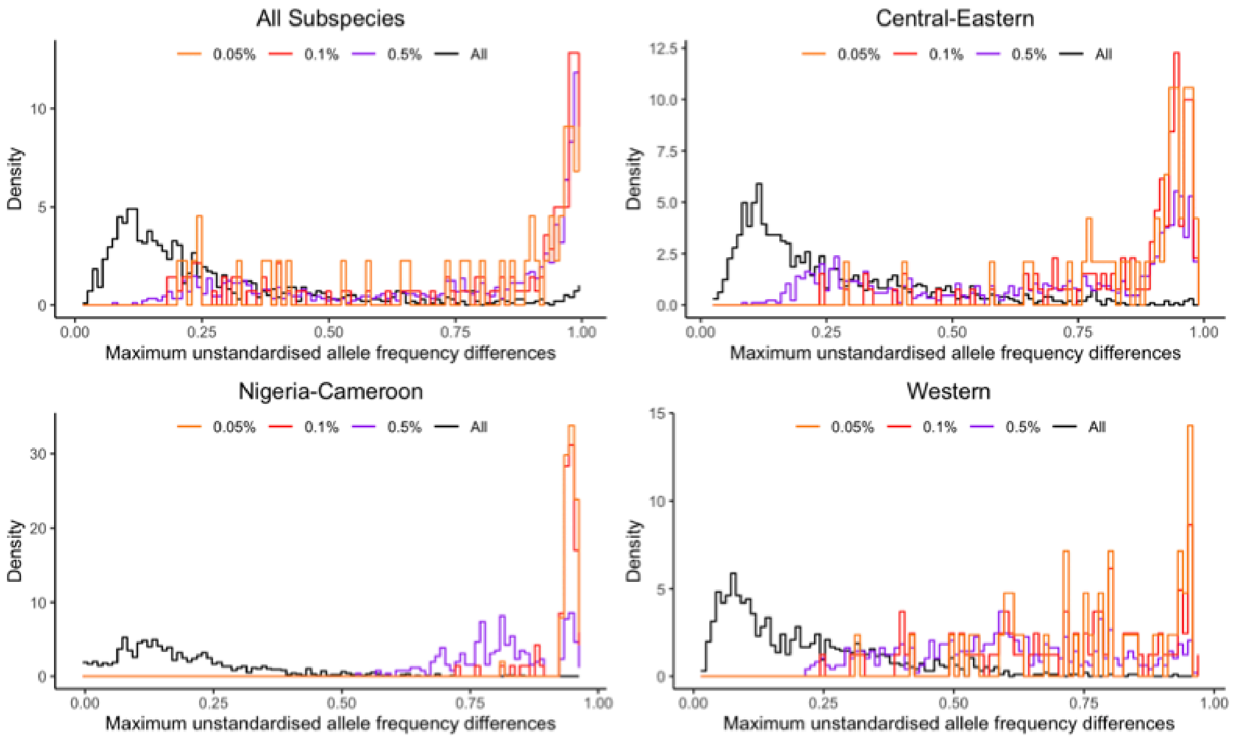


**Fig. S35. Distribution of the maximum absolute pairwise unstandardised population allele frequency differences.** Results shown for candidate SNPs in the 0.5%, 0.1% and 0.05% FPR tails compared to a random sample of 1,000 SNPs (All).

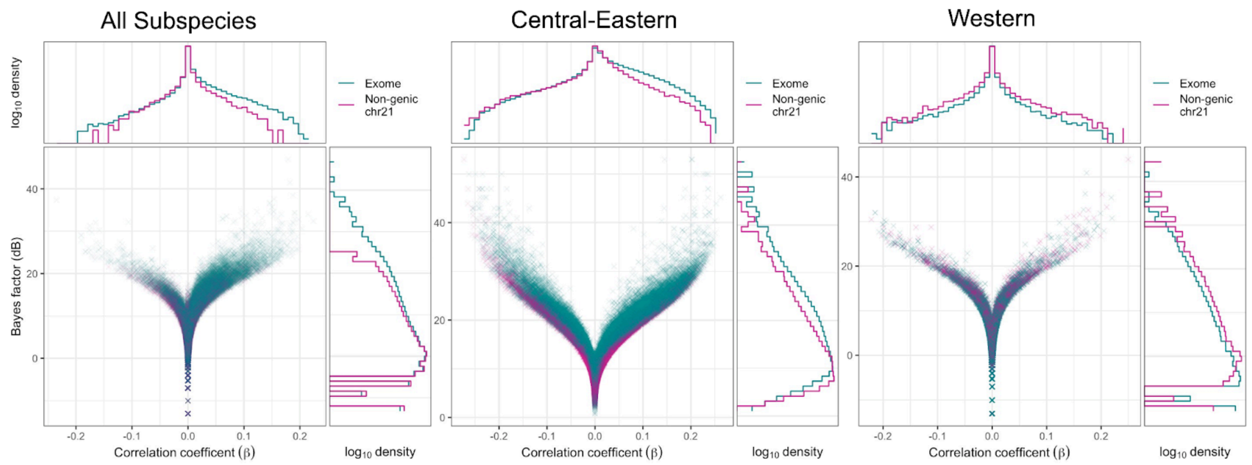


**Fig. S36. Volcano plots showing the BF and correlation coefficients estimated using the BayPass AUX model for each SNP in the exome (green) and non-genic-chr21 (purple) datasets.** The log density distributions of each axis are also plotted. As the significance of an association between a SNP and the covariable increases (higher BFs), the absolute value of the correlation coefficient is expected to increase resulting in a V-shape. The BF (y-axis) density plots show an excess of SNPs with very high BFs in the exome compared to non-genic-chr21 for *All* and *Central-Eastern* but not *Western*.

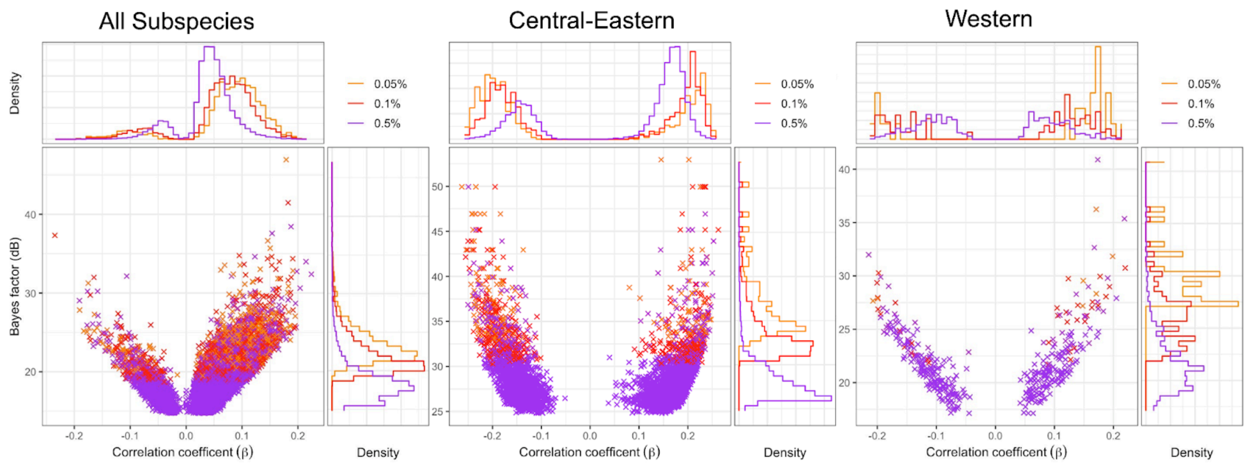


**Fig. S37. Volcano plots showing the BF and correlation coefficients at each FPR tail for each subspecies-dataset estimated using the BayPass AUX model.** The density distributions of each axis are also plotted. Note that these categories are nested (e.g. a SNP in the 0.1% tail is necessarily in the 0.5% tail but may not be in the 0.05% tail). In the volcano plot, more stringent tails are plotted on top of less stringent ones so points are coloured according to the most stringent tail for that SNP.

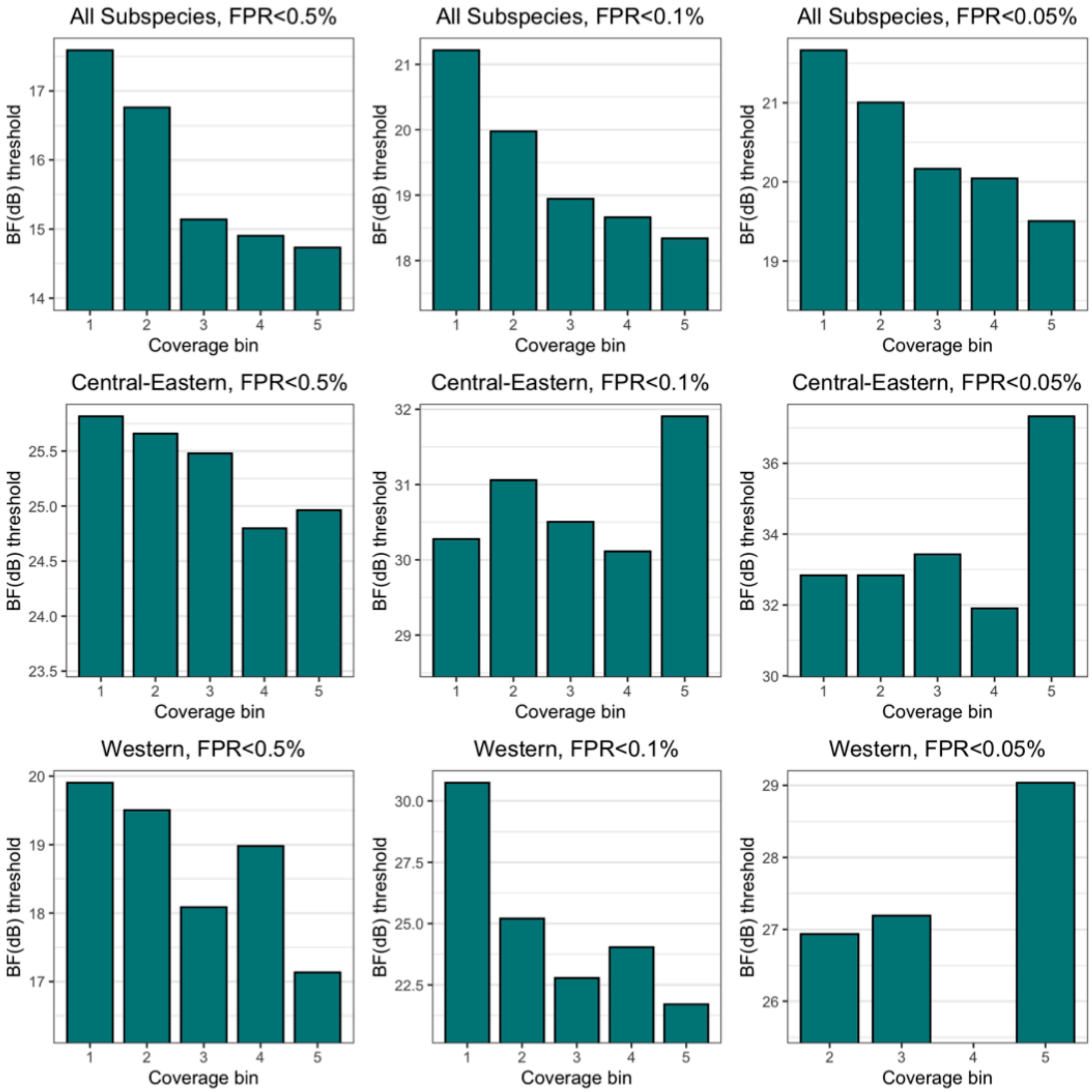


**Fig. S38. BF thresholds for each coverage bin, FPR tail and subspecies-dataset.**

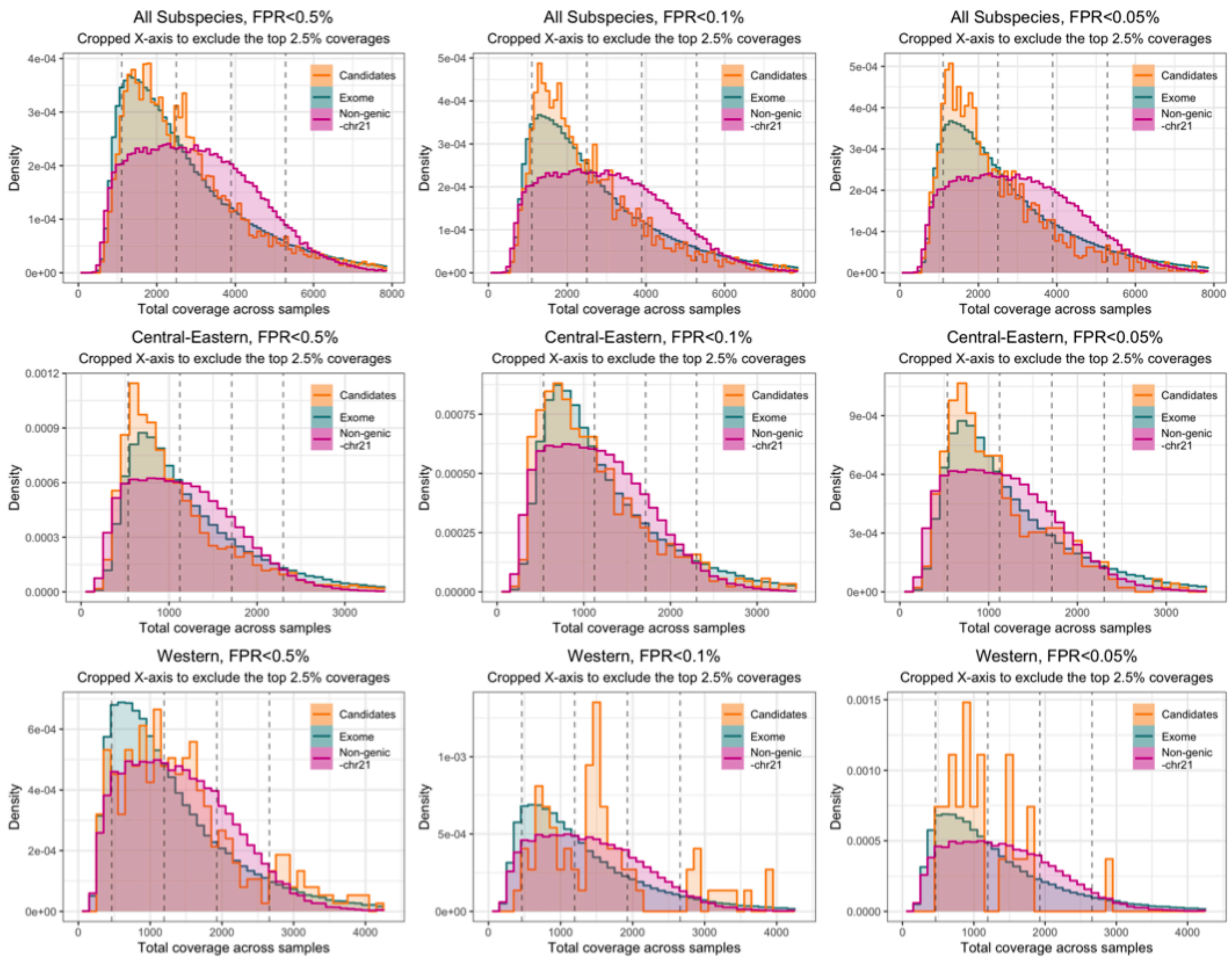


**Fig. S39. Distribution of total coverage across samples for the whole exome (green), non-genic-chr21 (purple) and the GEA candidates (orange).** Results shown for each FPR tail and each subspecies-dataset. The limits of the 5 coverage bins are shown as vertical dashed lines.

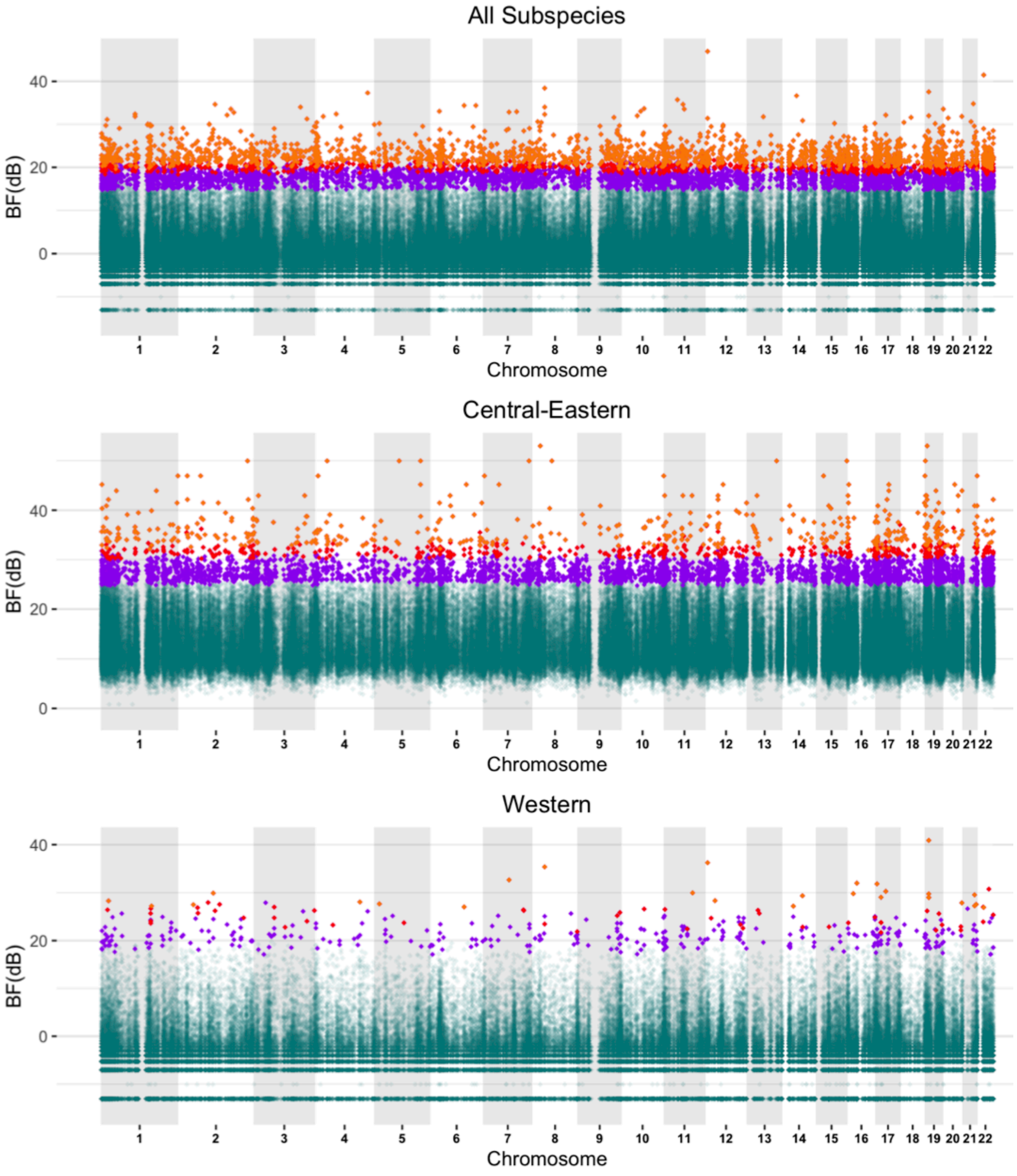


**Fig. S40. Manhattan plots showing the BF values from the GEA analysis for each dataset.** Points coloured purple, red and orange are in the 0.5%, 0.1% and 0.05% coverage corrected FPR tails respectively.

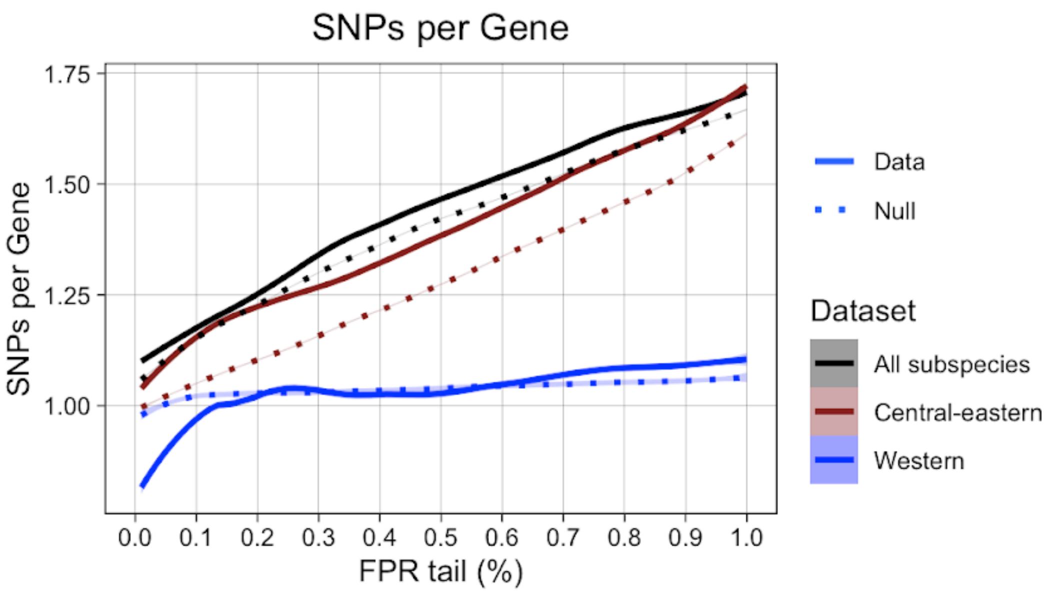


**Fig. S41. Number of SNPs per gene across FPR tails for the three subspecies-datasets analyses in the GEA.** Solid lines represent the real data sampled at 0.01% intervals and dotted lines represent null expectations generated by randomly sampling the same number of SNPs 50 times at each 0.01% interval. Shaded areas represent the 95% confidence intervals.

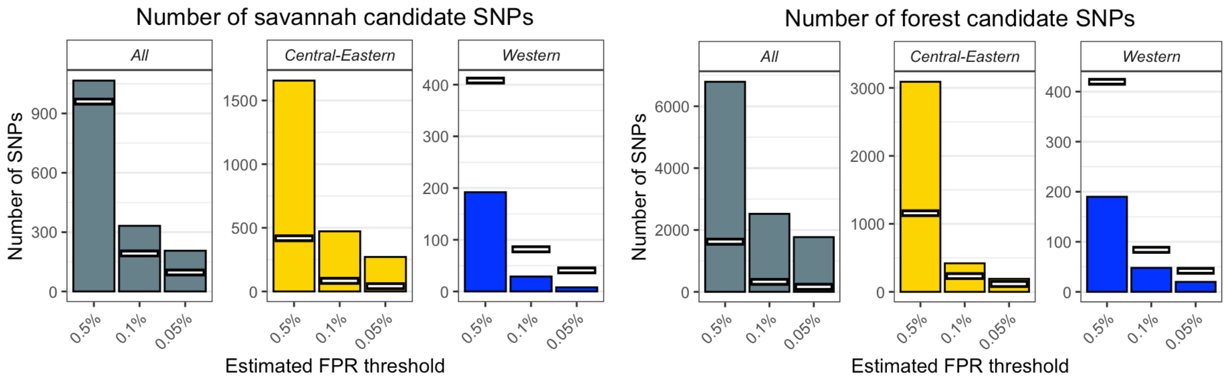


**Fig. S42. The number of candidate SNPs from the GEA (bars) compared to the null expectation (white lines) for savannah (left) and forest candidates (right).** Results shown for BF thresholds corresponding to estimated FPRs of 0.5%, 0.1% and 0.05%, for each subspecies-dataset tested Note that y-axis scales are not consistent across panels. This figure is the same as Fig. 4 only here savannah and forest candidates were selected separately; to select the savannah candidates, only exome and non-genic-chr21 SNPs with a negative correlation coefficient are considered and for forest only SNPs with positive correlation coefficients are used. The difference between the null expectations for forest and savannah candidate SNPs is explained by the skewed correlation coefficient distributions (Fig. S36). The fact that we find an excess for both forest and savannah candidates over null expectations in All and Central-Eastern using this method indicates that adaptation in either direction contributes to the overall excess of SNPs with high BF values in the exome. Note that the excess is much greater for the All forest candidates than for the savannah candidates indicating that adaptation to forests disproportionately contributes to the overall excess of SNPs highly correlated to habitat type in this subspecies-dataset. We note that using this method to select candidate SNPs results in a near-identical list of candidate SNPs as the main analysis. The only differences are due to the lower resolution of the FPR estimation resulting from fewer non-genic-chr21 SNPs use to generate each null distribution. This means that the candidates identified with this method are a subset of those identified in the main analysis.

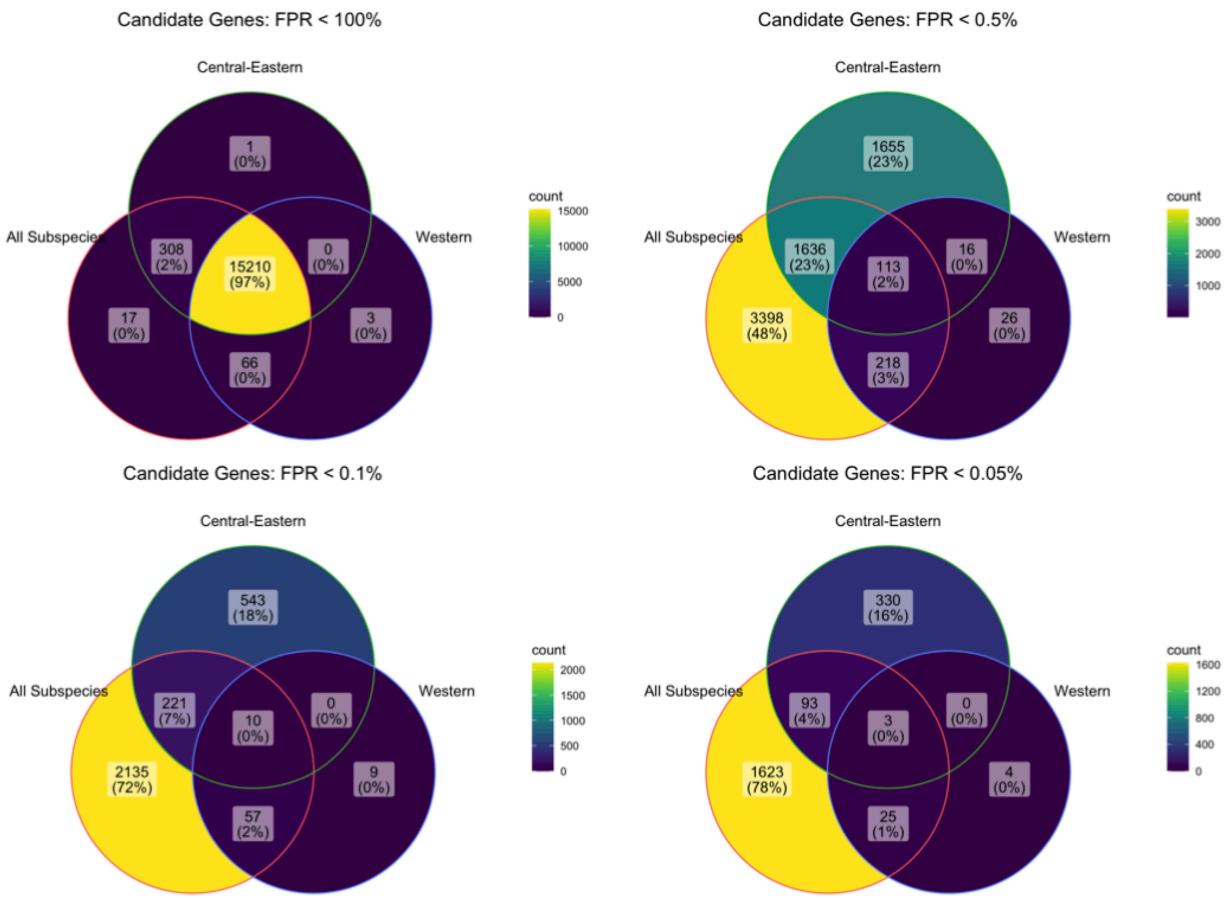


**Fig. S43. Overlap in genes between different datasets in all genes (i.e. FPR<100%) and the 0.5%, 0.1% and 0.05% FPR tails from the GEA analysis.**

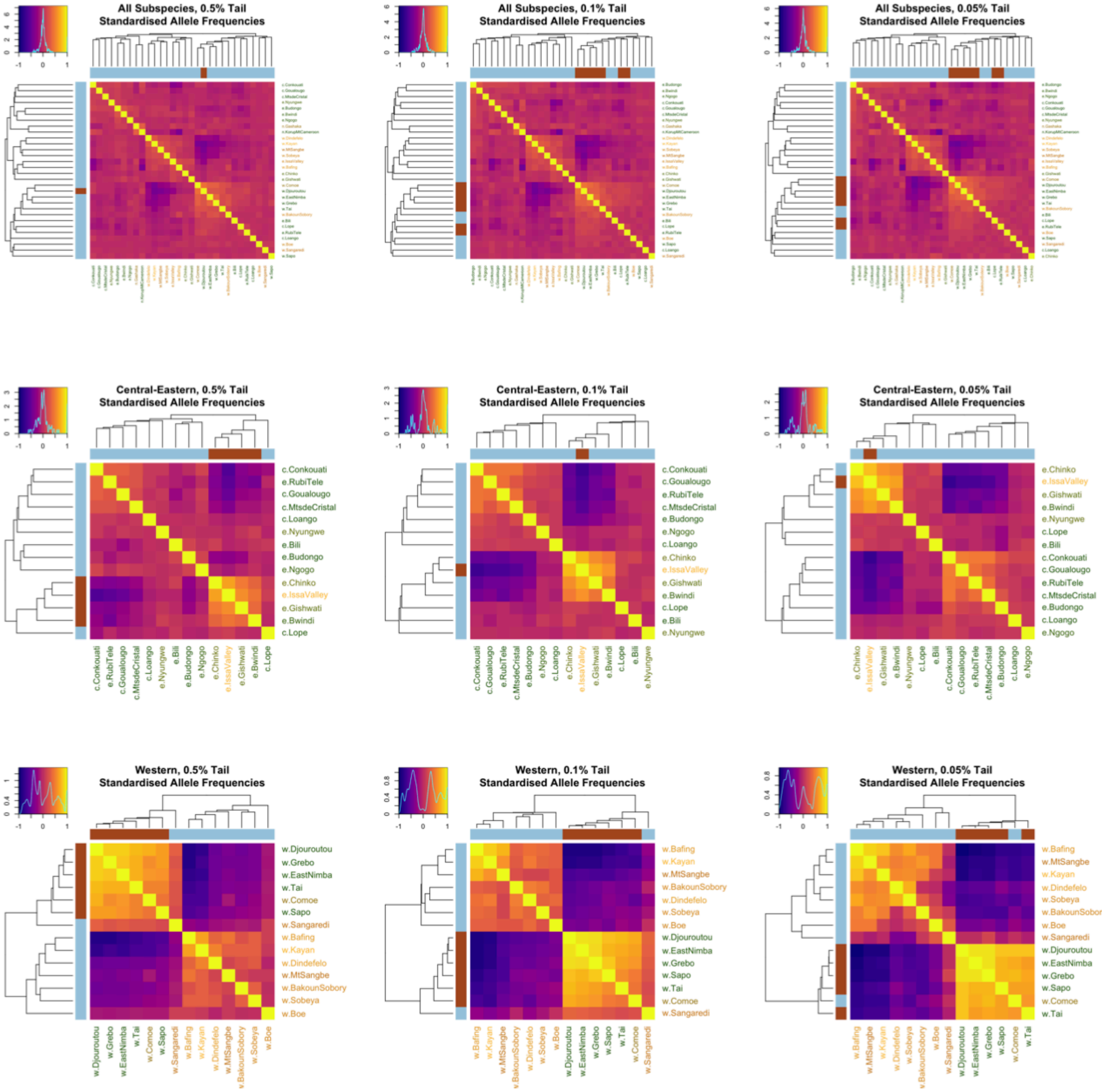


**Fig. S44. Correlation matrices calculated from the exome GEA candidate SNP standardised allele frequencies calculated under the BayPass core model.** Figures are shown for *All*, *Central-Eastern* and *Western* for the 0.5%, 0.1% and 0.05% GEA tails. Populations are ordered according to a dendrogram calculated using the average agglomeration method and the tips are coloured according to clusters assigned with k-medoids clustering. Population names are coloured according to forest-tree-percentage with greener colours representing higher values (more forest-like) and yellower colours representing lower values (more savannah-like). We can see structure driven by the habitat covariable.

**A**

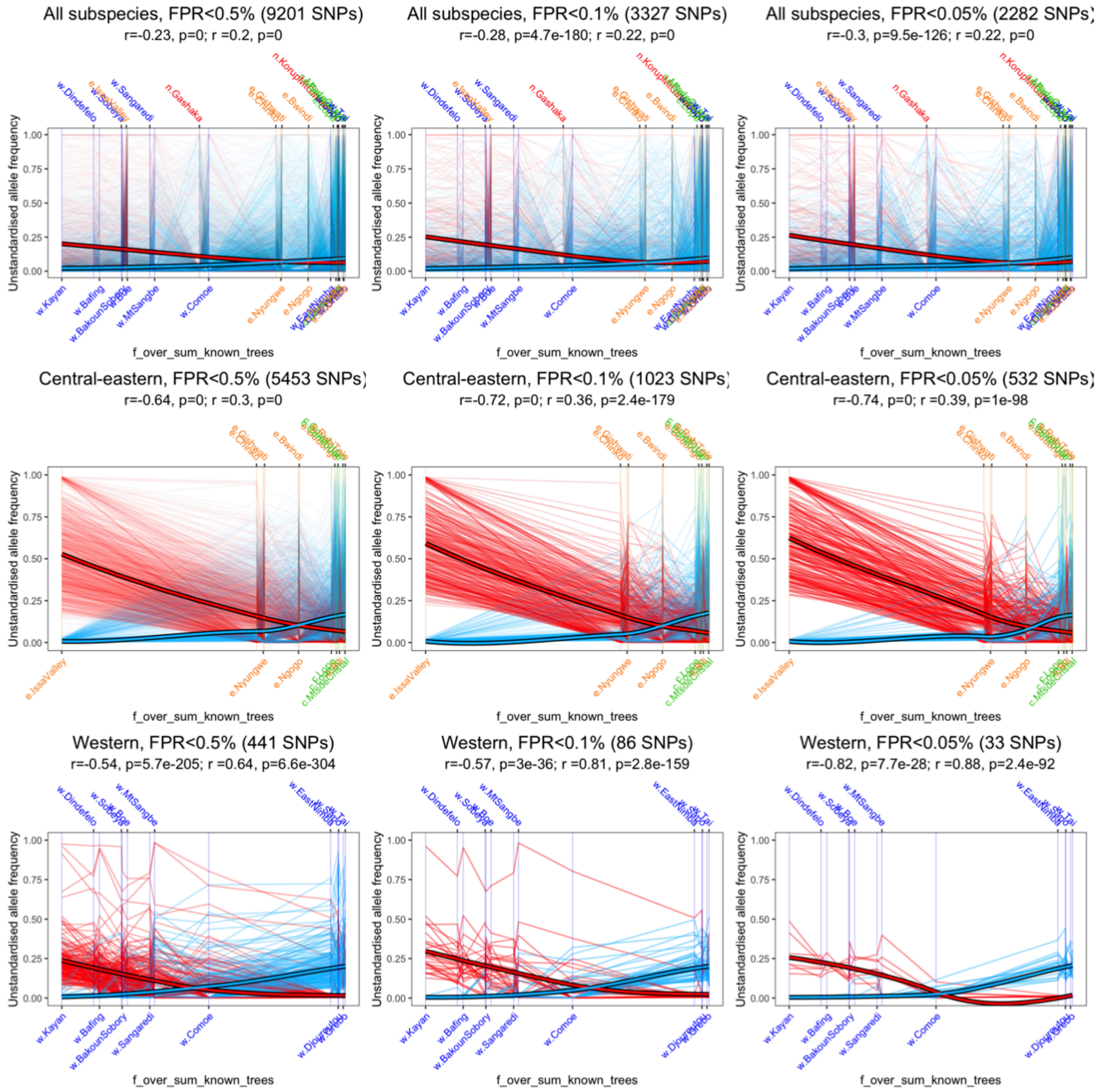
All subspecies, FPR<0.5% (9201 SNPs)
r=-0.23, p=0; r =0.2, p=0
All subspecies, FPR<0.1% (3327 SNPs)
r=-0.28, p=4.7e-180; r =0.22, p=0
All subspecies, FPR<0.05% (2282 SNPs)
r=-0.3, p=9.5e-126; r =0.22, p=0
Central-eastern, FPR<0.5% (5453 SNPs)
r=-0.64, p=0; r =0.3, p=0
Central-eastern, FPR<0.1% (1023 SNPs)
r=-0.72, p=0; r =0.36, p=2.4e-179
Central-eastern, FPR<0.05% (532 SNPs)
r=-0.74, p=0; r =0.39, p=1e-98
Western, FPR<0.5% (441 SNPs)
r=-0.54, p=5.7e-205; r =0.64, p=6.6e-304
Western, FPR<0.1% (86 SNPs)
r=-0.57, p=3e-36; r =0.81, p=2.8e-159
Western, FPR<0.05% (33 SNPs)
r=-0.82, p=7.7e-28; r =0.88, p=2.4e-92
Unstandardised allele frequency
f_over_sum_known_trees

**B**

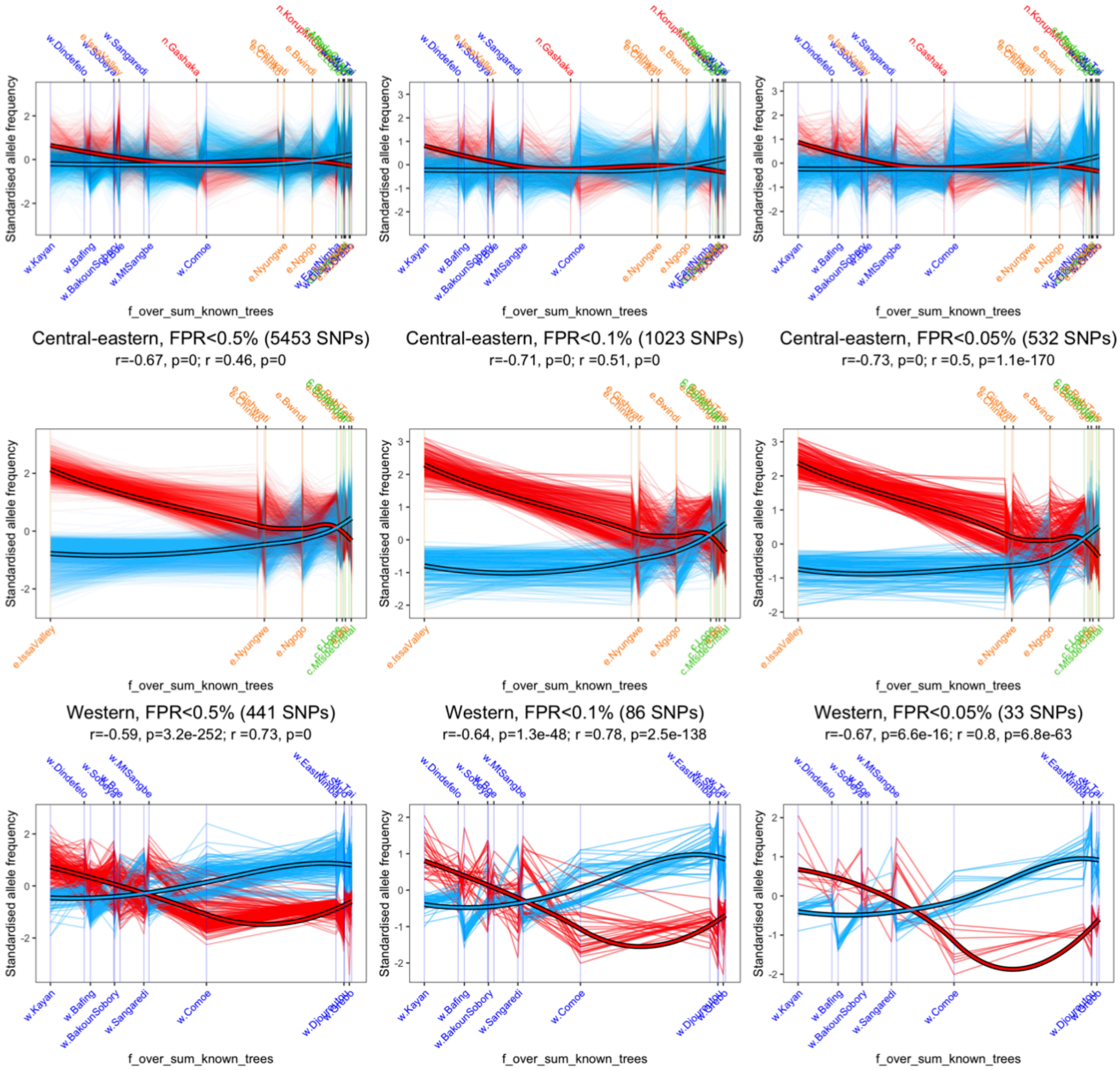


**Fig. S45. Candidate SNP unstandardised (A) and standardised (B) population allele frequencies (calculated under the core model) plotted against forest-tree-percentage.** Thin lines represent the estimated population allele frequencies for each candidate SNP, thick lines show the smoothed pattern of all candidate SNPs which are positively (i.e. forest candidates in blue) or negatively (i.e. savannah candidates in red) correlated with forest-tree-percentage. The position of each population is indicated on the x-axis and colour corresponds to the subspecies (green=central, orange=eastern, red=Nigeria-Cameroon, blue=western). Pearson correlation coefficients and p-values are given above the plot for savannah and forest candidates respectively.

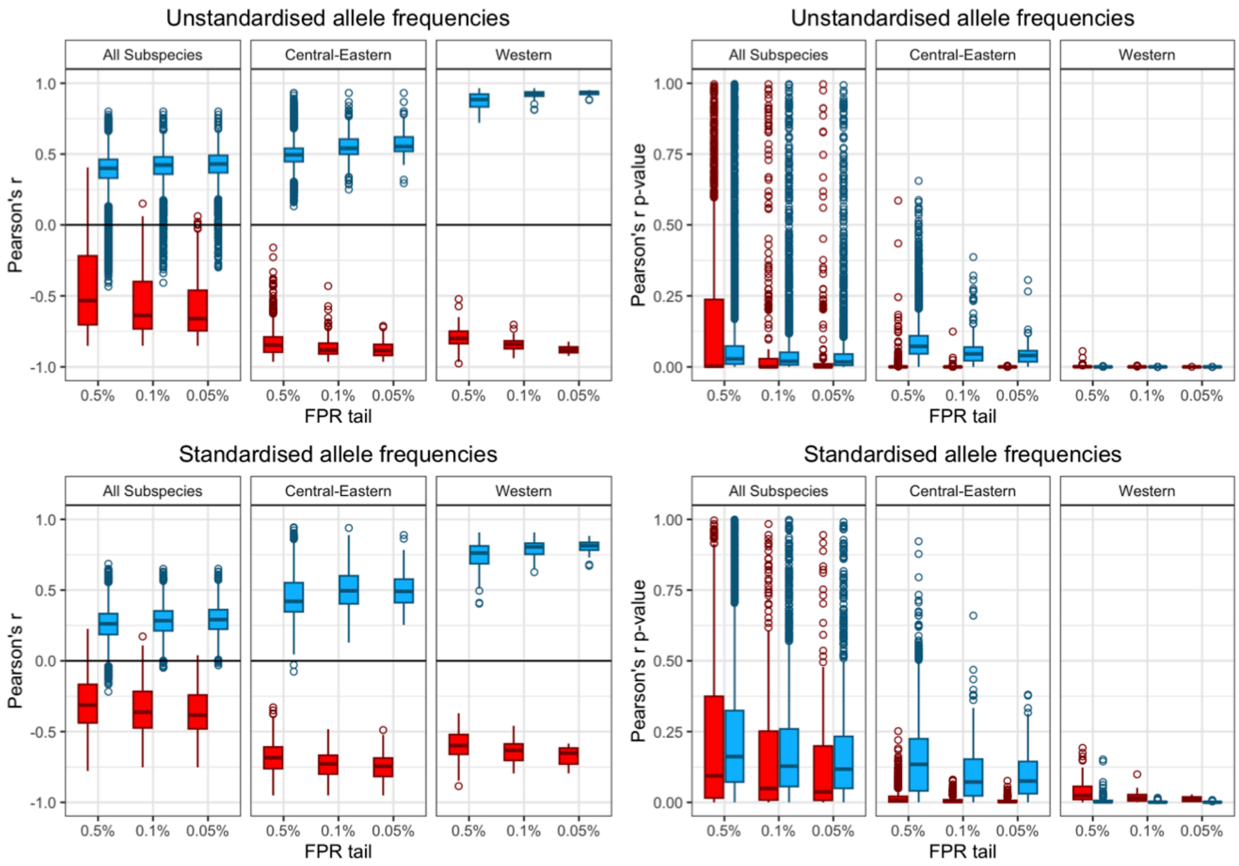


**Fig. S46. Pearson correlation coefficients (left) and p-values (right) for unstandardised (top) and standardised (bottom) allele frequencies (calculated under the core model) at each GEA candidate SNP with forest-tree-percentage.** Results are shown for the three subspecies-databases and three candidate tails. SNPs are separated into those where the derived allele is positively (i.e. forest candidates in blue) or negatively (i.e. savannah candidates in red) correlated with forest-tree-percentage in the BayPass GEA. More stringent tails generally result in higher absolute correlation coefficient values and lower p-values.

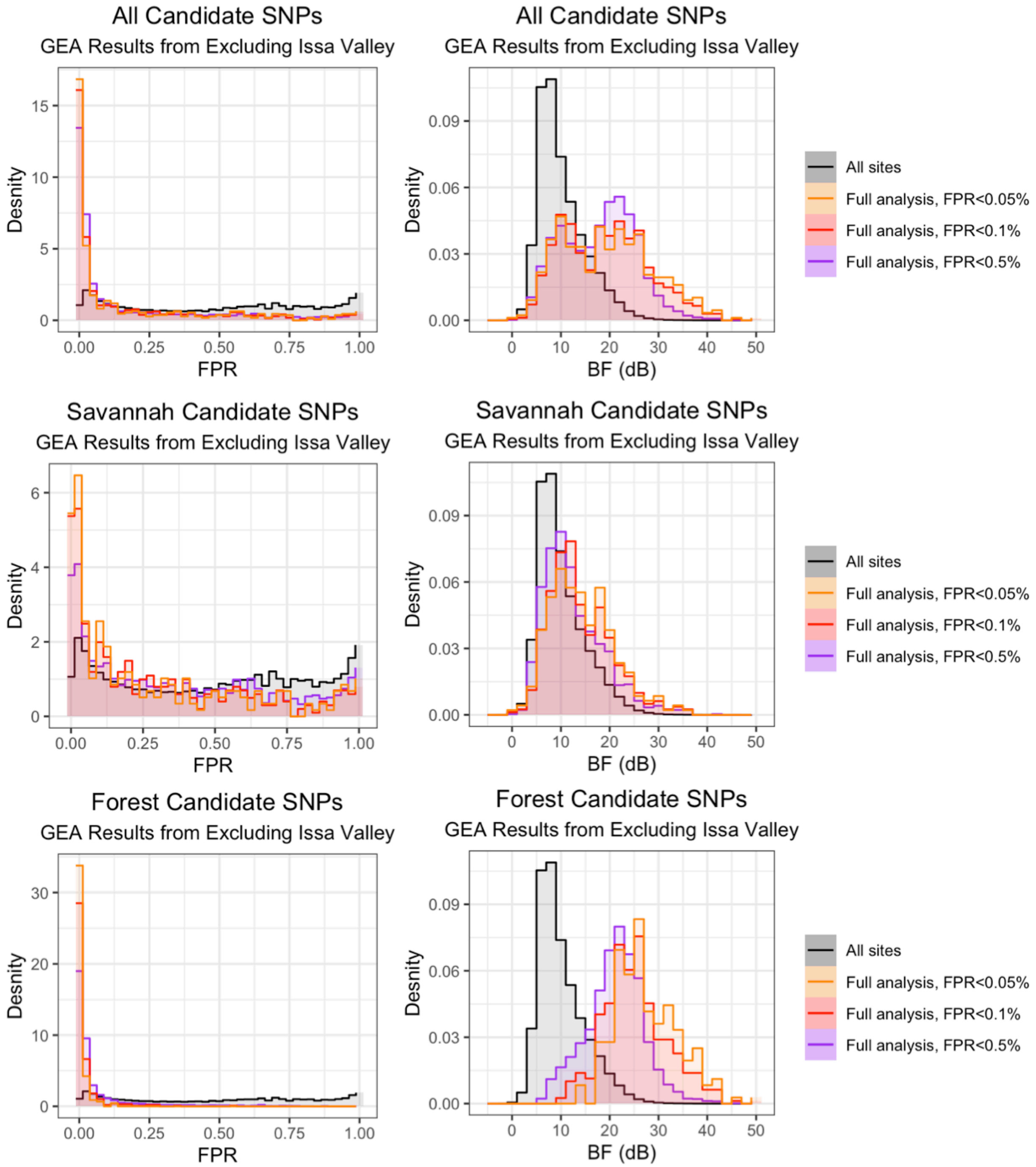


**Fig. S47. The distribution of FPRs (left) and BFs (right) from running BayPass on *Central-Eastern* with Issa Valley removed.** The distribution for all SNPs (black) and for SNPs identified as candidates (coloured by FPR threshold) in the full *Central-Eastern* analysis (i.e. with Issa Valley included) are shown. Top: all candidates. Middle: savannah candidates. Bottom: forest candidates. SNPs identified as candidates in the full analysis tend to have lower FPRs and higher BFs than the background.

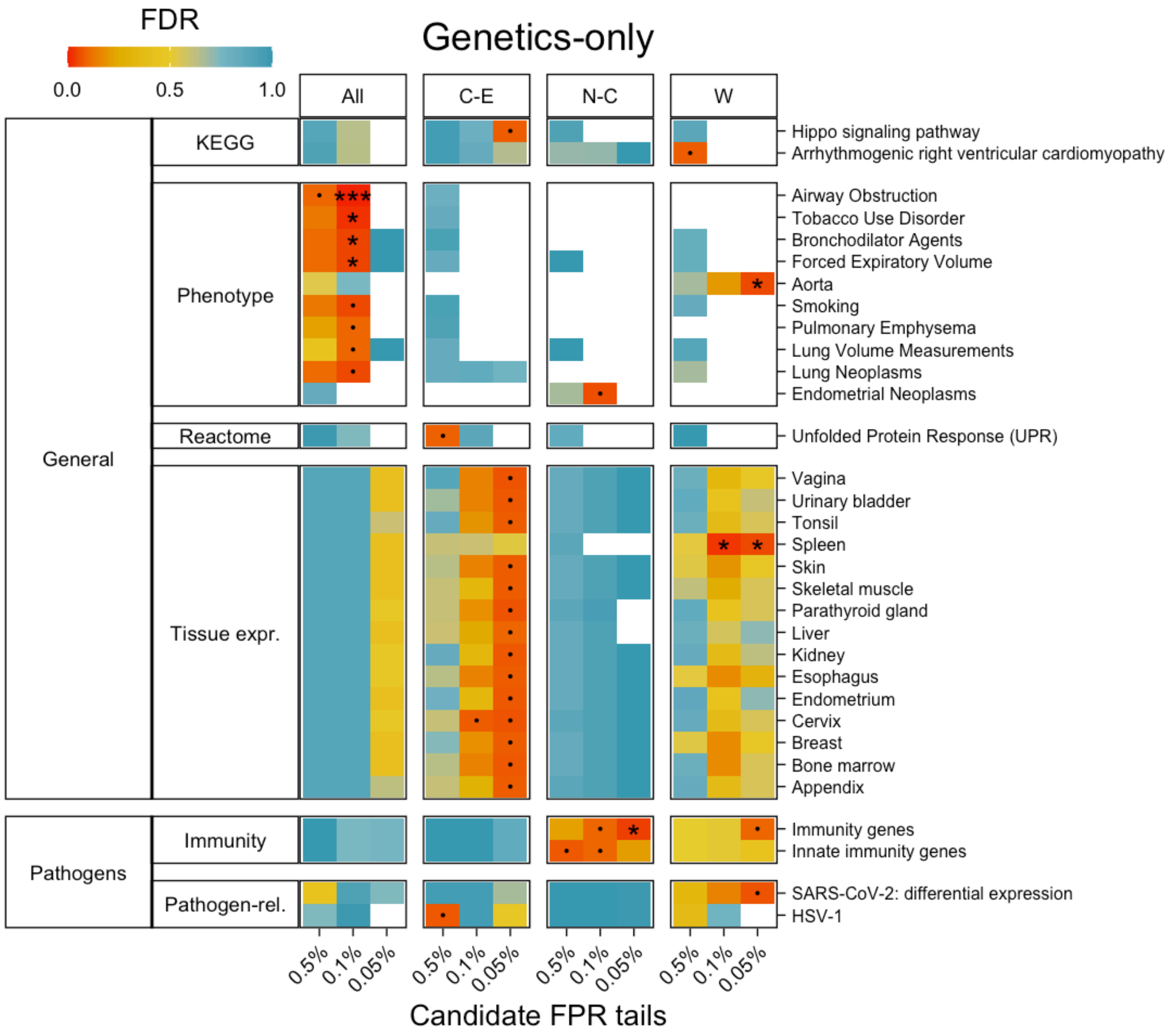


**Fig. S48. Gene set enrichment results for SNPs in the 0.5%, 0 .1% and 0.05% FPR tails of the genetics-only test ('.' FDR<0.1, '*' FDR<0.05, '**' FDR<0.01, '***' FDR<0.001).** Vertical panels indicate results from *All*, *Central-Eastern*, *Nigeria-Cameroon* and *Western* (from left to right). Horizontal panels show the categories that the gene sets belong to. Only gene sets with FDR<0.1 in any tail in any dataset are shown. Multiple testing correction was done within each gowinda run (i.e. each tail and gene set database such as KEGG, Phenotype etc.).

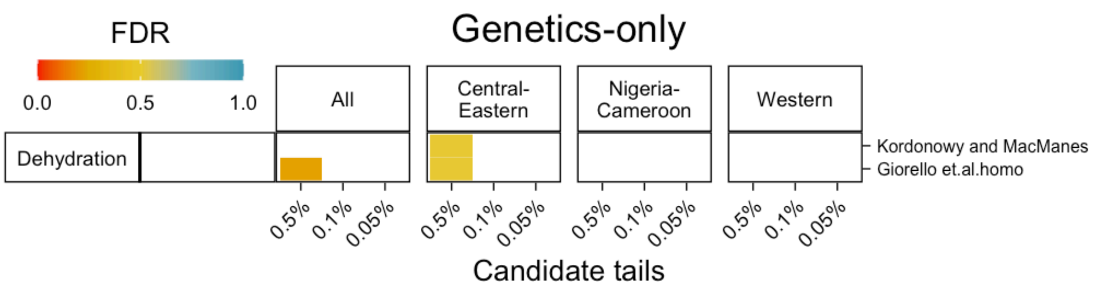


**Fig. S49. Enrichment of dehydration response genes in the genetics-only candidates.**

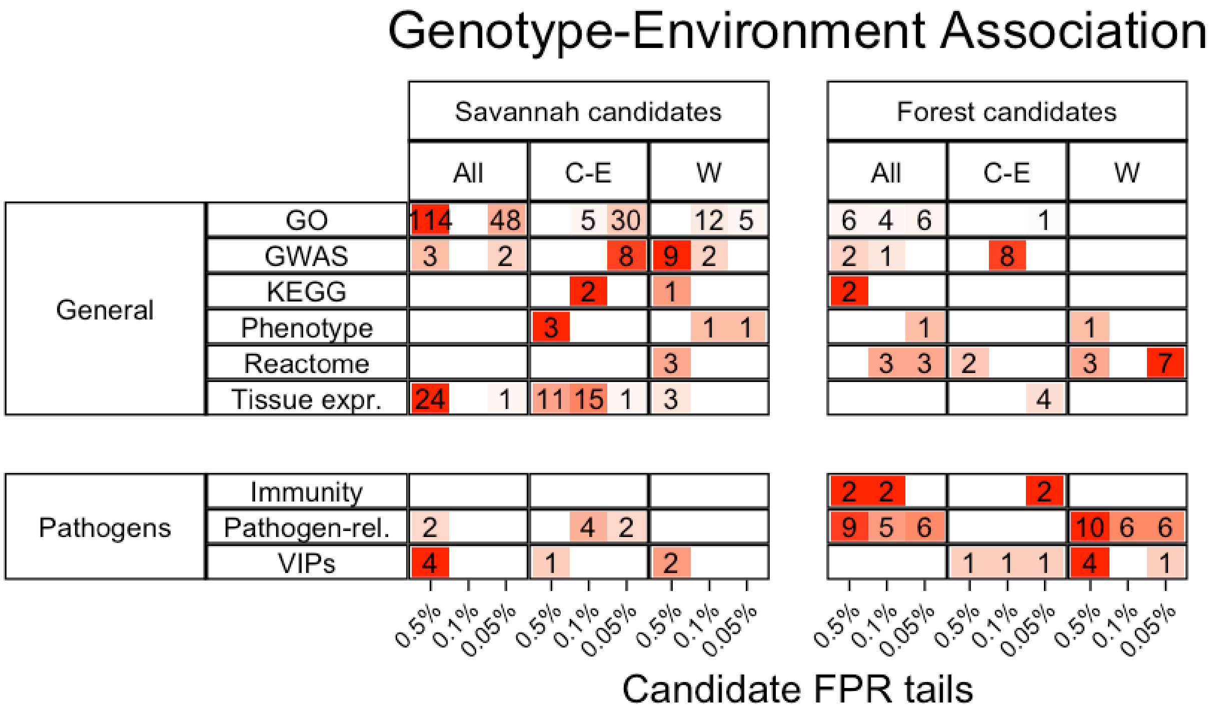


**Fig. S50. The number of gene sets with FDR < 0.5 from gene set enrichment results for GEA candidate SNPs.** Results are shown for 0.5%, 0.1% and 0.05% FPR tails for savannah and forest candidate SNPs. Vertical panels indicate results from each subspecies-dataset. Horizontal panels show the broad categories that the gene sets belong to. Multiple testing correction was done within each gene set enrichment analysis run (i.e., each tail and gene set database such as 'Pathogen-related', 'GWAS', 'Phenotype' etc.). Cells are coloured in a gradient from white (0) to red (the largest value per row). This is the same as Fig. 5A only with numbers indicated in the cells.

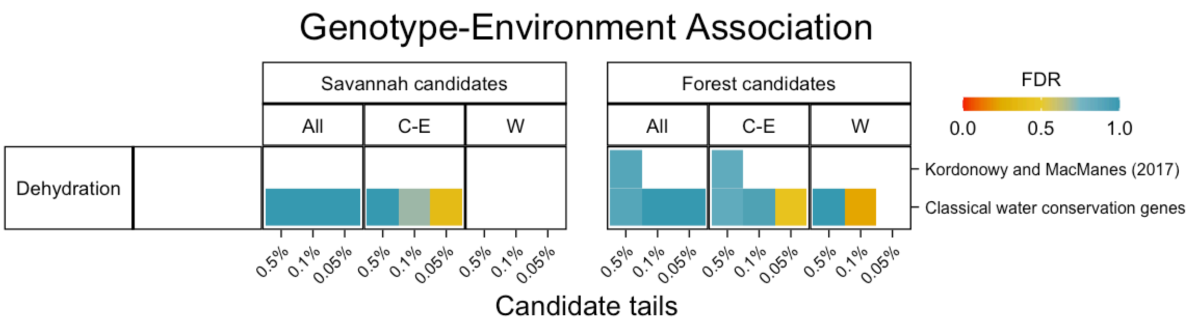


**Fig. S51. Enrichment of dehydration response genes in the GEA candidates.**

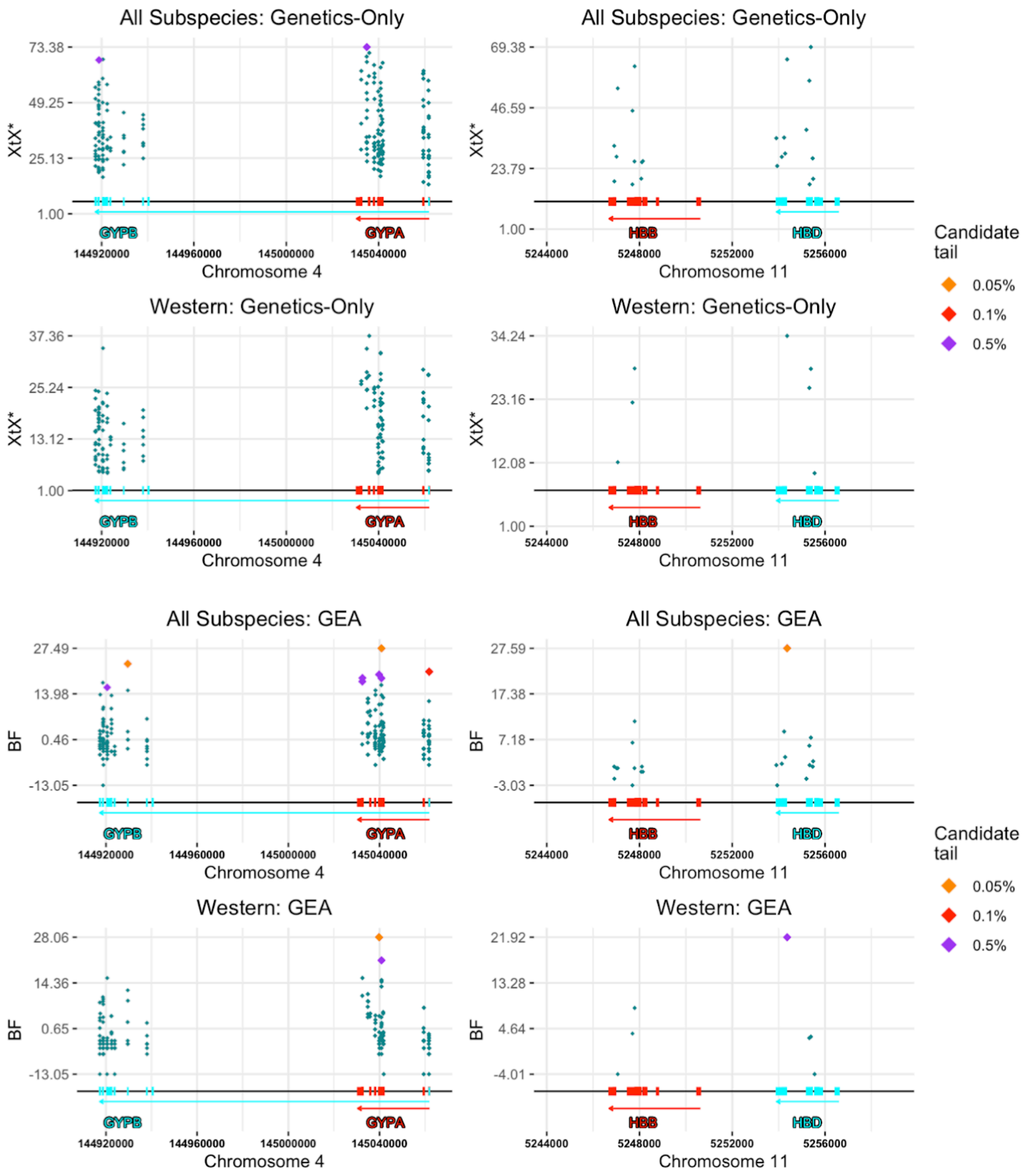


**Fig. S52. Manhattan plots showing the $X^tX^*$ and BF values from the genetics-only and GEA analyses respectively for *All* (top) and *Western* (bottom) datasets focusing on the *GYPA*/*GYPB* (left) and *HBB*/*HBD* (right) loci.** The position of exons are indicated as rectangles and arrows indicate total gene length and transcription direction (Ensembl/Havana merged) from the hg19 gtf annotation file downloaded from Ensembl (Methods). Annotations are coloured according to the gene within each panel. Points are coloured according to whether they are in the 0.5% (purple), 0.1% (red) or 0.05% (orange) FPR tails with all other SNPs in green.

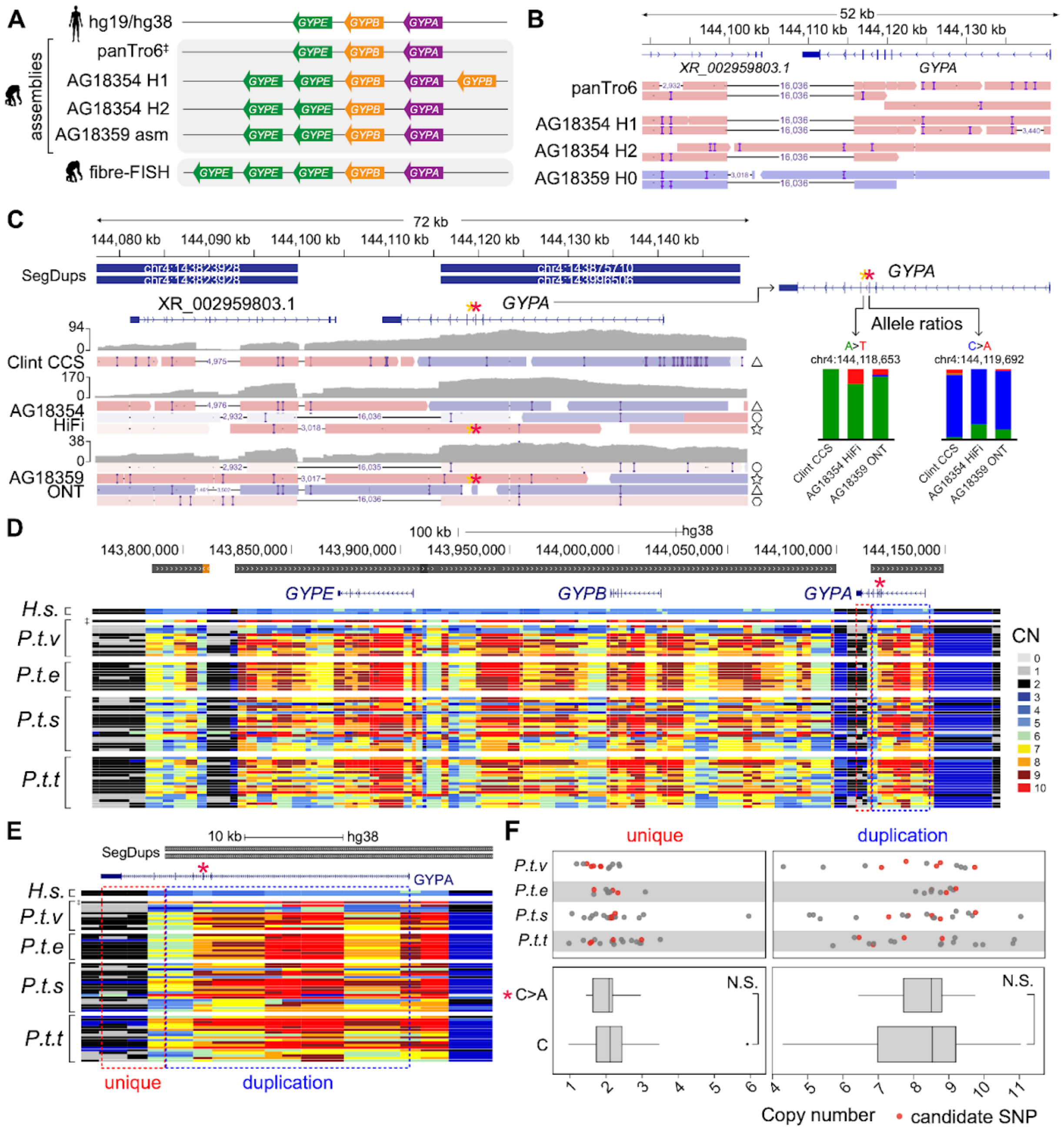


**Fig. S53. Glycophorin genes A-B-E copy-number analysis.** (**A**) Predicted gene annotations based on lift over of human genes to chimpanzee genome assemblies and previously published fibre-FISH data. (**B**) Mapping of chimpanzee genome assembly contigs at *GYPA* locus. (**C**) Direct mapping of long reads to *GYPA* locus displaying a representative set of reads carrying distinct molecular haplotypes annotated as triangle (non-delete haplotype not carrying the candidate SNPs), circle (deleted haplotype 1), hexagon (deleted haplotype 2), and star (non-deleted haplotype carrying the candidate SNPs [yellow asterisk: A>T at chr4:145,039,806; red asterisk: C>A at chr4:145,040,845 in hg19]). Grey shapes represent depth of coverage. Right insert shows allele ratios of A (green), C (blue), T (red) and G (ochre) nucleotides at candidate

SNP sites. (D) Gene-family copy number estimates in hg38 coordinates across chimpanzee subspecies for glycophorin genes A-B-E based on short-read depth, with copy numbers explained in legend (right). Symbol ‡ demarks Clint's assembly (panTro6) and short-read copy-number estimates. (E) Close up to *GYPA* copy number estimates highlighting unique and duplicated portions according to the segmental duplication track across chimpanzee subspecies. (F) Copy-number genotyping across unique and duplicated portions of *GYPA* obtained as the median copy-number across each region. Individuals carrying the C>A candidate SNP are highlighted in red. Significant differences were tested using Mann-Whitney U test. P.t.v.: *Pan troglodytes verus* (western). P.t.t: Pan *troglodytes troglodytes* (central). P.t.e.: *Pan troglodytes ellioti* (Nigeria-Cameroon). P.t.s.: *Pan troglodytes schweinfurthii* (eastern).

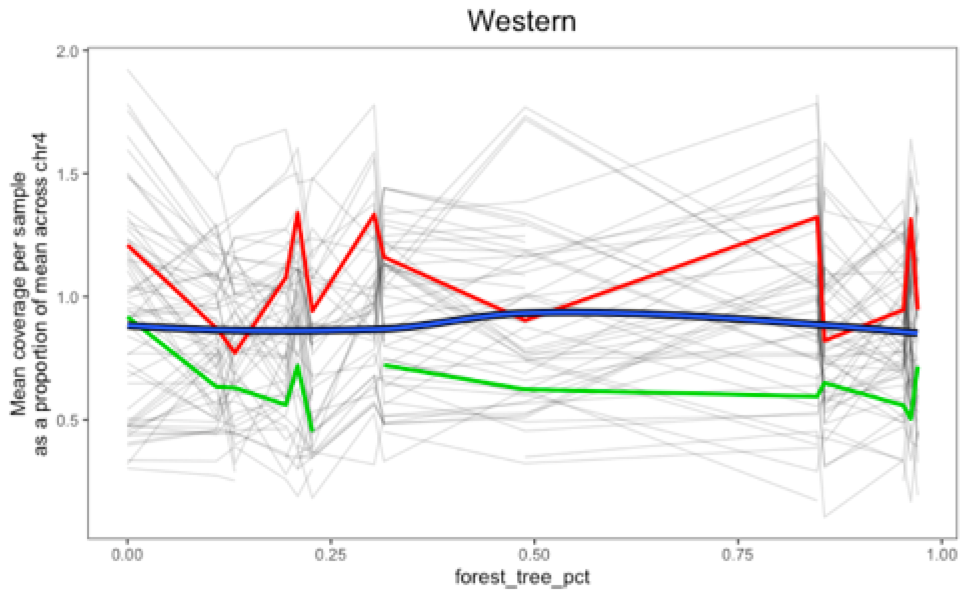


**Fig. S54. Mean read depth per sample for SNPs in the duplicated portion of *GYPA* corrected for the mean across the whole of chr4 per population in the exome data.** Thin lines represent individual SNPs, the green line represents the candidate SNP at chr4:145039806 (hg19) and the red line represents the candidate SNP at chr4:145040845 (hg19). The thick blue line represents the smoothed pattern using LOESS. Some of the per SNP lines are fragmented because they did not pass quality filters in all populations. We see no evidence of particularly high read depth or correlation between forest-tree-percentage and read depth at this locus in the exome data, suggesting observed SNP associations are not due to artefacts due to structural variation at this locus.

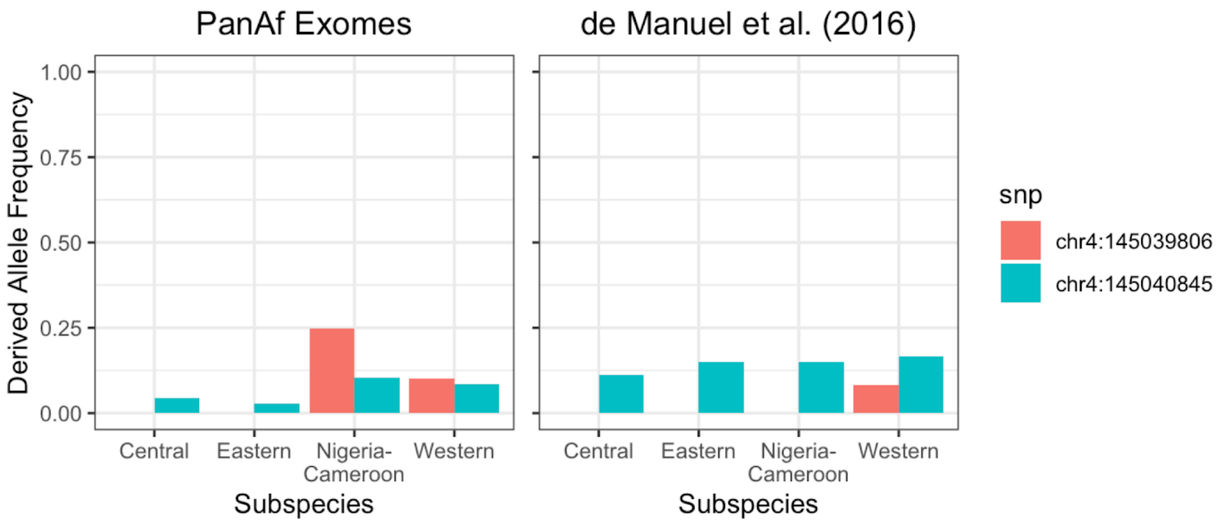


**Fig. S55. The allele frequency of the two *GYPA* candidate SNPs in the PanAf exome data (left) and the high-coverage short-read data (*62*) (right).**

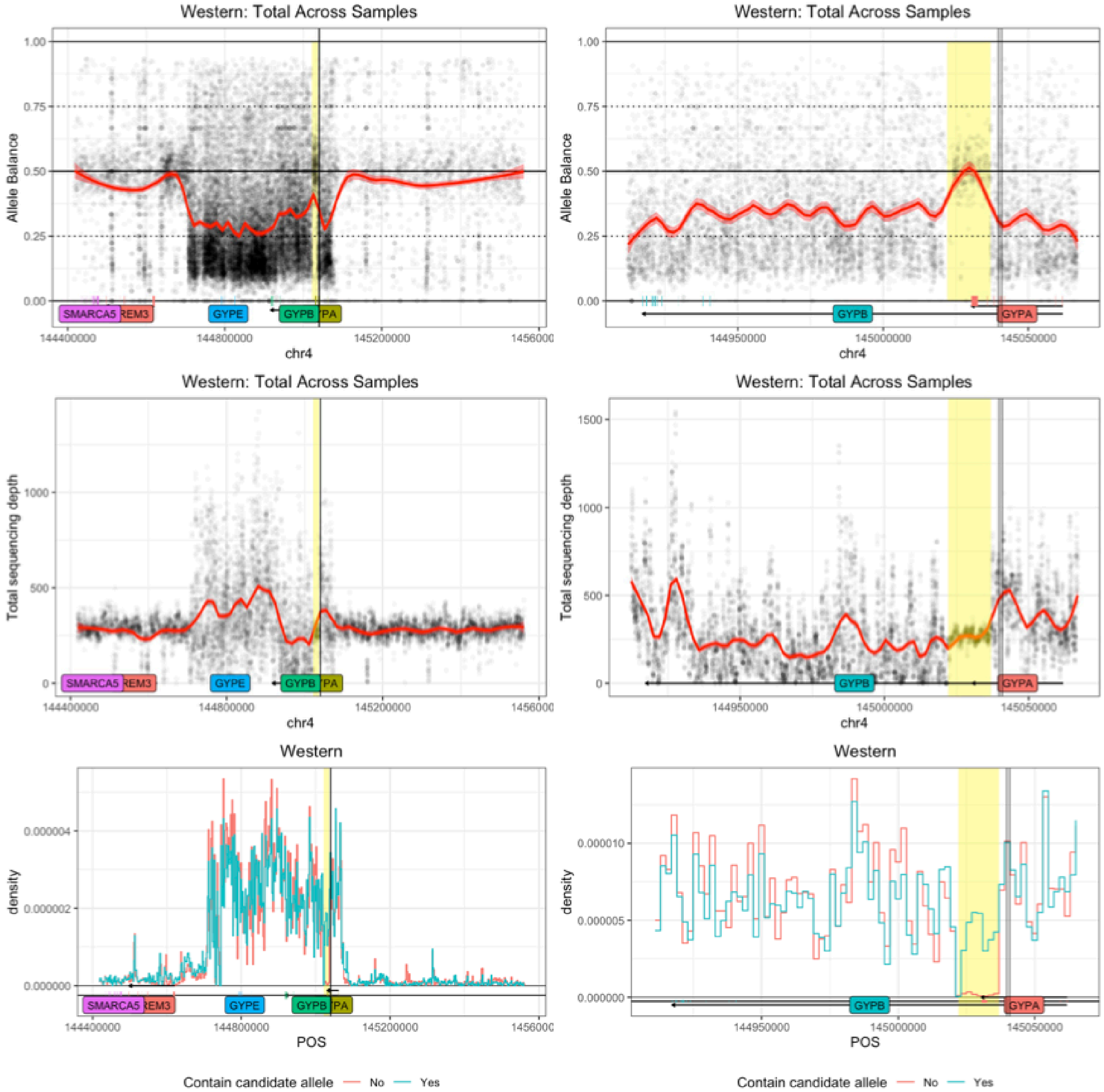


**Fig. S56. The allele balance (top), mean sequencing depth per sample (middle) and density of heterozygous sites (bottom) in the high-coverage short-read data from the western high-coverage short-read samples** (*62*) **at the *GYPA*/*GYPB* region ±500kb (left) or ±5kb (right).** Vertical black lines indicate the location of the *GYPA* candidate SNPs. 'Carriers' are samples which have at least one copy of the derived allele at the candidate SNP chr4:145040845. The skewed allelic balance, high coverage and high density of heterozygous sites across the region appear to indicate the presence of duplications. The yellow region highlights a 15kb long region which roughly coincides with an allelic balance of 50%, lower coverage and lower density of heterozygous sites than the *GYPA*/*GYPB* region as a whole suggesting that this region may not be duplicated.

**Table S1. Number of populations in each subspecies-dataset and number of SNPs which have allele count data for all populations in the dataset or at least 70% of populations.** Note that the number of SNPs in each subspecies-dataset is strongly influenced by factors such as coverage or number of populations (e.g., a relatively large number of SNPs pass the filter of no missing data in *Nigeria-Cameroon* because it has only two populations).

| | ***All*** | | ***Central-Eastern*** | | ***Nigeria-Cameroon*** | | ***Western*** | |
|---|---|---|---|---|---|---|---|---|
| Populations | 30 | | 14 | | 2 | | 14 | |
| **Populations with data** | **100%** | **≥70%** | **100%** | **≥70%** | **100%** | **≥70%** | **100%** | **≥70%** |
| Exome SNPs | 61,967 | 521,015 | 47,202 | 314,934 | 108,382 | 108,382 | 88,630 | 175,266 |
| Non-geneic-chr21 SNPs | 72,254 | 172,875 | 59,410 | 124,407 | 45,339 | 45,339 | 34,391 | 48,829 |

**Table S2. Gene set enrichment results for GEA candidate SNPs showing the number of candidate genes and FDR for all results with FDR < 0.1.** These numbers correspond to the results reported in Fig. 5.

| Direction | Category | Dataset | Subspecies | Gene set | Tail | Number of candidate genes | FDR |
|---|---|---|---|---|---|---|---|
| Savannah candidates | General | Gene ontology | All | Negative regulation of cellular macromolecule biosynthetic process | 0.50% | 96 | 0.023 |
| Savannah candidates | General | Gene ontology | All | Negative regulation of nitrogen compound metabolic process | 0.50% | 108 | 0.023 |
| Savannah candidates | General | Gene ontology | C-E | Positive regulation of biomineral tissue development | 0.05% | 5 | 0.063 |
| Savannah candidates | General | Gene ontology | C-E | Positive regulation of bone mineralization | 0.05% | 5 | 0.063 |
| Savannah candidates | Pathogens | Pathogen-related | C-E | GWAS: AIDS progression | 0.10% | 2 | 0.096 |
| Savannah candidates | Pathogens | Pathogen-related | C-E | GWAS: AIDS progression | 0.05% | 2 | 0.036 |
| Forest candidates | General | Reactome | All | IRE1alpha activates chaperones | 0.10% | 18 | 0.071 |
| Forest candidates | General | Reactome | All | IRE1alpha activates chaperones | 0.05% | 15 | 0.076 |
| Forest candidates | Pathogens | Pathogen-related | W | GWAS: HIV-1 susceptibility | 0.05% | 1 | 0.079 |
| Forest candidates | Pathogens | Pathogen-related | W | Malaria (conserved genes) | 0.50% | 8 | 0.050 |
| Forest candidates | Pathogens | Pathogen-related | W | Malaria (erythrocyte genes) | 0.50% | 4 | 0.034 |
| Forest candidates | Pathogens | Pathogen-related | W | Malaria (erythrocyte genes) | 0.10% | 3 | 0.020 |
| Forest candidates | Pathogens | Pathogen-related | W | Malaria (erythrocyte genes) | 0.05% | 2 | 0.033 |
| Forest candidates | Pathogens | Pathogen-related | W | Malaria | 0.50% | 5 | 0.009 |
| Forest candidates | Pathogens | Pathogen-related | W | Malaria | 0.10% | 2 | 0.091 |
| Forest candidates | Pathogens | Pathogen-related | W | Malaria | 0.05% | 2 | 0.033 |
| Forest candidates | Pathogens | Pathogen-related | W | SARS-CoV-2: GWAS | 0.50% | 2 | 0.063 |
| Forest candidates | Pathogens | VIPs | C-E | Rotavirus | 0.05% | 2 | 0.099 |

**Table S3. Gene set enrichment results (FDR) for GEA forest candidate SNPs in *Western* for pathogen response gene sets from running with every combination of removing *HBB* or *HBD* and *GYPA* or *GYPB*.** Within each gene combination category, gene sets are in descending order according to their FDR in the 0.5% tail and ties are resolved according to the FDRs for increasingly stringent tails. Categories containing no *Western* candidates are not displayed.

| Genes Removed | Gene Set | 0.50% | 0.10% | 0.05% |
|---|---|---|---|---|
| **HBB-GYPB** | **Malaria (conserved genes) (Ebel et al., 2017)** | **0.078685** | **0.15805** | **0.3497925** |
| **HBB-GYPB** | **Malaria (Daub et al., 2013)** | **0.078685** | **0.217846** | **0.18796** |
| HBB-GYPB | SARS-CoV-2: GWAS | 0.088506667 | | |
| **HBB-GYPB** | **Malaria (erythrocyte genes) (Ebel et al., 2021)** | **0.24366** | **0.06068** | **0.18796** |
| HBB-GYPB | GWAS: HIV-1 susceptibility | 0.4447375 | 0.15805 | 0.18796 |
| HBB-GYPB | HSV-1 | 0.4447375 | 0.56326625 | 0.437972 |
| HBB-GYPB | Influenza (Rogers et al., 2019) | 0.4447375 | 0.56326625 | 0.499005714 |
| HBB-GYPB | SARS-CoV-2: differential RNA expression | 0.4447375 | | |
| HBB-GYPB | GWAS: HIV-1 control | 0.452946667 | 0.217846 | |
| HBB-GYPB | SIV response | 0.466637 | 0.408291667 | 0.459905 |
| HBB-GYPB | SARS-CoV-2: host-protein interaction | 0.771970909 | | |
| **HBD-GYPB** | **Malaria (erythrocyte genes) (Ebel et al., 2021)** | **0.052986667** | **0.06088** | **0.188163333** |
| **HBD-GYPB** | **Malaria (conserved genes) (Ebel et al., 2017)** | **0.052986667** | **0.157013333** | **0.416055** |
| **HBD-GYPB** | **Malaria (Daub et al., 2013)** | **0.052986667** | **0.217254** | **0.188163333** |
| HBD-GYPB | SARS-CoV-2: GWAS | 0.0617475 | | |
| HBD-GYPB | GWAS: HIV-1 susceptibility | 0.4445425 | 0.157013333 | 0.188163333 |
| HBD-GYPB | HSV-1 | 0.4445425 | 0.56373875 | 0.421611667 |
| HBD-GYPB | Influenza (Rogers et al., 2019) | 0.4445425 | 0.56373875 | 0.49936 |
| HBD-GYPB | SARS-CoV-2: differential RNA expression | 0.4445425 | | |
| HBD-GYPB | GWAS: HIV-1 control | 0.447646 | 0.217254 | |
| HBD-GYPB | SIV response | 0.447646 | 0.408323333 | 0.421611667 |
| HBD-GYPB | SARS-CoV-2: host-protein interaction | 0.771597273 | | |
| **HBB-GYPA** | **Malaria (conserved genes) (Ebel et al., 2017)** | **0.08298** | **0.186293333** | **0.41852** |
| **HBB-GYPA** | **Malaria (Daub et al., 2013)** | **0.08298** | **0.240376** | **0.16957** |
| HBB-GYPA | SARS-CoV-2: GWAS | 0.09933 | | |
| **HBB-GYPA** | **Malaria (erythrocyte genes) (Ebel et al., 2021)** | **0.30221** | **0.08485** | **0.16957** |
| HBB-GYPA | GWAS: HIV-1 susceptibility | 0.456915556 | 0.186293333 | 0.16957 |
| HBB-GYPA | GWAS: HIV-1 control | 0.456915556 | 0.240376 | |
| HBB-GYPA | HSV-1 | 0.456915556 | 0.56736125 | 0.4403 |
| HBB-GYPA | Influenza (Rogers et al., 2019) | 0.456915556 | 0.56736125 | 0.537411429 |
| HBB-GYPA | SARS-CoV-2: differential RNA expression | 0.456915556 | | |
| HBB-GYPA | SIV response | 0.59879 | 0.560921667 | 0.537411429 |
| HBB-GYPA | SARS-CoV-2: host-protein interaction | 0.78399 | | |
| **HBD-GYPA** | **Malaria (erythrocyte genes) (Ebel et al., 2021)** | **0.06887** | **0.08583** | **0.192683333** |
| **HBD-GYPA** | **Malaria (conserved genes) (Ebel et al., 2017)** | **0.06887** | **0.176183333** | **0.4192525** |
| **HBD-GYPA** | **Malaria (Daub et al., 2013)** | **0.06887** | **0.240662** | **0.192683333** |
| HBD-GYPA | SARS-CoV-2: GWAS | 0.06887 | | |
| HBD-GYPA | GWAS: HIV-1 susceptibility | 0.45676 | 0.176183333 | 0.192683333 |
| HBD-GYPA | GWAS: HIV-1 control | 0.45676 | 0.2297475 | |
| HBD-GYPA | HSV-1 | 0.45676 | 0.5668975 | 0.441556 |
| HBD-GYPA | Influenza (Rogers et al., 2019) | 0.45676 | 0.5668975 | 0.538551429 |
| HBD-GYPA | SARS-CoV-2: differential RNA expression | 0.45676 | | |
| HBD-GYPA | SIV response | 0.589864 | 0.561503333 | 0.538551429 |
| HBD-GYPA | SARS-CoV-2: host-protein interaction | 0.783401818 | | |

**Table S4. Gene set enrichment results (FDR) for GEA forest candidate SNPs in the western subspecies for pathogen response gene sets after editing the annotation file so *GYPB* does not overlap with *GYPA*.** Within each gene combination category, gene sets are in descending order according to their FDR in the 0.5% tail and ties are resolved according to the FDRs for increasingly stringent tails. Categories containing no *Western* candidates are not displayed.

| Gene Set | 0.50% | 0.10% | 0.05% |
|---|---|---|---|
| **Malaria (Daub et al., 2013)** | **0.02675** | **0.241256** | **0.194206667** |
| **Malaria (conserved genes) (Ebel et al., 2017)** | **0.07495** | **0.15857** | **0.356205** |
| **Malaria (erythrocyte genes) (Ebel et al., 2021)** | **0.0775225** | **0.08409** | **0.194206667** |
| SARS-CoV-2: GWAS | 0.0775225 | | |
| GWAS: HIV-1 susceptibility | 0.413656667 | 0.15857 | 0.194206667 |
| SIV response | 0.413656667 | 0.41426 | 0.463906667 |
| HSV-1 | 0.413656667 | 0.56782 | 0.442466 |
| Influenza (Rogers et al., 2019) | 0.413656667 | 0.56782 | 0.501962857 |
| SARS-CoV-2: differential RNA expression | 0.413656667 | | |
| GWAS: HIV-1 control | 0.416786 | 0.241256 | |
| SARS-CoV-2: host-protein interaction | 0.78204 | | |
